# Evolution of novel host-microbiota coupling underlies adaptation to nutrient-scarce environments

**DOI:** 10.64898/2026.10.10.758241

**Authors:** Motoko Iwashita, Crystal Valdez, Alan G. Hudson, Xavier Barragan Gonzalez, Jaimee Kato, Emma Doy, Amity Tran, Lillian J. Tuttle Raz, Rebecca D. Prescott, Kaitlin J. Coyle, Aldo Carmona Baez, Natalie Roberts, Reade B. Roberts, Farzana Nasrin, Masato Yoshizawa

## Abstract

Animals colonizing food-limited environments need to overcome growth constraints by adapting to low, inconsistent nutritional inputs. Comparative studies of surface and cave populations of the Mexican tetra fish species *Astyanax mexicanus* provide a powerful system to explore evolution to food scarcity. In nutrient-scarce caves, evolved traits such as eye loss and increased fat storage have been shown to conserve energy in cavefish^1–3^. However, the role of microbiota-mediated metabolism in sustaining growth and survival under food limitation remains poorly understood. Here, we demonstrate that surface and cave populations distinctly modulate their gut microbiota, with cavefish recruiting microbes whose predicted metabolites specifically promote growth under nutrient-limited conditions. Using microbiota data from three distinct dietary regimes and a fasting treatment, integrated topological data analysis^4^ and sparse modeling identify sugar and fatty acid degradation pathways coupled with growth in cavefish. Notably, the metabolites of these pathways are beneficial only for cavefish growth, and not for surface fish.

Fasting in cavefish led to the recruitment of gut microbes capable of metabolizing aromatic compounds inaccessible to the host, whereas it recruited few microbial associates in surface fish. Our findings indicate that cavefish evolved novel host-microbiota coupling to sustain growth and resilience in their nutrient-scarce niche. We anticipate that our findings will broadly inform studies of evolution under extreme nutritional conditions and provide insights into the role of host-microbiota interactions in resilience to food scarcity at varied time scales.

## Introduction

Positive interactions between the host and gut microbiota are typical and crucial for the host’s development, growth, and typical functioning of the immune and nervous systems^5–13^. These beneficial couplings between the host and gut microbiota are thought to be shaped through their ecological/physiological needs and evolutionary history^14^. However, few studies have addressed evolutionary changes in such beneficial couplings, particularly the emergence of by-product mutualism between the host and gut microbes^15,16^. One possible approach to address this gap is to investigate host animals whose ancestral-type population lives in a typical environment while the derived population resides in nutrient-limited environments. For example, those that colonize diet-limited caves where they must adapt to chronic nutrient scarcity to grow and survive.

The Mexican tetra fish, *Astyanax mexicanus*, is an established experimental platform, consisting of two distinct morphotypes: surface-dwelling sighted populations (surface fish) and cave-dwelling blind populations (cavefish). Cavefish ancestors diverged from surface populations approximately 20,000 - 190,000 generations ago (generation time is about one year) by colonizing the perpetual darkness and limited-food environments^17–20^. Cavefish ancestors were likely forced to shift their dietary compositions from surface diets (crustaceans, fishes, and plant/algae) to bat guano and cave arthropods^21,22^. This dietary shift and the food scarcity of caves, where cave *A. mexicanus* resides, are thought to have driven numerous changes, including shifts in foraging strategy, reduced sleep and sociality^23–29^, and physiological changes, including reduced eyes, enhanced fat storage and fasting resistance^1,2,30–34^. This dietary shift would also induce a drastic change in the gut microbiota—the composition of the gut microbiota in cave populations differed significantly from surface populations in the wild^35,36^. Notably, when the same diets in a laboratory-based controlled condition, gut microbiota composition became very similar between surface fish and cavefish (alpha and beta diversities)^36^, yet cavefish still grew faster than surface fish. Previous attempts to detect the quantitative trait locus (QTL) for growth around genomic loci of growth hormone and IGF-1 pathways failed^39^. Introducing the cavefish-type P211L mutation of *insra* increased body length of zebrafish, but its effect was subtle^34,39^.

This fact implies unknown mechanisms that support rapid growth in cavefish, such as an evolved mechanism for nutrient intake from the gut microbiota.

Here, we reveal the evolutionary emergence of a beneficial relationship between host and gut microbiota that enables the host to thrive on a low-nutrient diet and supports resilience to chronic fasting. We hypothesized that the evolution of gut microbiota-host coupling facilitates adaptation to a leaner environment through encouraging body growth on a leaner diet and supporting fasting resilience. To capture such relationships, we applied two experimental interventions in both surface and cave *Astyanax*: (1) feeding nutritionally lean to rich diets to juvenile fish, and (2) fasting adults for one month. We then applied a new pipeline that overcomes difficulties in analyses of typical 16S ribosomal RNA (rRNA) gene amplicon data, conservatively interpreting 2-D visualized data structure (*e.g.*, skewing distance matrix^40^), and accounting for 0-inflated data with small sample sizes (N < 100)^41–43^. This strategy identified growth-associated gut-enriched metabolic pathways in developing juveniles, and acute responses of microbes in fully developed adults. Our results demonstrate an evolved by-product benefit from gut microorganisms that compensates for growth and energy sources in a lean environment.

## Results

### A lower-calorie diet promoted body growth over a high-calorie diet in cavefish

To test cavefish growth on the lean diet, we used 1-2-month-old juvenile fish, which are 1-2 cm in standard length, can swim freely in the middle of the water column, and are sexually immature. These fish were fed nutrient-rich live brine shrimp larvae until 1 month old (See Materials and Methods). Then, we fed three distinct diets: live brine shrimp larvae (continued to feed since 1 month old; 5.9 kcal/g)^44^, spirulina powder (3.3 kcal/g)^45^, and bat guano (unknown caloric amount), representing the nutrient-rich juvenile diet, leaner protein-rich diet, and the leanest diet similar to what cavefish may eat in caves^21^, respectively. We tested surface fish (a lineage-1 surface population^17^), Molino cavefish (blind cavefish; phylogenetically close to the lineage-1 surface fish, living in the moderately lean Molino cave^17,46^), and Pachón cavefish (the most derived blind cave population, living in a lean Pachón cave^17,46^). In addition, to highlight the involvement of genetic factors, we used the F_1_ hybrids of surface fish and Molino cavefish (F1_SM_), and F_1_ hybrids of surface fish and Pachón cavefish (F1_SP_) (Fig. 1). Due to the complexity of the experimental design, we excluded Tinaja cavefish from this growth study, though they were used in the fasting experiment.

**Fig. 1.**
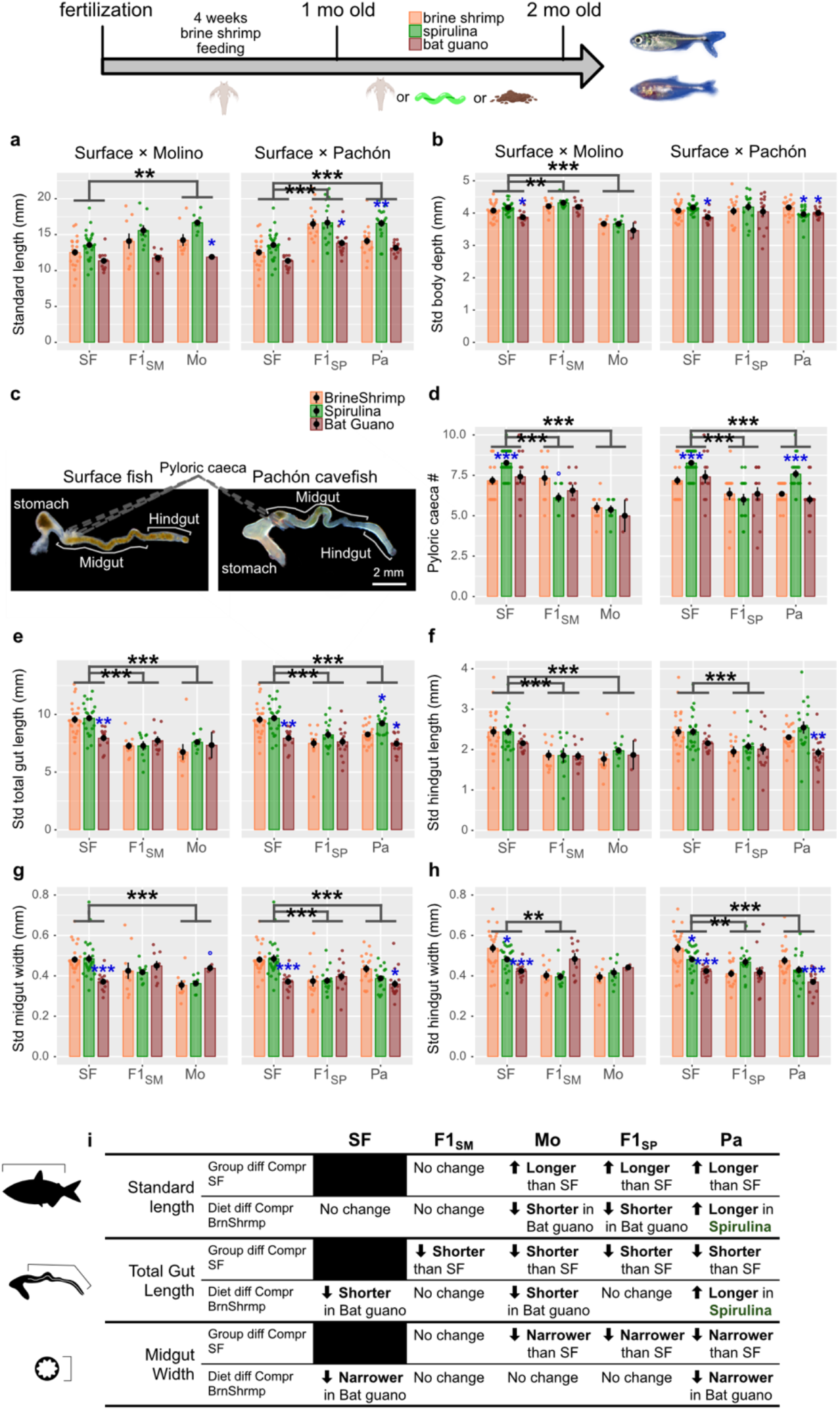
A lower-calorie diet and spirulina encouraged body growth in cavefish. **a, b**, The body standard length (**a**) and body depth standardized with the body standard length (**b**) of Surface fish (SF), Molino cavefish (Mo) and their F_1_ hybrid (F1_MO_) (left panels), and SF, Pachón cavefish (Pa) and their F_1_ hybrid (F1_SP_) (right panels), after a month of diet treatment. The feeding timeline is shown at the top. The populations and diet effects were compared via ANOVA, and if differences were detected, the post hoc comparisons were performed between surface fish vs. cavefish, and surface fish vs. their F_1_ hybrids for the population/groups, and brine shrimp vs. spirulina, and brine shrimp vs. bat guano for diets. Holm’s correction in the post hoc comparisons was applied for all these and the following studies. Blue stars indicated the significance compared with the brine shrimp diet. **c,** The gut morphologies of SF (left) and Pa (right). Measured areas are indicated in the pictures. **d-h**, Number of pyloric caeca (**d**), total gut length (midgut + hindgut) standardized with the body standard length (**e**), standardized hindgut length (**f**), standardized midgut width (**g**) and standardized hindgut width (**h**) of Surface fish (SF), Molino cavefish (Mo) and their F_1_ hybrids (F1_MO_) (left panels), and SF, Pachón cavefish (Pa) and their F_1_ hybrids (F1_SP_) (right panels). *: P < 0.05, **: P < 0.01, ***: P < 0.001 after post-hoc Holm’s correction. **i**, Summary of the morphological changes based on population/F_1_, and diets.

After 1 month of diet treatment, Molino and Pachón cavefish exhibited longer standard lengths than surface fish (Fig. 1a and Extended Data Fig. 1). Interestingly, Pachón cavefish grew larger under the spirulina diet feeding (intermediate caloric amount) compared to the brine shrimp diet (the right panel of Fig. 1a and Extended Data Fig. 1). Molino cavefish exhibited a similar trend relative to surface fish, although its growth rate with the spirulina diet was not significantly increased compared to the brine shrimp diet (the left panel of Fig. 1a). F1_SM_ and F1_SP_ hybrids exhibited comparable body length to their original cave populations instead of the surface population, suggesting cave allele dominancy (Fig. 1a and Extended Data Fig. 1), and brine shrimp feeding produced standard lengths indistinguishable from spirulina feeding (F1_SP_, Fig. 1a).

In contrast to standard length, body depth was not significantly increased under the spirulina feeding regime. Pachón cavefish showed reduced body depth comparable to that on the guano diet (Fig. 1b, right panel), indicating that the spirulina diet promoted growth in the longitudinal axis in Pachón. Molino cavefish had lower body depth than surface fish in any diet treatment (Molino in the left panel of Fig. 1b). Molino or Pachón hybrid F1s did not significantly differ in their body depths; therefore, larger fish had larger body depths (Fig. 1b and Extended Data Fig. 1).

In summary, spirulina feeding promoted fish growth along the longitudinal axis in the Pachón cave population, while bat guano feeding slowed growth in the tested groups (Fig. 1a). F1 hybrids in the longitudinal axis showed similar diet responses to their parental populations (F1_SM_) or cave population (spirulina in F1_SP_) and grew better under the brine shrimp feeding (F1_SP_) (Fig. 1a). Therefore, diet responses include interactions with genetic factor(s) of Molino and Pachón cavefish.

Gut morphology parameters, including pyloric caeca number, gut length, and gut width, similarly increased under spirulina feeding (Fig. 1c-h). Pyloric caeca are small pouches between the stomach and midgut that are thought to digest and absorb nutrients by expanding the surface area of the gut^47^, and act as a center of immune reactions^48^. Cave populations generally exhibited fewer pyloric caeca across diets than surface populations (Fig. 1c,d). Interestingly, spirulina feeding increased their number in surface and Pachón cavefish but not in Molino or F_1_ hybrids (Fig. 1d), indicating that homozygotes of surface fish alleles or Pachón cavefish alleles increase the number of pyloric caeca in response to the spirulina diet. The cavefish gut lengths (*i.e.,* midgut + hindgut lengths) standardized by body standard length were shorter than the surface fish (Fig. 1e, Extended Data Fig. 1). In Pachón cavefish gut length was longer under the spirulina feeding and shorter under the bat guano feeding, while the effect of the spirulina diet was undetectable in surface fish, F1s, or Molino cavefish (Fig. 1e). The midgut length, a subdivision of the gut length, varied almost no changes in response to that of the overall gut length (not shown), and the hindgut length exhibited less responses to the diet treatments (Fig. 1f). Gut widths were reduced under the bat guano feeding in surface fish and Pachón cavefish (Fig. 1g,h).

In summary, standardized gut length and width of the two cave populations were shorter (Molino and Pachón) and narrower (Pachón) than those of surface fish. Only Pachón cavefish had increased gut length in response to spirulina feeding. The bat guano diet generally reduced growth in both gut length and width. Thus, these distinct dietary responses indicate gene × diet effects on both body and gut growth.

Increased growth on spirulina diet in the Pachón body lengths was surprising because the brine shrimp diet (5.9 kcal/g) provides more calories than the spirulina diet (3.3 kcal/g) (Fig. 1i)^44,45^. Each diet was given *ad libitum.* We thus hypothesized that Pachón cavefish evolved the capacity to support and even increase growth on lean diets, perhaps through novel relationships with gut microorganisms.

### Topological Data Analysis (TDA) reveals hidden associations between morphological traits and gut microorganisms

The gut microbiota is a crucial interface between diet and the host gut epithelia, mediating complex nutritional exchange^49,50^. Mid- and hindgut microbial taxa of individuals from the dietary study were thus analyzed (Extended Data Fig. 2a-b), as well as environmental samples: the diets and housing water (Extended Data Fig. 2c), using sequencing of the variable region 4 (V4) of the 16S ribosomal RNA gene. Proteobacteria (now called Pseudomonadota) and Fusobacteria (now called Fusobacteriota) are a large proportion of the gut microbiota of surface fish and cavefish (Extended Data Fig. 2a-b), which is consistent with prior work in *A. mexicanus*^36^. Gut microbiota compositions were distinct from those of the environments: the housing water and diets (Extended Data Fig. 2c). For example, proteobacteria were the major phyla in the brine shrimp diet (> 70%) and housing water, which were not so consistently dominant in the gut, indicating that a selective mechanism exists even in the ∼2 cm-long juvenile fish.

Surface fish had higher Shannon-Weaver index alpha-diversity compared with Pachón cavefish, and bat guano feeding induced higher alpha-diversity in surface fish and F1 hybrids (Extended Data Fig. 2d,e). Diet and tank water displayed higher alpha-diversities than gut samples, with the exception of the spirulina diet with a low Shannon-Weaver index score (Extended Data Fig. 2f). Since the pattern of gut alpha diversity induced by diet (low in brine shrimp, and high in bat guano) was consistent in each fish group, gut alpha-diversity may be highly dependent on diet at this growth stage. Because the hindgut’s relative abundance (Extended Data Fig. 2b) and alpha-diversity (Extended Data Fig. 2e) were comparable to those of the midgut, and the midgut showed higher diversity (Extended Data Fig. 2a,d vs. Extended Data Fig. 2b,e), we focused on the midgut in the following studies.

Beta-diversity at the order level exhibits the best clustering resolution in non-metric multidimensional scaling (NMDS) and indicates that population groups (Extended Data Fig. 2g) and diet groups (Extended Data Fig. 2h) largely overlap with each other. Genus- and ASV-level comparisons produce similar results (Extended Data Fig. 2i-l). This indistinguishable diversity among populations was consistent with former reports using laboratory fish^36^, or wild-caught surface fish (Micos) and cavefish^35,36^. These low-structured results obstructed testing the hypothesis that beneficial coupling between host and gut microbiota supports cavefish growth. Thus, we introduced a two-step analysis that consists of (1) visualizing the data structure with topological data analysis (TDA) mapper^51,52^ and subsampling individuals based on their gut microbiota similarity along with the phenotypic score of interest (*e.g*., fish length), followed by (2) sparse modeling regression using this subsampled data by regressing the gut microbial abundances against the host phenotypic score (Supplementary Methods; Extended Data Fig. 3).

Visualization methods for high dimensional ASV data such as PCoA or NMDS compress samples into two or three dimensions and discard the variance carried by the unplotted axes; the TDA mapper instead retains every sample and renders the data as a connectivity graph, in which each vertex is a cluster of compositionally similar individuals and links mark similarity between clusters (Extended Data Fig. 3a–c; Supplementary Method). Applied to our data, TDA recovered the intermingled distribution of populations and diet treatments already evident in the NMDS (Extended Data Fig. 3a,b vs. Extended Data Fig. 2g-l), but additionally made explicit which compositional clusters of individuals are contiguous with others. Using a host trait, such as standard length, as the filter function revealed individuals of comparable body length carrying distinct ASV compositions and allowed us to extract subsets spanning the trait range (cyan and gray shades in Extended Data Fig. 3d). To each subset, separated by genetic background (SF, F1_SM_, Mo, F1_SP_, and Pa) but pooled across all diet conditions, we applied sparse modeling, which is tractable where ASVs (∼1,000) greatly outnumber samples and evaluates all ASVs within a single model (Extended Data Fig. 3e; Extended Methods). The retained taxa therefore constitute a combination whose abundances jointly account for trait variation, consistent with pooled microbial community activity rather than any single taxon acting alone^53,54^. The same procedure was applied to PICRUSt2-estimated pathway abundances.

### Topological mapping and sparse modeling methods consistently identified fish-morphology-associated gut microorganisms under a mild antibiotic disruption

The morphological trait-ASV regression analyses detected conserved couplings shared between surface fish and Pachón cavefish, including *Plesiomonas* (negative association with body length) and *Legionella* (negative association with gut length) (Extended Data Fig. 4a). In contrast, several unique couplings were identified in Molino and Pachón cavefish, which were not observed in surface fish. Specifically, positive associations with body length emerged with the genera of *Comamonadaceae* and *Delftia* in Molino, and *Candidatus-Arthromitus*, *Microbacteriaceae*, *Micrococcaceae* and *Pseudonocardia* in Pachón cavefish (Extended Data Fig. 4a). Our pipeline consistently detected gut microbial taxa associated with fish morphological measurements even after the gut microbiota were disturbed by a mild antibiotic intervention (Extended Data Fig. 5a-m; boldface taxa in Extended Data Fig. 4d-f). This proof-of-concept experiment supports the consistency of the TDA mapper plus sparse modeling method.

### The most derived cavefish, Pachón cavefish, evolved new associations between body length and gut-enriched short-chain fatty acid pathways, promoting growth on a lower-calorie diet

We used the same strategy to explore the functional diversity among coupled gut taxa^55–58^, estimating abundances of metabolic pathways present in microbial genomes inferred from ASV sequences^59^ (Extended Data Fig. 3d).

For fish standard length (Fig. 2), we analyzed each genetic background separately (*i.e.,* Surface fish, Molino cavefish, Pachón cavefish, F1_SM_ and F1_SP_). Within each genetic background, diet conditions (brine shrimp, spirulina, bat guano) were taken into account to estimate the microbial metabolic pathways associated with standard length (Extended Data Fig. 3d). Pachón cavefish exhibited 43 positively and 11 negatively associated/coupled metabolic pathways during the growing stage (Fig. 2a,b; 2 months old), which was the most among the five genetic backgrounds (Fig. 2b and Extended Data Fig. 6). Many of the estimated coupled metabolic pathways are in positive association in Pachón cavefish, that is, larger body size corresponds to a greater abundance of pathways, including acetylene degradation and glycogen/starch degradation (Fig. 2a,b). The standard length-coupled pathways could benefit fish growth through two metabolic categories, pyruvate generation and short-chain fatty acid (SCFA)-production (*i.e*., fermentation) (Fig. 2a and Extended Data Fig. 7)^60^. The estimated pyruvate generation and SCFA-directly-producing pathways appeared with high positive *r* values (Fig. 2a). Pyruvate is the central precursor for microbial SCFA synthesis^61^. Thus, the top-listed pyruvate-generation and SCFA-directly-producing pathways imply that a high abundance of SCFAs (*e.g.,* acetate) may promote fish growth.

To test this, we supplemented fish diet with β-hydroxybutyrate (BHB), a short-chain monocarboxylate that, like acetate and lactate, is oxidized to acetyl-CoA and enters the TCA cycle. One month of BHB supplementation accelerated growth in Pachón cavefish but slowed growth in surface fish (Extended Data Fig. 7a). To further test whether these identified pathways promote body growth in only Pachón cavefish and not surface fish or Molino cavefish, we pharmacologically targeted the pyruvate-generating pathway via quercetin (blue characters in Fig. 2a; Extended Data Fig. 7b)^62–64^. We supplemented quercetin in the spirulina diet, where pyruvate generating pathways were estimated as positively coupled with standard length in Pachón cavefish (see Fig. 2a ‘abundance’ column, and the green regression lines of Pachón row in Extended Data Fig. 7c). Through a course of 4 weeks of supplementation of quercetin, surface fish, Pachón, and Molino cavefish grew significantly on the control diet (body standard length and growth rate: black lines in Fig. 2c). However, the growth rate of the quercetin-fed Pachón cavefish was slowed (red lines in Fig. 2c), while there was no detectable effect of quercetin in surface fish and Molino cavefish (Fig. 2c). The quercetin-induced growth retardation was only detectable in the longitudinal axis (standard length; Fig. 2c) but not in body depth (Extended Data Fig. 7d), indicating that the inhibited pathway contributes to longitudinal growth.

**Fig. 2.**
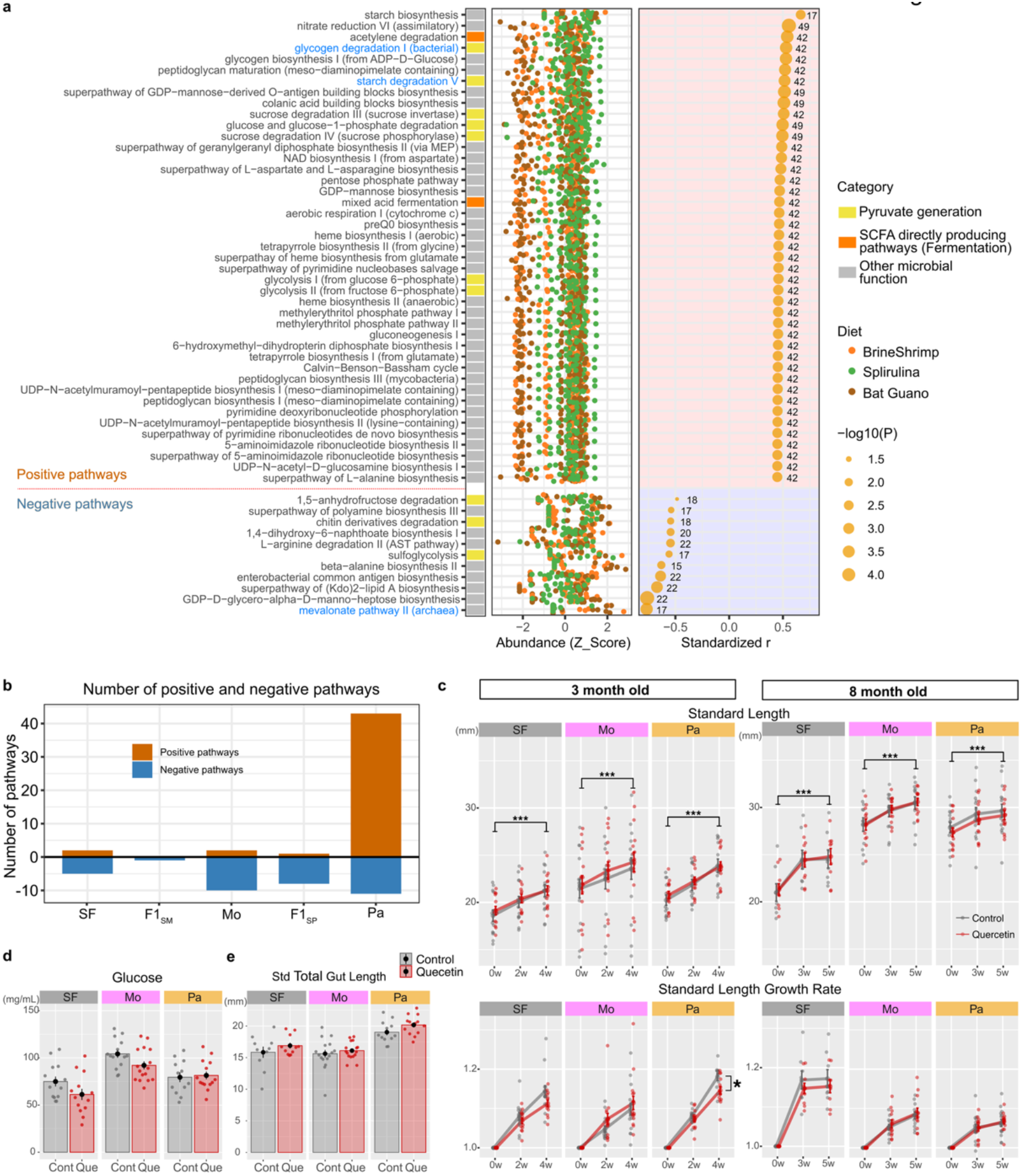
Pachón cavefish evolved new positive associations between body standard length and their gut-enriched pyruvate/short-chain fatty acid pathways, supporting growth upon a lower-calorie diet. **a**, Significant associations/couplings of gut-enriched metabolic pathways with the body standard length in Pachón cavefish. The gut-enriched pathways were aligned according to the levels of Pearson’s correlation coefficients (*r*: univariate regression, far right panel). To obtain the widest variation in body standard length scores for sparse modeling regression, all fish individuals treated with three distinct diets per population were used for the sparse modeling regression. Circle sizes represent the -log_10_(P-values of the correlation coefficient). The z-score of each pathway’s abundance (PiCRUSt2 estimated) is shown in the middle column. Colors of the dots are as follows—orange: brine shrimp, green: spirulina, and brown: bat guano. These gut-enriched pathways were also grouped as “Pyruvate generation”, “SCFA direct producing pathway” and “Other microbial function”—custom-made categories based on the MetaCyc ontology (far left panel; Supplementary Table 1). Blue colored pathways were targeted by quercetin pharmacological treatment used in **c-e**. **b**, The number of positively and negatively associated/coupled pathways in different populations (surface fish [SF], F1_SP_, Pachón [Pa], F1_SM_, and Molino [Mo]). Pachón cavefish enriched-pathways were dominated by positive pathways, while an ancestor-like surface fish, another cave population (Molino), and F_1_ hybrids (F1_SP_ and F1_SM_) exhibited fewer and more negative associations. **c**, Pharmacological inhibition of pyruvate generation pathways via quercetin resulted in growth retardation in Pachón cavefish. (Top row) Shifts of body standard length during 4 weeks of the control and quercetin diet treatment, starting from 3-month-old (top-left, juveniles) and 8-month-old (top-right, young adult). (Bottom row) Growth rates of standard length starting from 3-month-old juvenile fish (bottom-left) and 8-month-old adult fish (bottom-right). Quercetin reduced the growth rate in Pachón cavefish after 4 weeks, while no effect was detected in surface fish and Molino in juveniles. Also, no effect was detected in adult fish. **d,e**, Quercetin treatment did not significantly reduce blood glucose levels (**d**) or the standardized total gut length (**e**). *: P < 0.05, **: P < 0.01, ***: P < 0.001 after post-hoc Holm’s correction.

Quercetin is known to reduce blood glucose levels in humans. However, our dose of quercetin treatment did not affect the blood glucose level in the three populations tested (Fig. 2d), or other morphological traits (*e.g.,* gut length; Fig. 2e). Quercetin is also suggested to inhibit the mevalonate pathway II^65^ (Fig. 2a), whose inhibition may be negligible for fish growth. Together, we concluded that suppressed growth in quercetin-treated Pachón cavefish was consistent with a contribution of pyruvate-to-short-chain monocarboxylate metabolism to Pachón growth.

To explore differences in blood metabolites between surface fish and Pachón cavefish as mediators between their gut microbiota and body tissues, we fed a standard Zeigler zebrafish diet for 5 weeks *ad libitum* during the 3-month-old growing period and collected blood sera 2 hours after feeding^3^. The 2-hour point after feeding represents a peak of blood metabolites in response to feeding^34^. The sera of Pachón cavefish showed no increase in ketone bodies or SCFAs, perhaps due to fast uptake in tissues (Extended Data Fig. 7e; see Discussion). Additionally, the group of free sugars was significantly more abundant in Pachón cavefish than in surface fish (enrichment q = 0.026, p = 0.019, although no significance in each individual free sugar; Extended Data Fig. 7e). Free sugar was abundant in Pachón cavefish, but did not mirror the abundance of TCA cycle metabolites (*c.f.*, lower abundances of TCA intermediate metabolites than in surface fish; Extended Data Fig. 7e). Similarly, free amino acids were more abundant in Pachón cavefish, but their oxidated product, branched-chain amino acid (BCAA) catabolite, which can be catabolized to acetyl-CoA to fuel the TCA cycle, was lower than in surface fish (Extended Data Fig. 7e). Both TCA metabolite and BCAA catabolite findings imply that these free sugars and amino acids are not quickly oxidized or consumed as energy fuel in Pachón cavefish, but could be stored^3^ for another purpose, such as growth.

In summary, the topology mapping plus sparse modeling analysis predicts the coupled metabolic pathway well, revealing an evolved coupling between host growth and the gut microbiota that promotes the growth of Pachón cavefish on a lower-calorie spirulina diet. Such host-gut microbial coupling was not detected in surface fish, Molino cavefish, or F_1_ hybrids, thus, homozygosity for Pachón alleles significantly promotes growth on a low-calorie spirulina diet. We also addressed whether gut tissue morphology, such as gut length, would be coupled with gut microbiota-enriched pathways (see Supplementary Discussion; Extended Data Figs 8, 9). Briefly, surface fish gut length was estimated to be coupled with many gut-enriched pathways, but F1s and cavefish exhibited a very limited number of coupled pathways, suggesting that cavefish lost those gut length-associated pathways during evolution.

These findings may explain the rapid growth of juvenile fish during the rainy season in México, when cave pools receive organic matter from outside. Next, we examine how the gut microbiota may support the fasted state that adults may experience during the dry season.

### Fasting reduced liver weights and blood glucose in both surface and cave populations, and also reduced gut sizes in cave populations

To evaluate the diversity of adaptation to fasting, we included the Tinaja cavefish, which independently diverged from the surface fish population (lineage 1)^17^ at a phylogenetic distance from surface fish similar to that between surface fish and Pachón cavefish. Tinaja is known to have evolved an insatiable appetite^30^ and enhanced lipogenesis^2^. One month of fasting in adult fish (1-2 year old) slowed growth in body standard length (Fig. 3a) and reduced body depth (Fig. 3b), but such suppressions were subtle, possibly because they were fully developed adults (Fig. 3a,b and Extended Data Fig. 10). Fasting also showed a limited effect on weight loss (Fig. 3c) and reduced blood glucose level (Surface fish, and Tinaja and Pachón cavefish, Fig. 3d). Liver weight after one month of fasting exhibited drastic reductions in the surface, Tinaja, and Pachón populations, but not Molino cavefish (Fig. 3e). Gut size reduction under fasting was more pronounced in Tinaja and Pachón cavefish than in surface fish and Molino cavefish (Fig. 3f,h,i and Extended Data Fig. 10), while no detectable change under fasting was observed in pyloric caeca number (Fig. 3g).

**Fig. 3.**
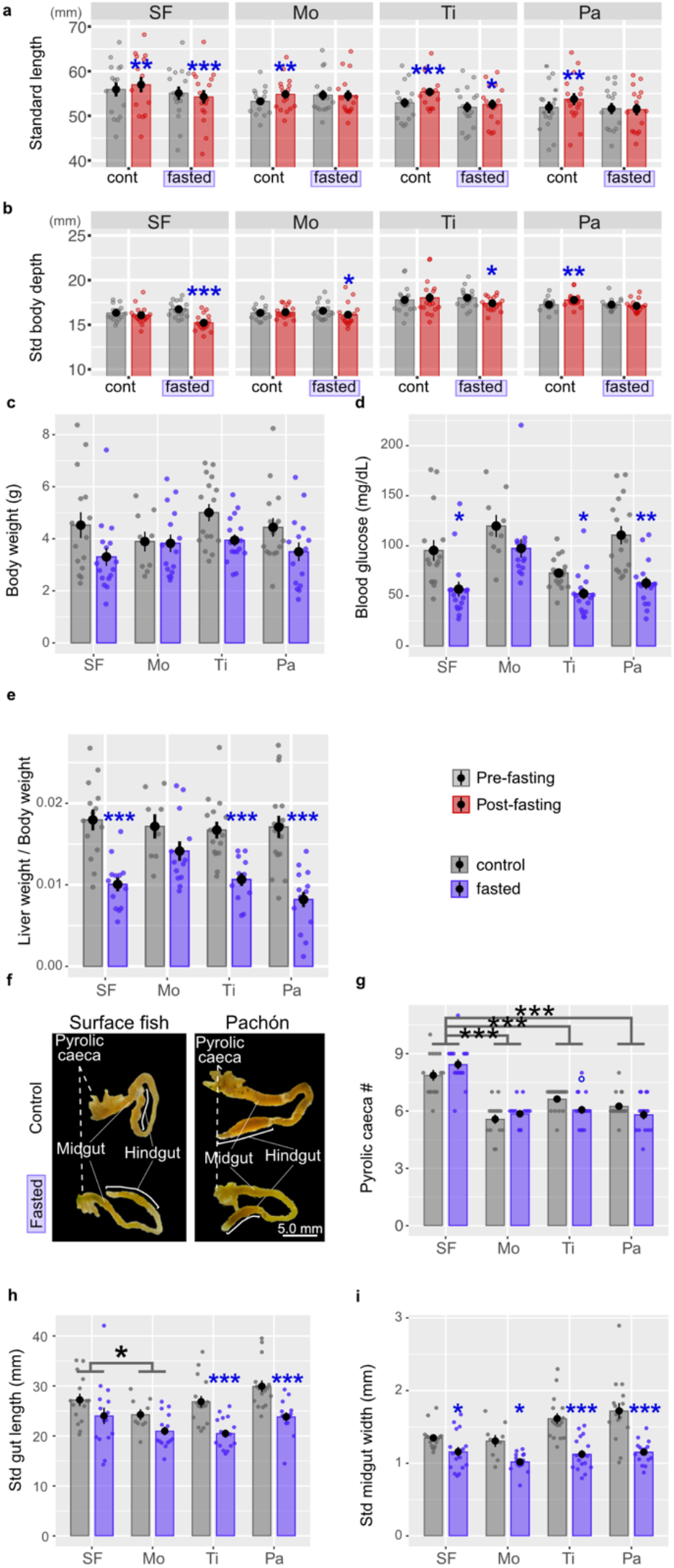
One month of fasting reduced blood glucose levels and liver weights in both surface fish and cave populations and also reduced gut sizes in cave populations. **a, b**, The body standard length (**a**) and body depth standardized with the body standard length (**b**) of surface fish (SF), Molino (Mo), Tinaja (Ti) and Pachón cavefish (Pa). Blue stars indicate a significant decrease compared with the control/fed group. **c-e,** Body weight (**c**), blood glucose (**d**) and relative liver weight (liver weight/body weight) (**e**) of SF, Mo, Ti and Pa, indicating significant decreases in blood glucose and liver weight but an undetectable decrease in body weight in most populations. **f**, Gut structure of SF (left) and Pa (right). **g-i**, Number of pyloric caeca (**g**), standardized total gut length (**h**) and standardized midgut width (**i)** of SF, Mo, Ti and Pa. Blue stars indicate the significant decrease compared with the control/fed group. More pronounced decreases of the gut length and width were observed in Tinaja and Pachón cavefish compared with surface fish and Molino cavefish. *: P < 0.05, **: P < 0.01, ***: P < 0.001, °: P < 0.05 before post-hoc Holm’s correction but P > 0.05 after the correction.

Under fasting conditions, the gut microbiota of surface fish and Pachón cavefish showed increased alpha diversity (Extended Data Fig. 11a-d). The beta-diversity plot again did not discriminate populations, though fed and fasted groups showed slight separation in their clustering (Extended Data Fig. 11e; reduced proportion of Firmicutes in fasted fish in Extended Data Fig. 11a,b).

### A loss of liver weight mirrored increases in aromatic compound degradation pathways in Pachón cavefish

The liver gains weight by storing glycogen and fat in response to rich feeding, and loses it during fasting by catabolizing glycogen and fat (Fig. 4a and Extended Data Fig. 12b)^2,30,33^. Thus, we surveyed PICRUSt2-estimated gut-enriched pathways associated with changes in liver weight under fasting conditions. Although the alpha diversity of gut microbiota increased in both surface fish and Pachón cavefish (Extended Data Fig. 11c,d), the number of liver weight-coupled pathways was only pronounced in Tinaja and Pachón cavefish, not in surface fish (Fig. 4b).

**Fig. 4.**
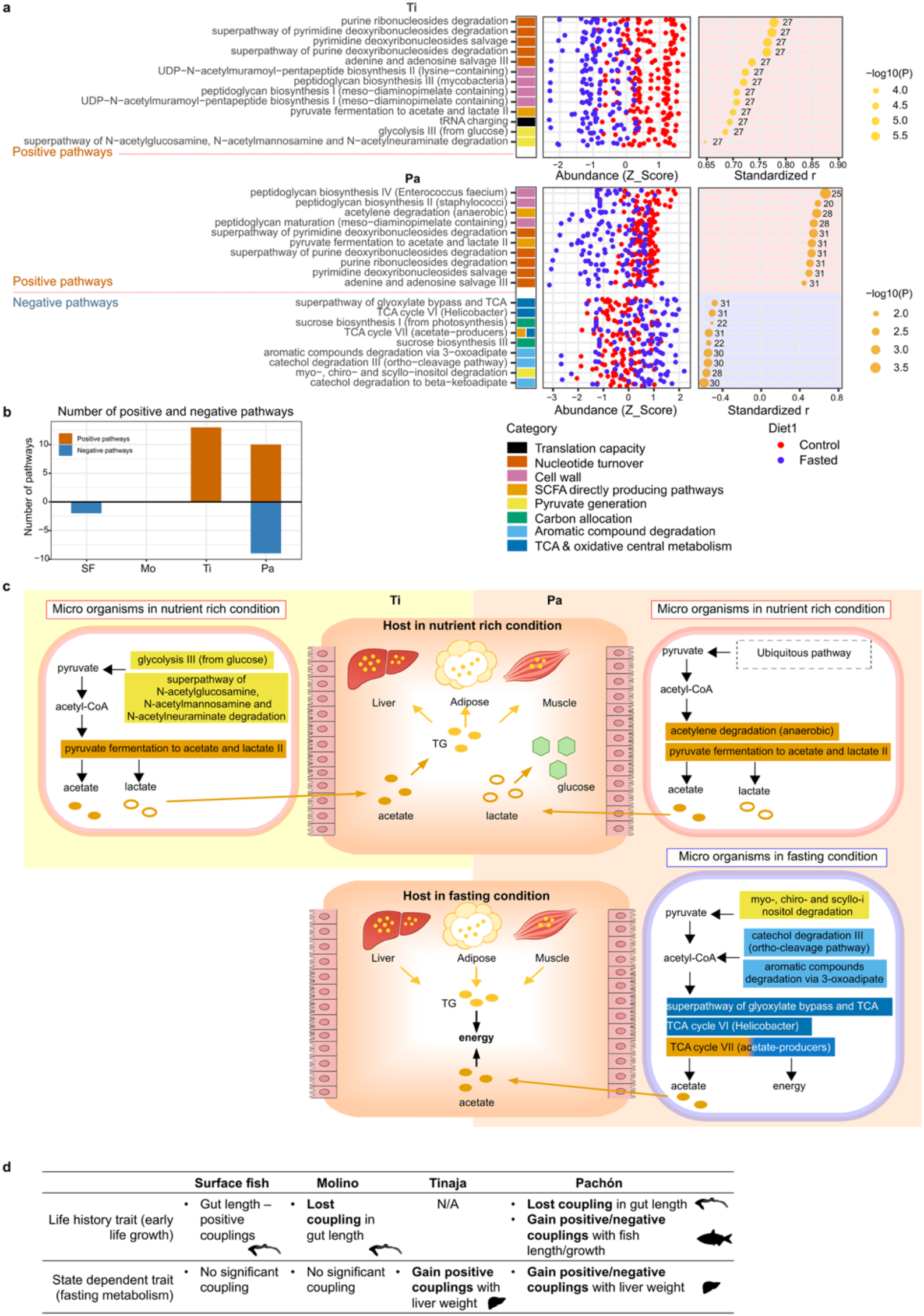
Liver weight loss mirrored reductions in the cell proliferation-related pathways in gut microbiota of Tinaja and Pachón cavefish, and Pachón cavefish further evolved couplings with aromatic compound degradation increased during liver weight loss. **a,** Significant associations/couplings of gut-enriched metabolic pathways with the liver weight 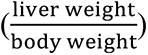 in Tinaja and Pachón cavefish. The gut-enriched pathways were aligned according body weight to the levels of Pearson’s correlation coefficients (*r*: univariate regression, far right panel). To obtain the widest variation in liver weight values for sparse modeling regression, all fish individuals treated with the fed and fasted conditions per population were used for the sparse modeling regression. Circle sizes represent the -log_10_(P-values of the correlation coefficient). The z-score of each pathway’s abundance (PiCRUSt2 estimated) is shown in the middle column. Colors of the dots are as follows—blue: fed control, red: fasted. These liver weight-coupled pathways were also grouped as “Translation capacity”, “Nucleotide turnover”, “Cell wall”, “SCFA direct producing pathway”, “Pyruvate generation”, “Carbon allocation”, “Aromatic compound degradation”, and “TCA & oxidative central metabolism”—custom-made categories based on the MetaCyc ontology (far left panel; Supplementary Table 4). **b**, The number of positively and negatively associated/coupled pathways in different populations (Surface fish [SF], Molino [Mo], Tinaja [Ti] and Pachón [Pa]). Ti and Pa have evolved positively coupled pathways, and both positively and negatively coupled pathways, respectively. In contrast, SF and Mo have no pathway coupled with liver weight. **c,** Hypothetical model of the host-gut microbiota interaction under either of the fed or fasted conditions in Ti and Pa. In a food-abundant season of a year, Ti and Pa store triglycerides (TG) in tissues by possibly using circulating acetate, including the ones derived from gut microorganisms. Conversely, during a food-limited season, stored TG may be catabolized by beta-oxidation and short-chain fatty acids (SCFAs) become circulated in Ti and Pa (liver weight loss), while Pa may still gain SCFAs from their gut microorganisms via a result of aromatic compound degradation (source can be debris of plants, percolating water, plant-eating bat guano, and/or biofilms), which the host cannot process. **d**, Summary of the early life growth and fasting metabolism and their couplings to the gut microbiome among the tested four populations.

Within these liver-weight-coupled pathways, liver weight gain is associated with an increase of cell proliferation and maintenance, and SCFA generating fermentation pathways in Tinaja and Pachón cavefish (Fig. 4a). These estimated positively associated pathways can release acetates which may be incorporated to synthesize fat for storage in the liver and other tissues (Fig. 4c). Pachón cavefish further evolved a coupling with aromatic compound degradation which could prevent liver weight loss (*r* < 0 negative association; Fig. 4a). These aromatic compounds are likely abundant in decaying plants^66^ and insect-eating bats’ guano^67^, and present in percolating water^68^ and biofilm metabolites^69^. Usually, animals cannot metabolize or catabolize these compounds^69^, but some gut-associated microbes can. One-month of dietary supplementation of the reaction intermediate of the aromatic compound degradation, 4-hydroxybenzoate (4-HBA, Extended Data Fig. 12c), decreased the liver weight in the fed Pachón cavefish. The reduced liver weight was surprising because Pachón cavefish store glycogen and fat in the liver, muscle, and other tissues during fed conditions^2,33^. In contrast, no significant effect on liver weight was detected in fasted 4-HBA-treated Pachón fish (Extended Data Fig. 12d,e), while we observed a sign of slowing body weight loss in the fasted condition (the bottom panel of Extended Data Fig. 12e).

In summary, we found numerous evolved couplings between liver weight and gut microbiota-enriched metabolic pathways in Pachón and Tinaja cavefish, but not in surface fish or Molino cavefish. The gut microbiota of Tinaja and Pachón cavefish may supply SCFA to support fat storage during nutrient-rich conditions to prepare for the lean season (Fig. 4c). In contrast, in fasting Pachón cavefish, gut microorganisms may help degrade host-inaccessible aromatic compounds, thereby supplying SCFA to the host while catabolizing fat and glycogen in the liver in the fasted condition (Fig. 4c).

Finally, to observe whether microbes carrying these detected pathways exist in the natural cave environments of these populations, we investigated the water microbiota of their native cave pools. The Pachón and Tinaja cave water samples were analyzed by 16S rRNA gene sequencing (Fig. 5a), and the same genera that were coupled with growth and fasting (pyruvate-to-short-chain monocarboxylate and aromatic compound degradation pathways; genera and pathways were listed in Supplementary Table 5) were detected in the cave water (Fig. 5d,e), where Pachón cave had a higher relative abundance of these genera than Tinaja cave (Fig. 5d,e).

**Fig. 5.**
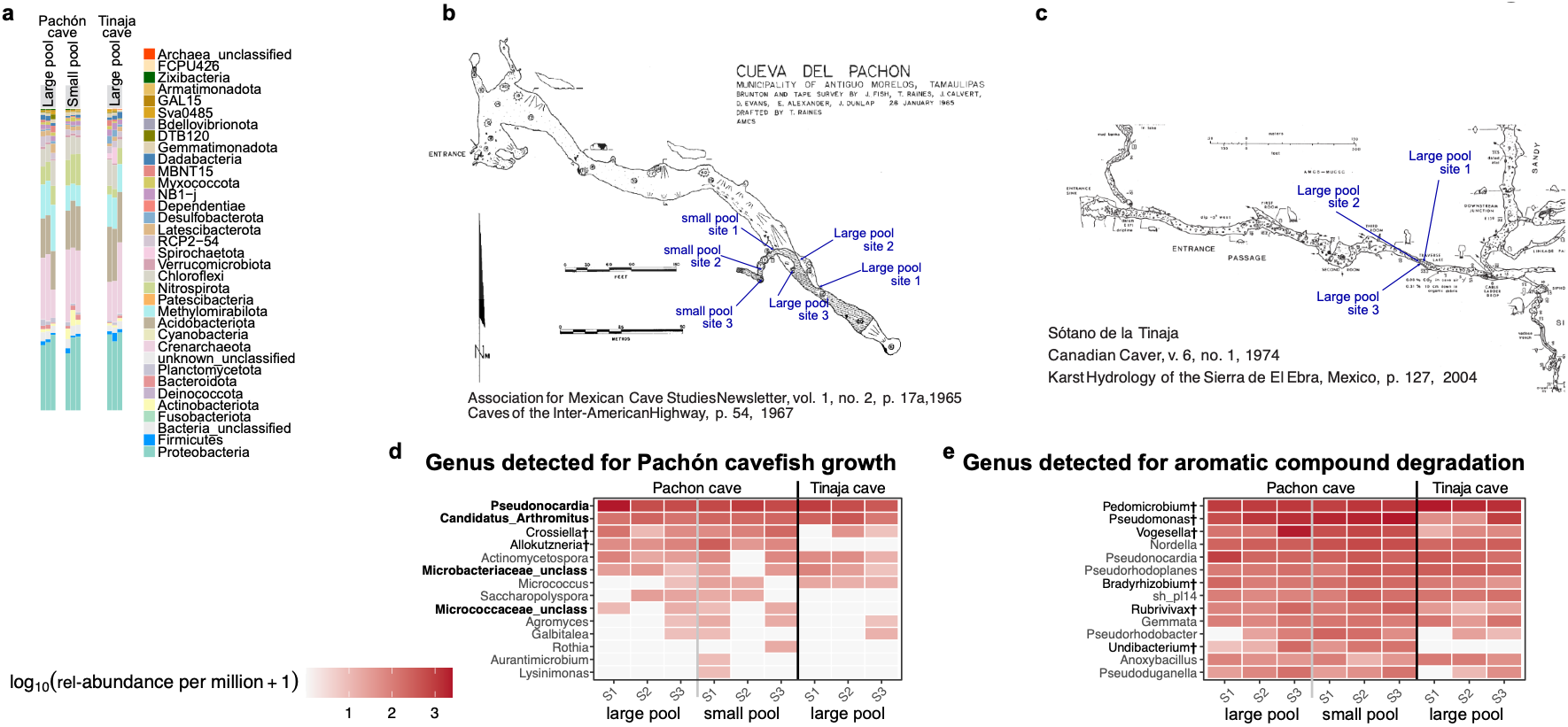
Microbiota in Pachón and Tinaja cave contain genera whose genomes can code pyruvate-SCFA and aromatic compound degradation pathways. **a,** Relative abundance of two Pachón cave pools (3 sampling sites in each), and a Tinaja cave pool (3 sampling sites). These samples exhibit similar profiles to each other. **b**, Pachón cave map based on the association for Mexican cave studies newsletter (1995). The approximate water sampling sites were indicated. The cave entrance is on the left. **c**, Tinaja cave map based on the Canadian Caver Report (1974). The approximate water sampling sites were indicated. The cave entrance is on the left. **d**, Relative abundance per million of genera that were positively associated for the body standard length, including *Candidatus-Arthromitus*, *Microbacteriaceae*, *Micrococcaceae*, and *Pseudonocardia* (Extended Data Fig. 4a). Other genera belong to the same families of these selected four genera. Pachón cave pool exhibit higher abundances of genera *Allokutzneria* and *Crossiella*. **e**, Relative abundance per million of genera whose genomes contain aromatic compounds degradation via β-ketoadipate, catechol degradation to β-ketoadipate, and/or myo-, chiro- and scillo-inositol degradation (contributing aromatic compound degradation and processing pyruvates to supply SCFAs) in Pachón and Tinaja cave pools. †: P < 0.05 between Pachón and Tinaja cave pools but P > 0.05 after post-hoc correction.

Overall, couplings between early life growth and gut microbiota evolved and promote Pachón cavefish growth in a low-caloric diet (Fig. 4d). The fasting-resilient Tinaja and Pachón cavefish evolved couplings with liver-weight-associated gut microbiota, which may encourage the storage of fat/glycogen (Tinaja and Pachón) and supply energy (SCFAs) via aromatic compound degradation (Fig. 4d).

## Discussion

In this study, we were interested in how animals evolved to overcome growth in a diet-restricted environment. Animals that colonize lean environments are known to shift their physiology and/or even reduce costly organs (*e.g.,* eyes) to save energy^1–3,30,31,70,71^. However, little is known about how commensal gut microorganisms contribute to such adaptation (*cf.,* insulin resistance promotes body weight^34^). Distinct nutritional inputs revealed growth ability found in Pachón cavefish but not detected in other genetic backgrounds: A lower-calorie diet, spirulina, led Pachón fish to grow the most among the tested diets. The 16S rRNA gene-based microorganism profiling indicated that a positive coupling between fish growth and gut-microbiota-enriched carbohydrate degradation pathways (involving glycogen, starch, and SCFAs) supports Pachón cavefish growth under low-calorie spirulina feeding. Also, adult fasting resilience in cavefish may be supported by gut microorganisms, which could promote fat storage under normal feeding and may supply some energy (*e.g*., SCFAs) by degrading aromatic compounds during fasting.

While our results offer exciting insights, it remains unclear how the cavefish gut recruits beneficial gut microbes or how the host genome evolved to use products of starch degradation and/or aromatic compound degradation produced by microbial metabolism.

### Cave adaptation

Results from the three-diet study indicate the evolution of new host-gut microbiota couplings that modulate body growth in Pachón cavefish. The candidate gut taxa positively impacting growth in Pachón cavefish are *Micrococcaceae, Microbacteriaceae*, *Pseudonocardia,* and *Candidatus-Arthromitus sp* (Extended Data Fig. 4). Elucidating the detailed molecular and metabolic interactions between these microbes and Pachón fish requires further study, but there is compelling evidence that suggests specific hypotheses. *Micrococcaceae* are aerobic, facultative Actinobacteria with documented proteolytic and lipolytic capacity, and can be found in fish intestine^72,73^, positioning them as candidate mobilizers of peptides, free amino acids (observed in Extended Data Fig. 7e), and free fatty acids. Free fatty acids can be further catabolized into short-chain fatty acids (SCFA) by other gut microbes via gut-enriched fermentation pathways identified here (Fig. 2a). *Microbacteriaceae* in the phylum of Actinomycetota includes species colonizing the surface of European cheeses and has been found in the fish intestine, possibly releasing lipolytic enzymes to degrade lipids^74^, which can eventually supply SCFAs as cooperative products processed with other taxa. *Pseudonocardia* in the phylum of Actinomycetota has a rich biosynthetic gene cluster and is a chemo-organo-heterotroph. It could also produce antibiotics, thereby impacting which microbes colonize the gut^75,76^. Lastly, (4) *Candidatus-Arthromitus*, a segmented filamentous bacteria (SFB) and Clostridia-related Firmicutes commensal bacteria, carries a highly reduced genome, which may not significantly contribute to producing SCFAs or other metabolites. However, this bacterium can promote immune and gut epithelial maturation, thereby supporting the uptake of gut metabolites^77^. We also note that *Cetobacterium* in the phylum Fusobacteriota is highly abundant among *Astyanax* populations (ASV_2 and ASV_3 in the three-diet experiment). *Cetobacterium* exhibits many metabolic capabilities, including the biosynthesis of short-chain fatty acids (SCFAs)^78^. Thus, it may be a major supplier of short-chain monocarboxylates to the host.

Although there are possible contributions to SCFAs from the microorganisms identified, serum metabolites under the standard diet did not indicate increased SCFAs or ketone bodies in Pachón cavefish (Extended Data Fig. 7e), but this data did not rule out the possibility that the gut microbiota supply acetate, lactate, and/or ketone bodies to the host. High expression of PPARγ^2^ and a higher number of adipocytes^3^ in Pachón cavefish may explain the indistinguishable levels of serum SCFAs and ketone bodies, due to rapid uptake of serum SCFAs and ketones to be stored as lipids^2,3,33^. Free sugars and free amino acids were abundant in Pachón cavefish, and may be supplied in part by their gut microbiota (Extended Data Fig. 7e). This abundance could also be due to a lower rate of uptake of these metabolites, supported by known Pachón cavefish insulin resistance based on an insulin receptor mutation (*insura*^34^), and lower abundance of downstream oxidation metabolites including TCA cycle metabolites (sugar) and branched-chain amino acid catabolites (KIC for leucine) found in this study (Extended Data Fig. 7e). In contrast to reduced uptake of free sugar and amino acids, imidazole propionate (ImP) was increased in Pachón sera (Extended Data Fig. 7e). ImP is a metabolite for which the host does not have a known synthetic pathway, indicating that Pachón cavefish indeed absorbed gut microbiota-mediated metabolites. ImP is also known to inhibit the insulin pathway and fatty acid uptake^79^; thus, high serum ImP in Pachón may discourage blood sugar and fat uptake into tissues^80,81,79,80^. The role of ImP in inhibiting fat anabolism is confusing, yet it was at an undetectable level (Extended Data Fig. 7e).

These results imply that Pachón cavefish are unique in that they do not use sugars and amino acids for immediate oxidation or as energy sources. Previous studies indicate that Pachón cavefish promote anabolism of fatty acids (lipids) and glucose (glycogen)^2,33^, suggesting that these free sugars and amino acids in sera can be stored, possibly used for growth at the early stage.

Short-chain monocarboxylate and ketone-generating pathways may play a major role in the growth of Pachón cavefish, but not in surface fish. This asymmetry could be explained by Pachón’s unique physiology, which may involve primarily storing fatty acids and sugars via anabolism. In general, short-chain monocarboxylates are known to stimulate the release of growth hormone and IGF-1 from the liver in fish^81^, and IGF-1 release from adipose tissues in mice^82^. In addition, the excess of free sugars and amino acids observed in Pachón serum may stimulate somatic growth via mTORC1 pathways^83^. Surface fish are not specialized to store these carbon sources, but use them for energy production via oxidation/catabolism^2,33^, as is typical for fish. Thus, surface fish may not be able to respond to the gut-microbiota-short-chain monocarboxylates-IGF-1 axis to promote growth as Pachón cavefish do. Indeed, surface fish showed slowed growth under a short-chain monocarboxylate supplementation (Extended Data Fig. 7a).

Fasting resilience has been reported in Pachón and Tinaja cavefish, which do not lose a large amount of body weight, but exhibit decreased blood glucose levels after a month of fasting^30,33,34,84^. Fasting Molino cavefish also do not lose weight, but are distinct in maintaining high blood glucose levels after a month of fasting^34^. These results were consistently supported in our results (Fig. 3a-d). Furthermore, in our careful study, Pachón and Tinaja exhibited an evolved association with their gut-enriched pathways that was not found in the Molino population. In the fed condition, many SCFA-directly-producing pathways are positively associated with liver weight, where SCFAs possibly stimulate anabolism and IGF-1, as observed in fish growth (see above). These pathways would be reduced during fasting (*i.e*., during liver weight loss). Pachón cavefish further evolved associations with gut microbiota that provide pathways supporting increased aromatic compound degradation during fasting (Fig. 4a,c). Aromatic compounds can be found in decaying plant material, guano from bats, and/or biofilms^85–87^, and are difficult to break down in the fish digestive system. Thus, carbon and energy supply might be maintained in Pachón cavefish via their gut microbiota processing plant-derived nutrients, which are unique to the Pachón cave population, and not found in the Tinaja or Molino populations. Supplementation of an intermediate compound, 4-hydroxybenzoate (4-HBA), reduced liver weight in the fed condition but not during fasting (the top panel of Extended Data Fig. 12d). It is possible that 4-HBA can act as a signal to the liver about the leanness of the environment, and thus lower liver weight by catabolism. In addition, the positive effect of metabolites of aromatic compound degradation on protecting body weight during fasting (Extended Data Fig. 12e) should be revisited with extended fasting.

It is a mystery why Tinaja cavefish, another starvation-resistant cave population, did not evolve an association with microbes that degrade aromatic compounds, although bacteria that can catabolize them seem to exist in Tinaja cave as well (Fig. 5). A possible explanation is that Tinaja cavefish gained an insatiable appetite through a mutation in the *melanocortin 4 receptor* (*MC4R*), while Pachón cavefish exhibit a comparable level of satiation as surface fish^30^.

Therefore, Tinaja cavefish adaptation may involve ingesting as much as possible to store carbon sources to survive fasting conditions. In contrast, Pachón cavefish may have adapted by recruiting gut microbes that provide novel nutritional output during lean conditions, as well as the capacity to utilize that nutritional output.

Overall, the combination of topological mapping and sparse modeling revealed hidden relationships in noisy, complex data. We find that Pachón cavefish evolved a new coupling with gut microbiota that may contribute to faster growth in resource-limited environments, which may have increased fitness in the cave environment. Our findings clearly indicate the importance of genome by microbiota by diet interactions for the survival of populations and individuals, informing the possible use of microbes to promote health based on individual genotype.

## Materials and Methods

### Fish maintenance and rearing in the lab

*A. mexicanus* surface population fish used were the laboratory-raised descendants of original collections created in Balmorhea Springs State Park in Texas in 1999 by Dr. William R. Jeffery. Cave populations were laboratory-raised descendants originally collected from Cueva de El Pachón (Pachón cavefish) in Tamaulipas, México in 1999 by Dr. William R. Jeffery and in 2013 by Dr. Richard Borowsky, from El Sótano de Molino (Molino cavefish) in Tamaulipas, México in 2013 by Dr. Richard Borowsky, and from the Sótano de la Tinaja (Tinaja cavefish) in San Luís Potosí, México in 1999 by Dr. William R. Jeffery.

Fish (surface fish and three cave populations) were housed in the University of Hawai‘i at Mānoa Aquarium facility with temperatures set at 21 ± 0.5°C for rearing and 25 ± 0.5°C for breeding ^26,88^. Lights were maintained on a 12-h:12-h light:dark cycle ^26,88^. The light intensity was maintained at 30–100 Lux for rearing experiments. Fish husbandry was performed as previously described ^26,88,89^. Fish were raised to adulthood and maintained in standard 42-L tanks in a custom-made water-flow tank system. Adult fish were fed a mixed diet to satiation twice daily starting 1 h after the lights were turned on (Zeitgeber time 1 [ZT1] and ZT9; Zeigler Adult zebrafish irradiated diet, Zeigler Bros, Inc, Gardners, PA; TetraColor Tropical Fish Food Granules, Tetra, Blacksburg, VA, USA; Jumbo Mysis Shrimp, Hikari Sales USA, Inc., Hayward, CA, USA). All fish in the behavioral experiments were between 1.0 and 5 cm in standard length and between 2 and 12 months old. Fish ages were stated in each experiment. All fish care and experimental protocols were approved under IACUC (17-2560) at the University of Hawai‘i at Mānoa.

### Diet, Antibiotics, Fasting, quercetin, BHB and 4-HBA treatments

For the three diet experiment, 1-3 day-old brine shrimp larvae (*Artemia* sp.; Brine Shrimp Direct, Inc., Ogden, UT, USA), powdered spirulina (Pure Spirulina Powder, Kens Fish Inc. Taunton, MA, USA), and bat guano (bat guano fertilizer, Sunleaves Garden Products Inc., Bloomington, IN, USA) were used as diet for juvenile fish. First, hatched larvae of surface fish, Pachón cavefish and Molino cavefish, and their F1s (F1_SP_: a hybrid generation derived from a cross of surface fish female and Pachón cavefish male; F1_SM_: a hybrid generation derived from a cross of surface fish female and Molino cavefish male) were reared in still-water ziplock containers (15.6 × 15.6 × 5.7 cm: Ziplock® containers, S. C. Johnson & Son, Inc., Racine, WI, USA) until 1 month old by feeding live brine shrimp larvae because *A. mexicanus* larval fish cannot ingest difficult diets such as bat guano at the earlier stage (between the 5 days post fertilization when fish start eating and the 1 month old juvenile stage). The ziplock containers were filled with ∼500 mL conditioned water (pH 7.0-7.2 and conductivity: 700-800 µS; conditioned with Seachem Neutral Regulator®, Madison, GA, USA, and Instant Ocean Reef Crystals® Reef Salt, Blacksburg, VA, USA). The conditioned water was fully replaced twice a week. This one-month rearing allowed larvae to develop their gut system sufficiently to take a lean diet, bat guano. After a month of rearing, fish were transferred to the 1-L tanks of Techniplast ZebTEC circulating system (Techniplast USA, West Chester, PA) (∼15 fish per 1L tank, two experimental replicates), and each of the three diet treatments was fed twice a day *ad libtum* every day to surface fish, Pachón cavefish, and Molino cavefish, and their F1s for a month (4 weeks). Fish were aged 2 months old when morphometric measurements, blood samples, and gut tissue samples were collected.

In the ZebTEC circulating system, filtered water was treated with a UV filter before flowing into each fish tank. Yet, the environmental microbes were shared among different fish tanks (*i.e.,* not sterilized condition).

For the antibiotics experiment, we orally supplemented BiFuran+™ (Hikari Sales USA, Inc.), containing Nitrofurazone and Furazolidone, which has been shown to reduce culturable gut microorganisms, compared with other antibiotics (Supplemental Information). BiFuran+ (6.5 % w/v) was mixed with a ground fish diet (Tetra Tropical Color Granules, Tetra U.S., Blacksburg, VA, USA: final 20 % w/v) and with 1 % agar (agar, MilliporeSigma, Burlington, MA, USA) and poured into a 6-cm Petri dish (VWR International, Radnor, Pennsylvania, USA). The control diet consisted of the same composition without adding BiFuran+. 11-month-old age-matched surface fish and Pachón cavefish (∼2.0 cm in standard length, growing) were grouped into 4 fish per still-water container (15.6 × 15.6 × 5.7 cm: Ziplock® containers, S. C. Johnson & Son, Inc.) with ∼500 mL of conditioned water (pH 7.0-7.2 and conductivity: 700-800 µS). The conditioned water was fully replaced twice a week. We fed one to two 3 mm^3^ cubes of the above agar diet per four-fish container every morning (ZT 0:00–3:30) and afternoon (ZT 8:00–12:00). The fish were fed *ad libitum* at each feeding, and the remaining food was removed 1 hour after feeding using a pipette. We fed the antibiotic and control diets for 1 month (4 weeks) before the following assay/analyses.

For the fasting experiment, the one-year-old adult fish (12 months old) were fasted for 1 month (4 weeks), while the control fish were fed with Zeigler Adult zebrafish irradiated diet. The water was circulated in the ZebTEC system, 10% of the whole water was replaced twice per day, and home tanks were cleaned by following our IACUC-approved protocol.

Surface fish and cavefish used in this fasting study were 1-2 years old (adult: 2.5-3.0 cm in the standard length) when we started the fasting trial.

To supplement β-hydroxybutyrate (BHB), we used a commercial fish diet (TetraColor Tropical Granules, Tetra, Blacksburg, VA, USA) mixed with BHB (DL-β-Hydroxybutyric acid sodium salt, MilliporeSigma, St. Louis, MO, USA) at a dosage of 10 mg per gram of body weight (10 mg/body g = 78.7 µmol/body g). In detail, fish generally consume 3% of their body weight (g) per meal. Accordingly, the BHB supplemental diet contained 0.333g/mL of BHB mixed with 0.2 g/mL of the ground Tetra fish diet (20% w/v) in 1% agar. Fish with a body weight of 1 g would consume 30 mg (3%, approximately 30 µL) of this diet, which contained 10 mg of BHB. The control diet consisted of 20% w/v of the Tetra fish diet in 1% agar. After solidification, both the BHB and control diet agar were cut into 3-mm^3^ cubes, and each group of four fish was given 1–2 pieces every morning (ZT 0:00–3:30) and afternoon (ZT 8:00–12:00).

The fish were fed *ad libitum* at each feeding and any remaining food was removed 1 hour after feeding using a pipette. Surface fish and cavefish used in this BHB study were 10-11 months old (small young adults: ∼1.5 cm in standard length, rapidly developing) at the start of the trial.

For the quercetin experiment, we orally supplemented quercetin (Quercetin hydrate, MilliporeSigma, Burlington, MA, USA). First, we prepared a 30 mg/mL quercetin stock solution suspended in dimethyl sulfoxide (DMSO, MilliporeSigma) and stored it at -20°C. Then, 10 µg/ body g of quercetin was supplemented by assuming that fish intake approximately 3% of their body weight, thus 333 µg/ mL (final) quercetin was mixed with a spirulina agar diet (Ultra Intense Spirulina Flake, Ken’s Fish & Pet Supplies, Taunton, MA, USA; final 20 % w/v, with 1 % agar, MilliporeSigma) and poured into a 6-cm Petri dish (VWR International, Radnor, Pennsylvania, USA). The control diet consisted of the same composition with DMSO only. 3- and 8-month-old age-matched surface fish, Pachón and Molino cavefish, were grouped into 4 fish per container (11.5 × 15.5 × 6 cm) with ∼500 mL of conditioned water (pH 7.0-7.2 and conductivity: 700-800 µS) in a water flow system. The sump water in the flow system was replaced twice a week with conditioned water. We fed one to two 3 mm^3^ cubes of the above agar diet per four-fish container every morning (ZT 0:00–3:30) and afternoon (ZT 8:00–12:00). The fish were fed ad libitum at each feeding. We fed the quercetin and control diets for 4 weeks and 5 weeks for 3-month-old and 8-month-old fish, respectively, before the following assay/analyses.

For the 4-hydroxybenzoic acid (4-HBA) supplementation experiment, we used a 2 × 2 design crossing 4-HBA supplementation (with or without 4-HBA) with feeding status (fed vs. fasted). All four diets were prepared in 1% (w/v) agar (MilliporeSigma, Burlington, MA, USA) with 0.5 µM of L-alanine (MilliporeSigma) to induce an appetite. This alanine agar gel was poured into 6-cm Petri dishes (VWR International, Radnor, PA, USA). For the fed conditions, the agar also contained a standard diet (final 20% w/v; Zebrafish Diet, Zeigler Inc., Gardners, PA, USA), whereas the fasting conditions contained agar without the standard diet. In the 4-HBA conditions, 4-HBA (MilliporeSigma) was added at 1.33 mg per gram of agar diet; because fish typically consume approximately 3% of their body weight per feeding, this corresponds to an estimated dose of ∼40 µg per gram of body weight. The corresponding no-4-HBA diets were identical in composition except that 4-HBA was omitted.

Twelve-month-old, age-matched Pachón cavefish were housed in groups of four per container (11.5 × 15.5 × 6 cm) with ∼500 mL of conditioned water (pH 7.0–7.2; conductivity 700–800 µS) in a flow-through system. The sump water was replaced with fresh conditioned water twice a week. Each four-fish container received one to two 3-mm³ cubes of the assigned agar diet every morning (ZT 0:00–3:30) and every afternoon (ZT 8:00–12:00), and fish were allowed to feed *ad libitum* at each feeding. Fish were maintained on their assigned diets for 4 weeks before the subsequent assays and analyses.

### Measurement of body

Fish were anesthetized with ice-cold conditioned water (pH: 7.0; conductivity: 700 µS) for 3-5 min, and weight was measured after pictures were taken with a standard camera (Pentax K-1 DSLR with 35-70 mm zoom lens, Ricoh, Tokyo, Japan). The standard body length and body depth were measured using ImageJ software^90^.

### Blood ketone and glucose measurements

All blood samples were collected 2–3 h after feeding. Fish were then deeply anesthetized in ice-cold water, and blood was collected from the tail artery. Blood ketone and glucose levels were measured using either the Keto-Mojo GK+ (Keto-Mojo, Napa, California, USA) or the Abbott Precision Xtra blood glucose and ketone monitoring systems according to the manufacturers’ instructions. The readings of Abbott Precision Xtra were standardized by the reads of the same blood sample with Keto-Mojo GK+. Both reads from the Abbott and Keto-Mojo meters were highly linearly correlated (R^2^ = 0.93, P = 0.000103 and R^2^ = 0.74, P = 0.00565 for glucose and ketone reads, respectively; N = 8).

### Serum metabolomics for 300 metabolites

All blood samples were collected 2–3 hours after feeding. Fish were then deeply anesthetized in ice-cold water, and blood was collected from the tail artery by using a P10 pipet. Typically, 1.5 – 2.0 µL of blood were recovered. Recovered blood was immediately mixed with 10 µL of 5 mg/ml Heparin (Heparin sodium salt, VWR International, Radnor, PA, USA) in ice-cold phosphate-buffered saline (PBS). The blood solution was then centrifuged with a mini-spinner (approx. 2,000 × g, BenchMate C8-M, Oxford Lab Products, San Diego, CA, USA) for 10 min, and the supernatant serum was recovered in a new 1.5 mL tube and immediately frozen in liquid nitrogen. These samples were stored at -80 °C until analyzed. Before analysis, four fish sera were pooled into one sample (∼12 µL × 4 fish = ∼48 µL per sample) to obtain the required volume for analysis. These sera group was then processed for non-targeted metabolomics at the University of Hawaii Cancer Center (Q300 assay)^91^. Briefly, 20 µL samples were diluted with the internal standard solution, centrifuged, and then supernatant was derivatized with 200 mM 3-nitrophenylhydrazine (3-NPH) and 96 mM N-(3-(dimethylamino)propyl)-N′-ethylcarbodiimide (EDC), which target carboxyl residues, thereby facilitating detection of organic acids and amino acids. Detection of metabolites was performed using ultra-performance liquid chromatography coupled to tandem mass spectrometry (UPLC-MS/MS), which detected 135 metabolites.

### Tissue dissection and measurements

After blood collection, under anesthesia with ice-cold conditioned water, fish heads were decapitated with decontaminated dissection scissors (Fine Science Tools Inc, Foster City, CA, USA, and RNase *AWAY*™ Decontamination Reagent, Thermo Fisher Scientific Inc., Waltham, MA, USA), and their guts and livers were carefully extracted from their body cavity without contamination from the body surface or anywhere. The gut tissues were then transferred to a new sterile 6 cm dish with a sterile PBS droplet by decontaminated dissection forceps (Dumont No.2 and No.3; Fine Science Tools Inc; RNase *AWAY*™). The fish guts were then carefully dissected by detaching livers, stomachs, extrinsic nerves, gut mesentery, and fat tissue under sterile conditions. Liver weight was measured on the scale, recovered into a 1.5 mL tube after rinsing with PBS, and quickly frozen in liquid nitrogen. Then, pyloric caeca, mid- and hindguts were photographed under a binocular microscope attached with a colored charge-coupled device (CCD) camera (Olympus SZX-10 Stereo Zoom microscope, EVIDENT, Tokyo, Japan; Infinity 3-1UR Digital Color Camera, Teledyne Luminera, Ottawa, Canada). The number of pyloric caeca was counted under binocular microscopes. The total gut length, hindgut length, and mid-/hindgut widths were measured on the saved TIFF images using ImageJ or Fiji software^90,92^. Each of mid-/hindgut widths was the average of three different measurements within either mid- or hindguts to standardize the variations within each of the gut segments. The midgut with pyloric caeca and hindgut were then separated, and stocked into the different tubes, by quickly frozen in liquid nitrogen, and stored at -80°C until the set of samples was collected.

### Environmental samples

For the water samples from Pachón and Tinaja cave, approximately 500 mL of water was filtered with the 0.2 µm pore size Whatman® nylon filter discs (25 mm diameter; MilliporeSigma, Burlington, MA) attached to a manual vacuum pumping unit. After filtering the water, the 0.2 µm filter was carefully folded and stored in a 2.0 mL screw-capped tube (VWR International, Radnor, PA), filled with RNA*later*™ Stabilization Solution (ThermoFisher Scientific, Waltham, MA), stored at room temperature for < 1 week, and stored at -80°C until 16S rRNA gene sequencing.

### 16S rRNA gene sequencing and analyses

Gut samples stored at -80 °C were then processed in the standard 16S rRNA gene V4 protocol for library preparation. Briefly, each gut sample (midgut or hindgut. Approx. 10-50 mg each per individual fish) was added to five ceramic 1.8 mm beads and 80 µL of sterile phosphate-buffered saline (PBS) and homogenized using an OMNI Bead Ruptor 12 (OMNI International, Kennesaw, GA, USA) spun at speed 2.10 for 30 seconds. DNA was extracted on a KingFisher™ Flex™ System (Thermo Fisher Scientific, Waltham, MA, USA) with a MagAttract PowerSoil KF Kit (Qiagen, Hilden, Germany), or manually with DNeasy PowerSoil Pro Kit (Qiagen) following the manufacturer’s protocols. The V4 region of the bacterial 16S rRNA gene was amplified in a single PCR using ribosomal RNA gene primers containing a 12 base pair Golay-indexed code for demultiplexing, (515F: GTGYCAGCMGCCGCGGTAA; 806R: GGACTACNVGGGTWTCTAAT ^93^)

PCR reactions were performed with the Platinum™ II Hot-Start PCR Master Mix (Thermo Fisher Scientific) with 2,000 nM of a blocking primer (see below) under the following conditions: 94°C for 2 min, followed by 35 cycles of 94°C for 15 s, 55°C for 15 s, 68°C for 7 s and a final extension for 68°C for 3 min. A blocking primer with C3 Spacer phosphoramidite (18S Rev BLPr: GTATCTATCGTCTTCGACCTCC/3SpC3/; Integrated DNA Technologies, Inc., Coralville, Iowa, USA) was introduced to block the amplification of the fish 18S ribosomal RNA gene (∼600 bp while 16S rRNA gene is ∼470 bp). The above PCR parameters, including selecting PCR enzymes with the blocking primer, were optimized by the M-GAL metagenomics core at the University of Hawai‘i at Mānoa.

The PCR products were cleaned and normalized using the Just-a-plate kit (Charm Biotech, San Diego, CA, USA). High-throughput sequencing (HTS) was performed with Illumina MiSeq and V3 300 bp paired-end kits (Illumina, Inc., San Diego, CA, USA) at ASGPB core at the University of Hawai‘i at Mānoa. The sequences were demultiplexed into FASTQ files. Read filtering, denoising, and contig generation were performed by DADA2 by dropping reads shorter than 20 bp. Paired reads are merged (= contig) if the overlap is at least 20 bp with a maximum 1 bp mismatch^94^. The contigs generated by DADA2 were subsequently processed using MOTHUR^95^, which aligned and annotated these contigs using the SILVA v138 database^96^, and generated a count matrix for Amplicon Sequence Variants (ASVs). Chimeric contigs were removed with VSEARCH^97^. Then, ASVs with a total abundance below 2 were discarded. Finally, the LULU algorithm^98^ with the following settings was used to remove spurious ASVs from PCR errors and/or merge ASVs from multiplicated 16S rRNA operons per genome. Our LULU setting was that a daughter ASV is merged with its parent if (i) they share at least 97% similarity, (ii) the daughter ASV abundance < parent ASV abundance on average, and (iii) the relative co-occurrence (proportion of time the daughter is present when the parent is present) must be 1.0.

We introduced negative controls (PBS only) and positive controls with a known bacterial community structure (ZymoBIOMICS Microbial Community Standard, Zymo Research, Irvine, CA, USA) at the step of DNA extraction and at the PCR (ZymoBIOMICS Microbial Community DNA Standard) to monitor potential taxonomic biases introduced during the benchworks. Ten replicate samples were also used to check the reproducibility of the resulting sequencing data.

Our result indicated that there was no significant bias on taxonomies in our dataset.

### Topological Data Analysis (Topological mapping) and PICRUSt2 estimates

The raw count data were filtered with taxa whose total counts are more than 250, and samples whose total counts are more than 500. DyNeuSR package^99^ was used to generate mapper graphs and used the second principal component (PC2) of principal component analysis as the filter function for Extended Data Fig. 3a,b. The filter functions were then the host trait scores in the following analyses (*e.g.,* body standard length and gut length). Mapper graphs using the host trait of interest as the filter function were used for the downstream sparse modeling analysis described below.

ASV abundances were normalized using GMPR size factors^100^ to explore associations between host trait scores and ASV counts. To assess host’s trait–associated gut-enriched pathways, we applied GMPR normalization to ASV abundances before PICRUSt2 pathway inference – although this may appear to differ from PICRUSt2 recommendations, it is consistent with the PICRUSt2 usage guidelines as described below:

GMPR^100^ was formulated and benchmarked for zero-inflated, non-negative integer ASV count tables, which our ASV table (2,335 features, 92.5% zeros) exemplifies, and the large number of quasi-independent features gives the majority-invariance assumption a broad basis. We did not adopt GMPR normalization for pathway abundances, which are non-integer derived quantities coupled to one another by functional redundancy among taxa; across only 397 such features, fewer effectively independent features remain to support the assumption. GMPR estimates each pairwise ratio as a median over features non-zero in both samples, and this shared domain covers a median of 51.4% of a sample pair’s abundance at the ASV level versus 99.8% after the PICRUSt2 aggregation (13.3% versus 91.0% by feature count). Consistent with the assumption being less well supported after the ASV → pathway aggregation, applying GMPR to pathway abundances removed each sample’s own profile level almost completely (between-sample standard deviation [s.d.] of each sample’s median log_2_ level, across the 179 pathways fully prevalent in both pipelines: 0.150), whereas applying GMPR to the ASV table left that level intact (s.d. = 1.556).

Lastly, PICRUSt2 itself divides the ASV table by predicted 16S copy number before aggregation, providing in-tool precedent for ASV-level normalization preceding aggregation into pathways.

Per-sample depth correction prior to PICRUSt2 is more generally a sanctioned category of operation: the PICRUSt2 tutorial explicitly permits rarefied input, and GMPR can be viewed as a non-destructive, deterministic alternative to rarefaction within that category.

### Screening of ASVs/ metabolic pathways using combined topological data analysis and lasso regression (sparse modeling)

The following analysis framework is intended as an exploratory screening for candidate taxa, where the selected candidate taxa would be applied for experimental validations; therefore, this framework does not test hypotheses. Also, analyses were performed on each population or F_1_ group individually and across the treatments (*i.e*., combined brine shrimp, spirulina, and bat guano data; combined fed and fasted conditions). This approach was chosen to widen the data validation (across treatments) but kept the dataset simple (each population individually). The gut microbiome exhibits functional redundancy, a concept originating from community ecology^101,102^, whereby distinct microbial taxa across host individuals can contribute equivalently to a given host trait^103,104^. This redundancy renders standard regression approaches unstable across the full sample space, as different taxa carry the trait-associated signal in different individuals.

To address this, we applied TDA Mapper with the host trait score as the filter function, partitioning samples into locally coherent subsets along a gradient from low to high trait values, in which a single community type dominates. This stratification reduces the confounding influence of community-level redundancy on variable selection and improves the interpretability of identified taxa as trait-associated features.

LASSO regression was subsequently applied either to the full sample set or to the Mapper-derived sample subset depending on model fit. The detailed sparse modeling pipeline, including repeated LASSO with varying L1 ratios, stability-based filtering, univariate validation, and recovery of collinear taxa, is described in Extended Data Fig. 3. Sparse modeling was implemented using two complementary regularization approaches: ElasticNet regression (Python, scikit-learn; L1 ratio range: 0.1–1.0, 10-fold cross-validation × 10 repeats) and LASSO-GLM (MATLAB lassoglm; L1 ratio range: 0.1–1.0, 10-fold cross-validation × 10 repeats).

ElasticNet combines L1 and L2 penalties, facilitating the joint selection of correlated taxa, whereas LASSO applies an L1 penalty alone, tending to select a single representative from a group of correlated features. ASVs selected in ≥80% of repeats within each L1 ratio were retained as consistently selected predictors. For each L1 ratio, retained ASVs were refit using ordinary least squares regression (Python: statsmodels OLS; MATLAB: fitglm, Gaussian family) to obtain unbiased coefficient estimates and model fit statistics (Extended Data Fig. 3e, Step1). Predictors identified by either or both implementations were pooled, and this combined set was carried forward for downstream filtering steps (Extended Data Fig. 3e, Step2).

The biological validity of candidate taxa identified through this pipeline was confirmed through independent experimental perturbation (antibiotics and quercetin treatment: Extended Data Fig. 4, Extended Data Fig. 5, Fig.2c), providing empirical justification for the analytical framework independent of its statistical assumptions.

### Statistical analysis

Regarding the power analysis, we designed our experiments based on three-way repeated-measures ANOVA with a moderate effect size (f = 0.25), alpha-error probability of 0.05, and power of 0.80, and the number of groups was eight (surface fish vs. cavefish × non-treated vs. treated × pre-treatment vs. post-treatment). G*Power software^105,106^ estimated that the sample size needed for this experiment was nine per group. We thus aimed to use at least 12 fish in each group for all experiments in this study.

For statistical comparisons of our data, we performed tests including Student’s *t*-test and two- or three-way generalized linear model analyses to compare surface and cavefish, treatment and non-treatment, and pre-treatment and post-treatment. We applied the linear model to non-processed data, generalized linear model (Poisson or negative binomial family) to discrete data, and generalized linear model (Gamma family) to continuous data. In repeated measures, we included the fish IDs as random effects. Tank was not included as a random effect because, with only two or three replicate tanks per treatment, the among-tank variance could not be reliably estimated and the model failed to converge (singular fit). Holm’s post hoc correction was used to determine which contrasts were significant^107^.

We performed the aforementioned calculations in R version 4.3.2 using the RStudio 2024.04.2 user interface (packages: car, lme4, glmmTMB, and lmerTest)^108–111^, and all raw data are available in Supplemental Data 2. Statistical scores are available in Supplementary Data 3 or the figure legends.

## Supporting information

Supplemental data 2 (raw data)

Supplemental data 3 (statistics scores)

Supplementary Info (Method and Discussion)

## Acknowledgment

Dr. Karen Gillemin for suggesting an analysis strategy for an affected microbiome under antibiotic treatment. We are grateful to J Choi, L Lu, J Nguyen, VFL Fernandes, K Lactaoen, M Worsham, H Hernandez, N Doeden, M Ito, A Martinez, D Mones, A Velazquez, A Jee, B Weise, B Espejo, C Tong, D Inouye, D Burbano, E Stevenson, H Hardy, I Matsuyoshi, I Riethbrock, J Southworth, J Boggs, J Cashon, J Hwang, J Tong, K Politan, K Cousens, K Fukumoto, K Wong, M Rai, M Hasegawa, M Quadri, M Pugh, N Choi, R Chung, S Do, S Singh, S Blattner, T Okamura, TM Ohta, V Garcia-Tobar, and H Yoshizawa for fish care assistance and useful scientific inputs.

## Contributions

MI: designed the quercetin and 4-HBA experiments, performed the experiments, developed a topological data analysis plus sparse-modeling pipeline, reanalyzed all 16S rRNA gene data, interpreted the data, and edited the manuscript.

CV: performed the 3-diet and fasting experiment and analyses, edited the manuscript.

AGH: co-designed and performed the fasting experiment and initial 16S rRNA analyses, edited the manuscript.

XBG: performed the quercetin experiment and analysis, interpreted data, and edited the manuscript.

JK: performed the antibiotic experiment and culturable colony analysis and interpreted the data.

ED: performed the antibiotic experiment with BIFRAN+, performed all measurements and dissection, interpreted data, and edited the manuscript.

AT: performed the BHB experiment and analysis.

LJTR: co-designed and performed the fasting experiment and analysis, edited the manuscript.

RDP: performed water sample DNA/ 16S rRNA gene analysis and edited the manuscript.

KC: performed the 3-diet 16S rRNA experiment, its initial analysis, and interpreted the data.

ACB: performed the 3-diet 16S rRNA experiment, its initial analysis, and interpreted the data.

NB: performed the 3-diet 16S rRNA experiment and its initial analysis, interpreted the data, and edited the manuscript.

RBR: co-designed the 3-diet 16S rRNA experiment and its initial analysis, interpreted the data, and edited the manuscript.

FN: developed topological data analysis methods, applied them in the analysis, supported the pipeline development, and edited the manuscript.

MY: designed the overall project and experiments, performed the experiments, analyzed the data, interpreted the results, wrote the initial draft, and edited the manuscript.

---

**Extended Data Fig. 1.**
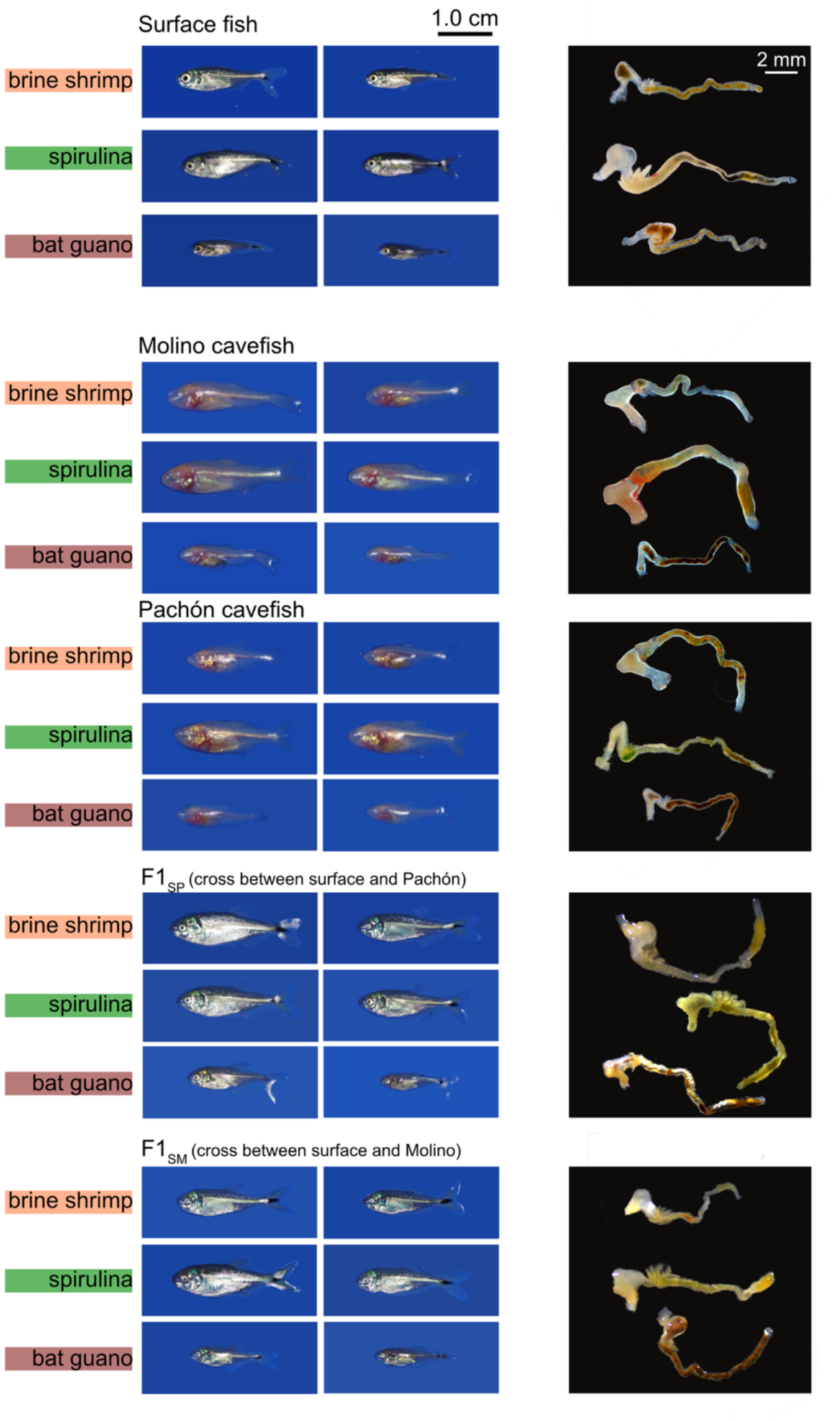
Fish body and gut pictures at the end of 1 month of treatment with the diet. Two representative individuals of fish in each category (populations and diet) are shown on the left panel, and representative gut pictures are shown in the right panels. Cavefish and F_1_s grew well under a spirulina diet.

**Extended Data Fig. 2.**
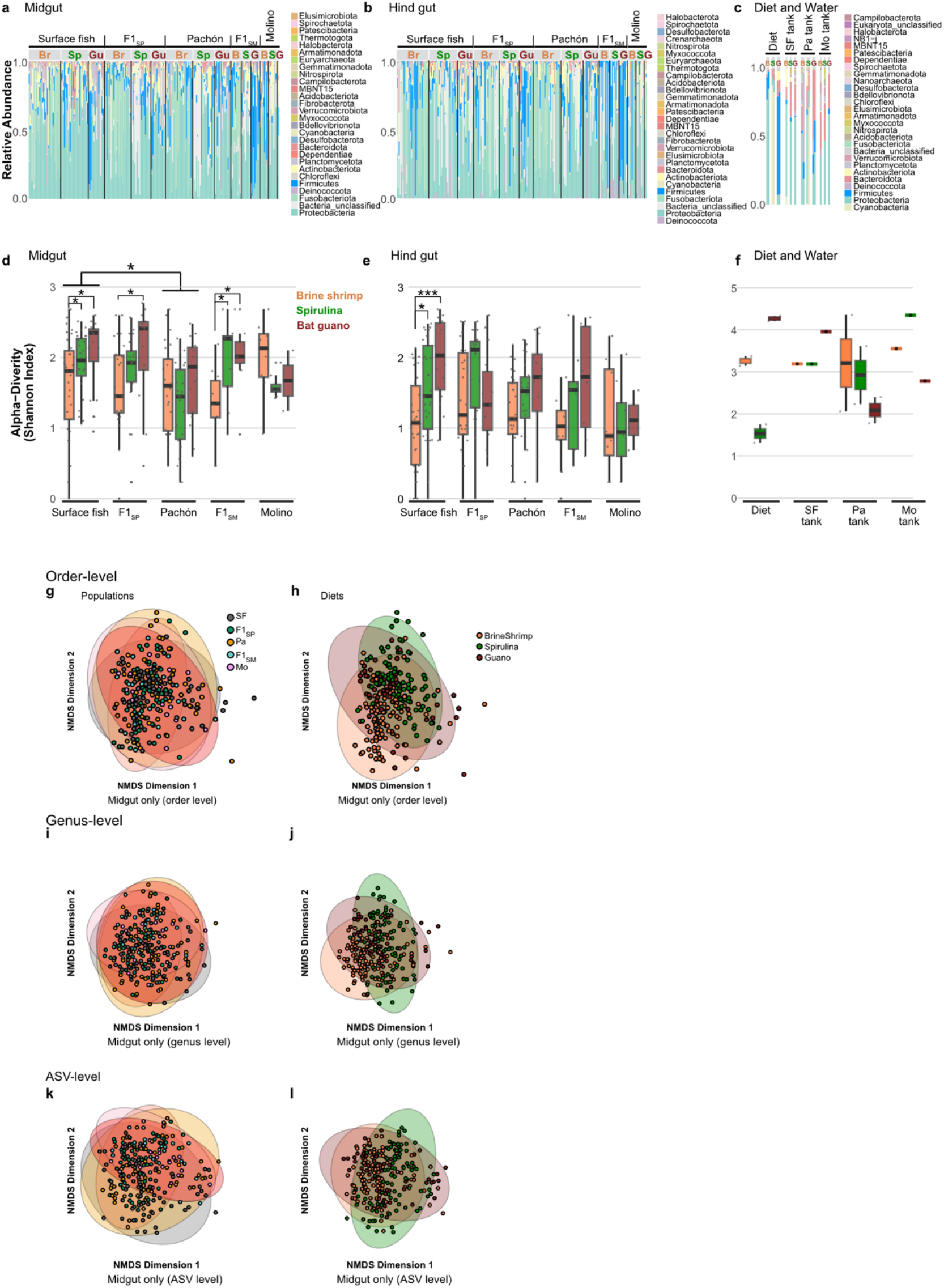
Gut microbiota in the midgut exhibited more alpha diversity than that of the hindgut. The gut microbiota profiles are distinct from those of the diets or aquarium water. **a-c**, Relative abundance of gut microbiota from midgut (**a**), hindgut (**b**) from surface fish (SF), F_1_ hybrid between surface fish female × Pachón male (F1_SP_), Pachón (Pa), F_1_ hybrid between surface fish female × Molino male (F1_SM_), and Molino cavefish (Mo), and the relative abundance of the microbiota of the diet and housing water (**c**). **d-f**, Alpha diversity (Shannon-Weaver Index, ASV level) of the microbiota in the midgut (**d**) and hindgut of SF, F1_SP_, Pa, F1_SM_, and Mo (**e**), and relative abundance of the microbiota of the diet and housing water (**f**). **g-l**, Beta diversity of gut microbiota from midgut was colored with population (**g**) and diet (**h**) in the order level, genus level (**i,j**) and ASV-level (**k,l**). *: P < 0.05, **: P < 0.01, ***: P < 0.001 after post-hoc Holm’s correction

**Extended Data Fig. 3.**
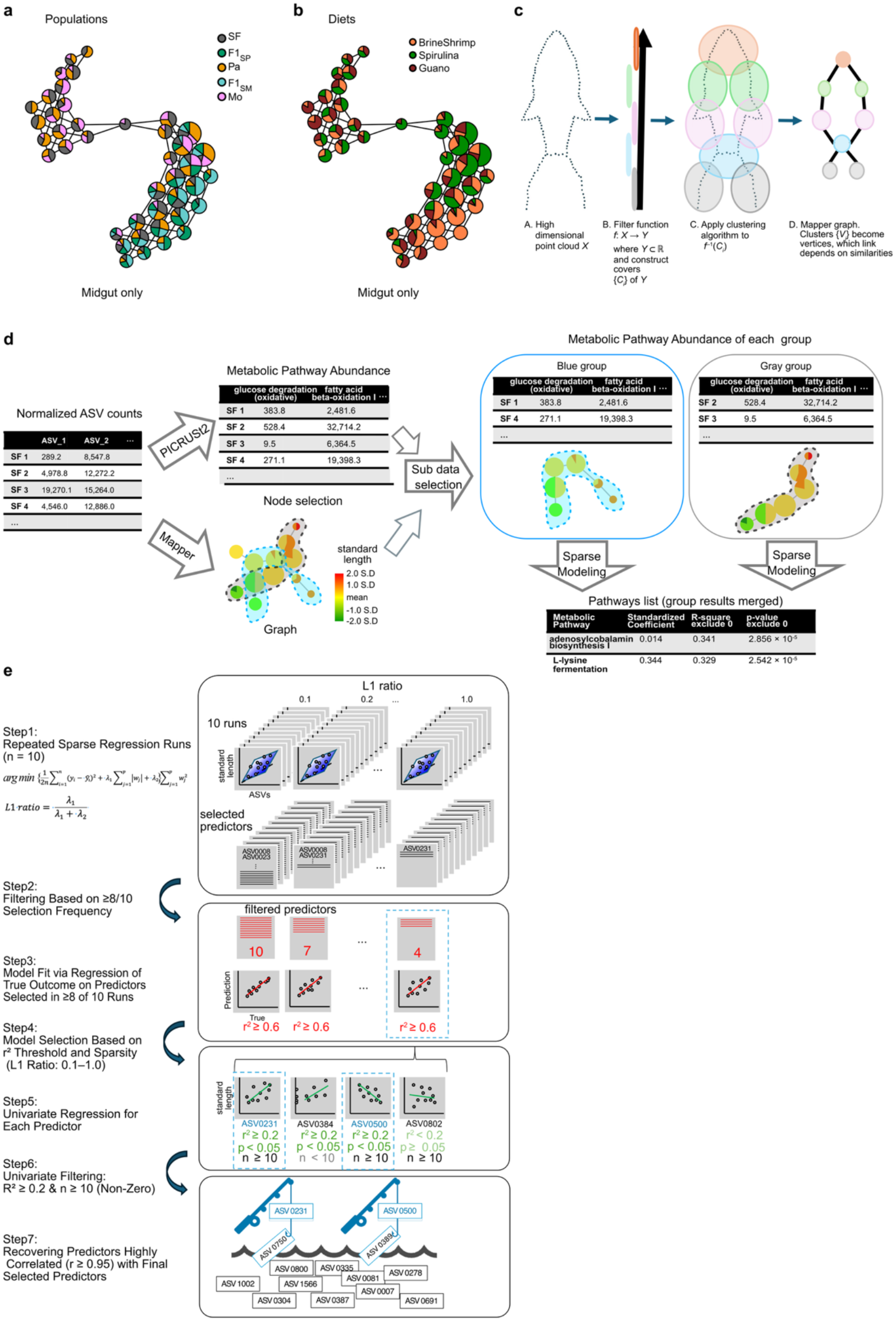
Topological Data Analysis (TDA) Mapper guides the filtering of sample sets, followed by sparse modeling, revealing hidden couplings between morphological traits and gut microorganisms. **a, b**, Topological mapper graph colored by population (**a**) and diets (**b**). A Pie chart represents a cluster consisting of multiple fish individuals whose populations (a) or diet treatments (b) are shown. The principal component (PC) 2 was used for the 1-D filter function, and DBSCAN was used for clustering. **c**, A schematic of the topological mapper visualization process. High dimensional raw data (A) are projected to a 1-D quantitative score (*i.e.,* filter function). The 1-D quantitative score (*e.g.,* PC1 in PCA, or body standard length) is then binned with some overlap (B), followed by clustering the fish individuals based on the similarity of gut microorganism compositions within each bin (C), and then each cluster is linked depending on the similarity of gut microbiota compositions (D: graph structure). The topological mapper graphs captured some separation in populations and diet treatments but still exhibit complex structures. **d**, Workflow for screening gut microbial taxa and functions coupled to host traits beyond conventional ordination. ASV counts from 16S rRNA gene sequencing (mothur-based C-MAIKI pipeline^112^) were normalized by the Geometric Mean of Pairwise Ratios (GMPR)^100^. Normalized counts were then used in parallel for (i) PICRUSt2^59^ prediction of metabolic pathway abundance and (ii) Mapper, with the host trait of interest as the 1-D filter function (body standard length in this example). GMPR was applied before PICRUSt2 because PICRUSt2 expects depth-adjusted input; unlike rarefying, GMPR removes the technical scale imposed by sequencing depth while retaining biologically meaningful differences in overall abundance and discarding no reads. Sample subsets were defined by traversing the Mapper graph from low- to high-trait clusters along their links (blue and gray groups). Within each subset, ASV or pathway abundance was regressed against the trait by sparse modeling (**e**). **e**, Sparse-modeling pipeline used in (**d**), designed for zero-inflated, small-n datasets. *Step 1*, elastic-net regression is run ten times at L1 ratios of 0.1, 0.2, … 1.0. *Step 2*, predictors (ASVs or pathways) selected in ≥ 8 of 10 runs are retained. *Step 3*, retained predictors are refit by ordinary multiple regression. *Step 4*, among models with r² ≥ 0.6 for observed versus predicted trait scores, the one with the fewest predictors is selected (here, L1 ratio = 1.0). *Steps 5–6*, each selected predictor is regressed univariately against the trait and retained only if *r*² ≥ 0.2, P < 0.05, and the number of non-zero samples ≥ 10 (blue dashed boxes). *Step 7*, predictors dropped during regularization owing to collinearity are recovered if their correlation with a retained predictor is *r* ≥ 0.95. These processes are to screen the candidate ASVs or pathways; therefore, we skipped statistical tests intentionally (see Materials and Methods)

**Extended Data Fig. 4.**
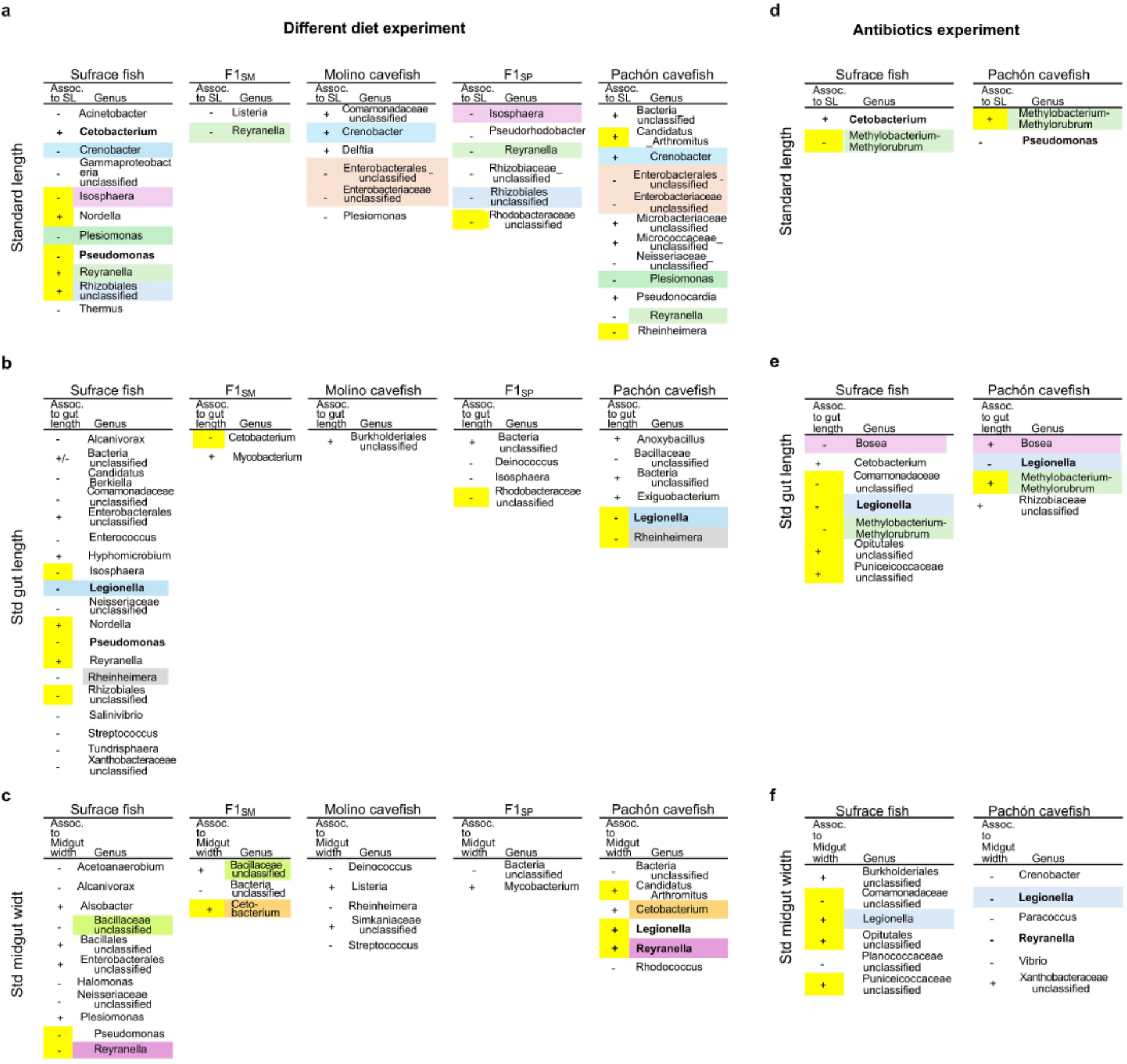
ASVs that were associated with the host traits were distinct among populations, and consistently emerged after the antibiotic intervention. **a-c,** Microorganisms’ taxa coupled to standard length (**a**), total gut length (**b**), and midgut width (**c**), which were detected in the midguts of surface fish, Molino and Pachón cavefish, and F_1_ hybrids in the three-diet experiment. To gain the widest variation of morphological measurements for the sparse modeling regression, all fish individuals treated with three distinct diets per population were used for the sparse modeling regression (*c.f*., the blue hatched lines in Extended Data Fig. 7c; note that the x-axis is for ASV counts, instead of pathway abundances). **d-f**, Microorganisms’ taxa coupled to standard length (**d**), total gut length (**e**), and midgut width (**f**) in the antibiotic intervention experiment. All fish individuals treated with the control and antibiotics per population were used for the sparse modeling regression. Color-shaded genus names are shared across populations. Yellow shadings with + or – signs are the shared genus among standard length, total gut length, and/or midgut width. Boldfaced genus names indicate the shared taxa between the three-diet and antibiotic intervention experiments.

**Extended Data Fig. 5.**
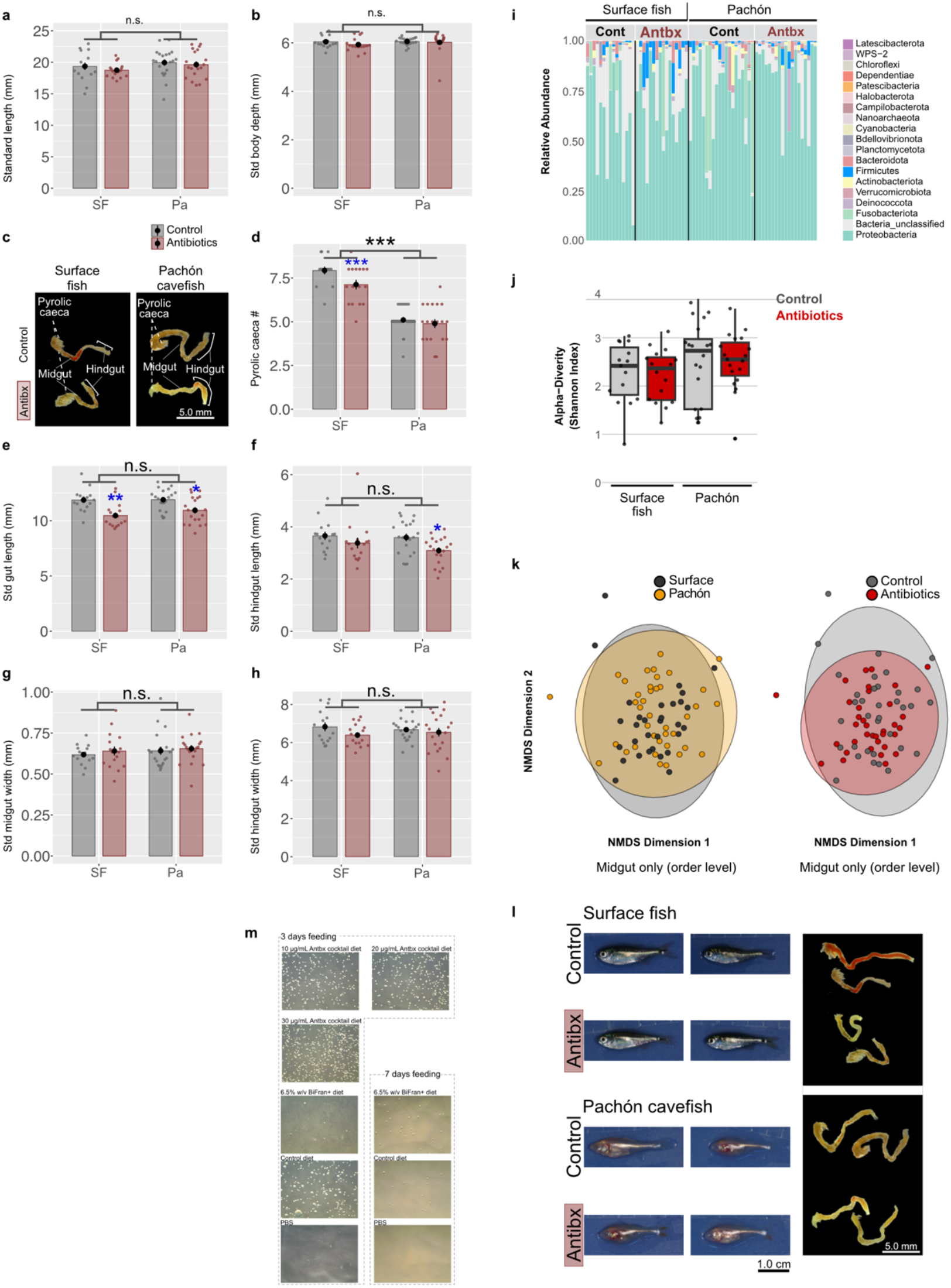
Antibiotic effects on body morphology, gut morphologies and gut microbiota. **a, b**, The body standard length (**a**) and body depth standardized with the body standard length (**b**) of surface fish and Pachón cavefish under antibiotic supplementation. **c,** Gut images of surface fish (left) and Pachón cavefish (right) in the control (tops) and antibiotics treatments (bottom). **d-h**, Number of pyloric caeca (**d**), standardized total gut length (**e**), standardized hindgut length (**f**), standardized midgut width (**g**) and standardized hindgut width (**h**) of surface fish and Pachón cavefish. **i**, Relative abundance of gut microbiota in the midguts of surface fish and Pachón cavefish. In the antibiotic treatment, firmicute seems to increase in relative abundance in both surface fish and Pachón cavefish. **j**, Alpha diversity (Shannon-Weaver Index) of gut microbiota from midgut. **k**, 2-D plot of non-metric multidimensional scaling (NMDS) colored by population (left) and experimental condition (right). The clusters of populations and experimental conditions (control and Antibiotics) were largely overlapping. **l**, Fish body and gut pictures after 1 month of antibiotics diet feeding. Two representative individuals of fish in each category (populations and antibiotics: Antibx) are shown on the left panel, and representative gut pictures are shown in the right panels. Body growth and gut morphologies showed no detectable change between non-treated and Antibx treated. **m**, Culturable colonies on the BHI plates from the gut homogenates of surface fish, treated with food mixture of antibiotics cocktail (vancomycin, metronidazole, cefoxitin and gentamycin) or BiFran+ for 3 or 7 days.*: P < 0.05, **: P < 0.01, ***: P < 0.001 after post-hoc Holm’s correction

**Extended Data Fig. 6.**
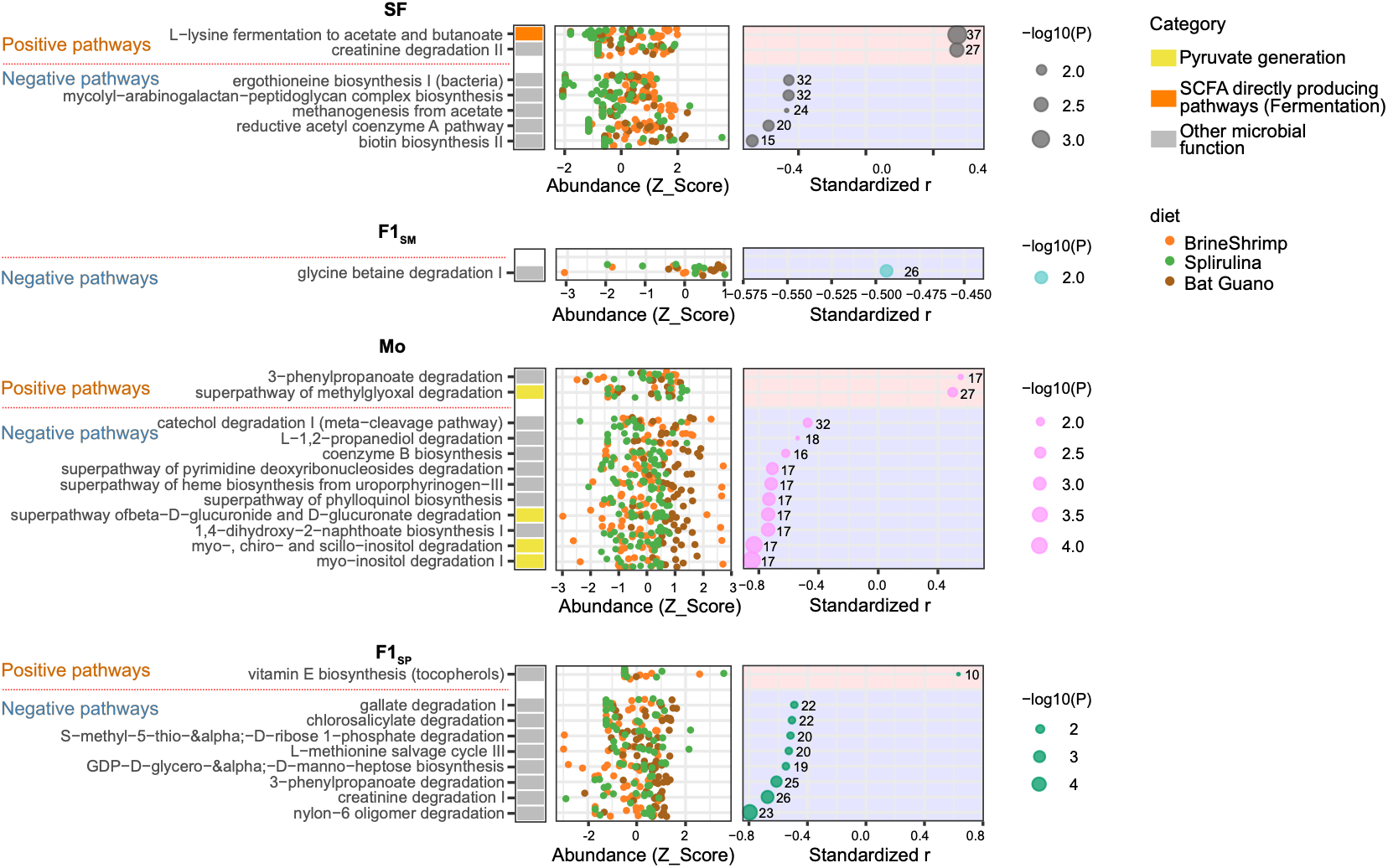
Significant associations/coupling between body standard length and their gut-enriched pathways in the ancestor-like lineage, surface fish, Molino cavefish and F1 hybrids (F1SP and F1SM). The gut-enriched pathways were aligned according to the levels of Pearson’s correlation coefficients (*r*: univariate regression, far right panel). All fish individuals treated with three distinct diets per population were used for the sparse modeling regression. Circle sizes represent the -log_10_(P-values of the correlation coefficient). The z-score of each pathway’s abundance (PiCRUSt2 estimated) is shown in the middle column. Colors of the dots are as follows—orange: brine shrimp, green: spirulina, and brown: bat guano. These gut-enriched pathways were also grouped as “Pyruvate generation”, “SCFA direct producing pathway” and “Other microbial function”—custom-made categories based on the MetaCyc ontology (far left panel; Supplementary Table 1).

**Extended Data Fig. 7.**
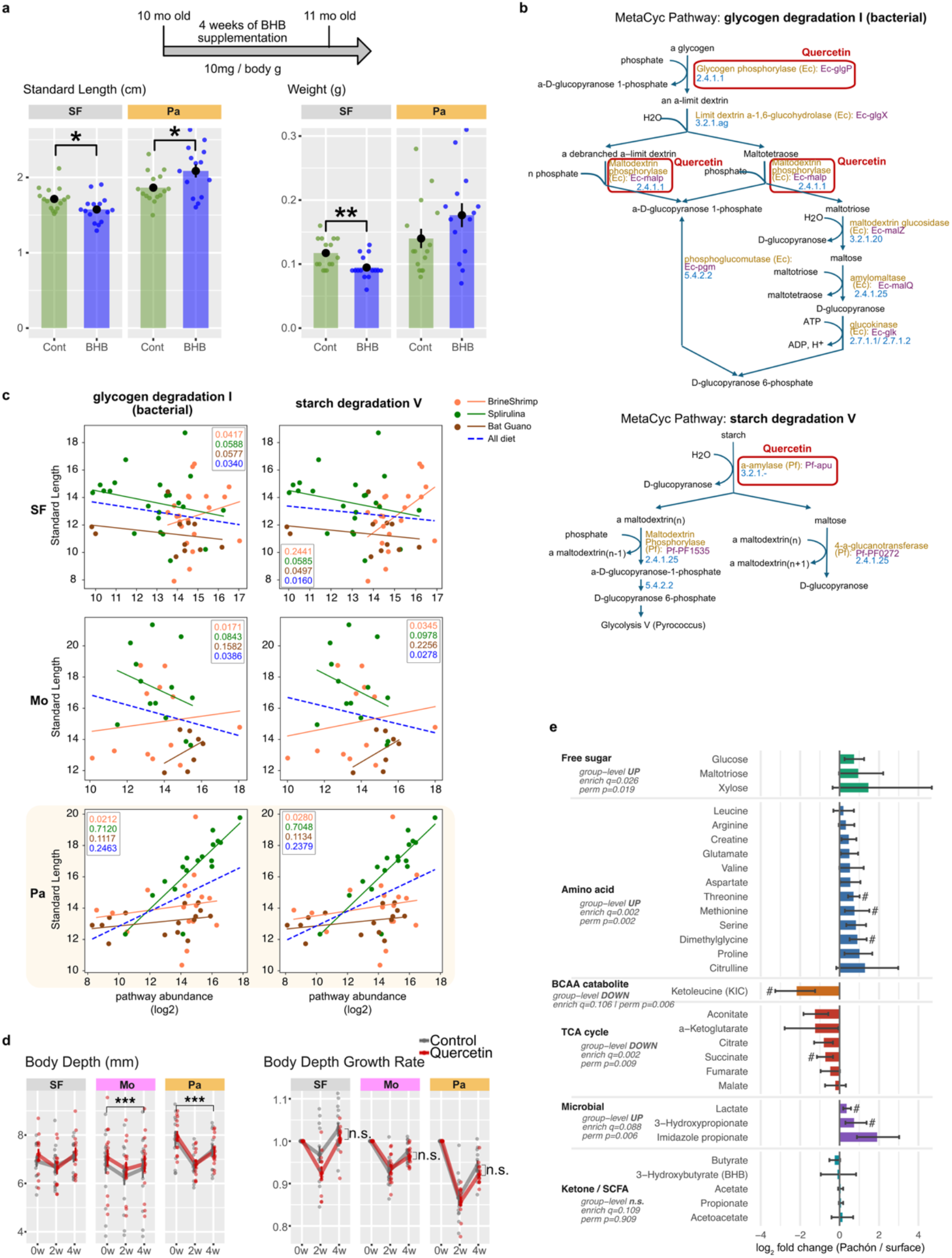
Glycogen degradation I and starch degradation V, associated with Pachón cavefish growth, would promote Pachón cavefish growth via ketone bodies. **a**, After 4 weeks of supplementing beta-hydroxybutyrate (BHB) at 10-month-old fish (∼1.5 cm in standard length), the body standard length was increased in Pachón cavefish (Pa, left panel) while it was reduced in surface fish (SF). Please note that these cohorts were much smaller than the 8-month-old fish used in Fig 2c, and they were rapidly developing. Body weight was reduced in SF, but there was no detectable difference between the control and BHB-treated Pachón cavefish (right panel). **b**, Quercetin inhibits glucokinase (circled in red at top panel) in the glycogen degradation I pathway and α-amylase in the starch degradation V pathway (circled in red). **c**, Scatter plot of body standard length vs. abundances of each gut-enriched pathway in surface fish, Pachón cavefish, and Molino cavefish. Different dot colors represent individual fish under different diet treatments. The colored solid lines show regression lines of each diet treatment; the blue dotted line indicates the regression of all samples under the three diet treatments. Numbers in the inset represent *r²* values. **d**, The body depth change under quercetin supplementation (*c.f.,* Fig. 2c). No detectable changes in growth rate. **e**, Log_2_ fold change in 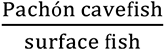 of the serum concentration of each metabolite. Each metabolite was grouped into surface fish free sugars, free amino acids, a catabolite of branch-chain amino acid (BCAA), TCA cycle-related metabolites, microorganism-contributed metabolites, or ketone/short-chain fatty acids. Error bars indicate 95% confidence intervals calculated by 10,000-time permutation simulations. #: The Benjamini-Hochberg false discovery rate (BH-FDR) > 0.10. Group-level up- or down-regulations were also statistically tested and shown. *: P < 0.05, **: P < 0.01, ***: P < 0.001 after post-hoc Holm’s correction.

**Extended Data Fig. 8.**
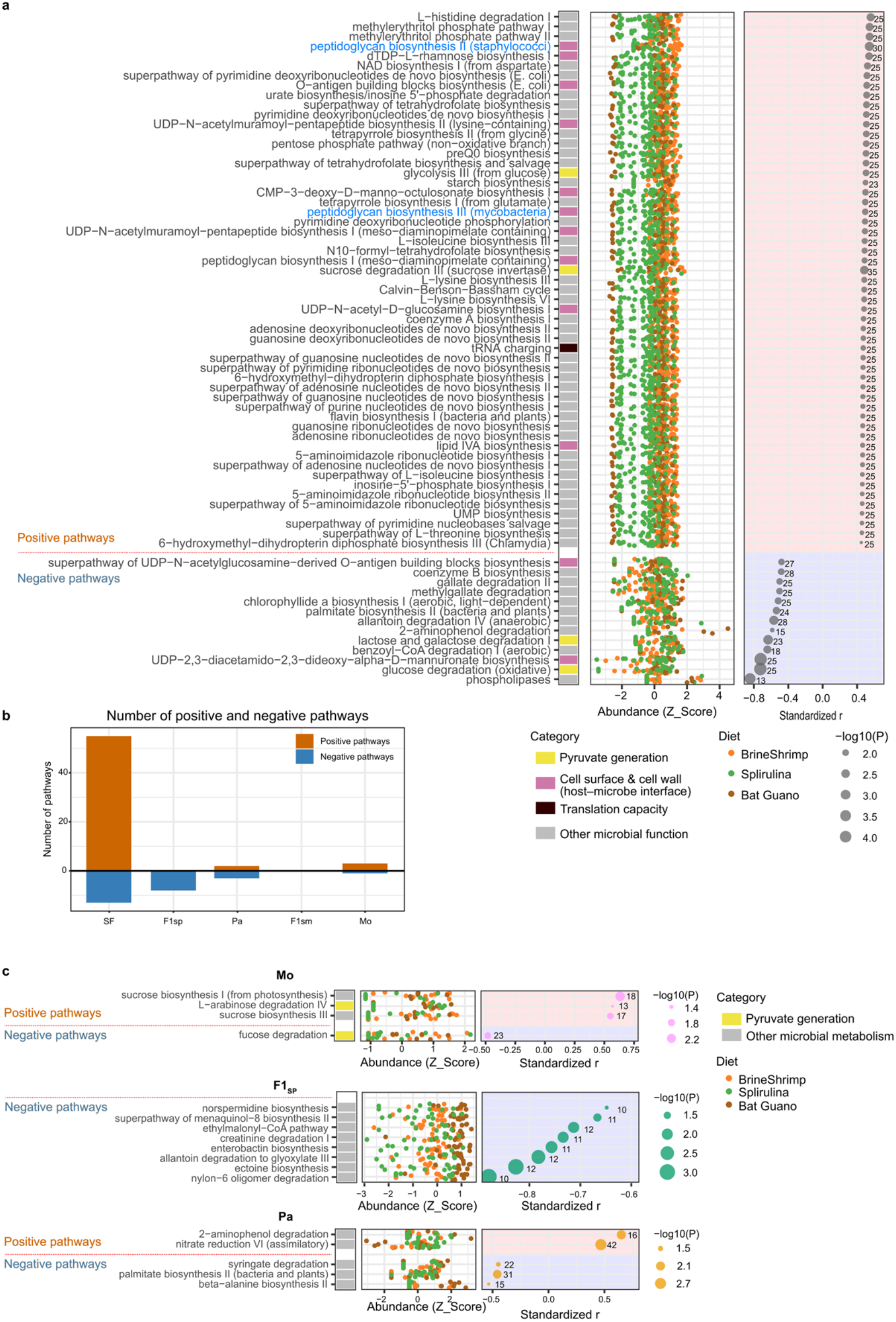
The gut length-associated gut microorganisms and their enriched metabolic pathways were depleted in cave populations and F1 hybrids. **a**, Significant associations/couplings of gut-enriched metabolic pathways with the standardized gut length in surface fish. The gut-enriched pathways were aligned according to the levels of Pearson’s correlation coefficients (*r*: univariate regression, far right panel). All fish individuals treated with three distinct diets per population were used for the sparse modeling regression. Circle sizes represent the -log_10_(P-values of the correlation coefficient). The z-score of each pathway’s abundance (PiCRUSt2 estimated) is shown in the middle column. Colors of the dots are as follows—orange: brine shrimp, green: spirulina, and brown: bat guano. These gut-enriched pathways were also grouped as “Pyruvate generation”, “Cell surface & cell wall”, “Translation capacity”, and “Other microbial function”—custom-made categories based on the MetaCyc ontology (far left panel; Supplementary Table 2). Blue colored pathways were targeted by quercetin pharmacological treatment used in Fig. 3c-e. Z-scored pathway abundances of individual samples from different diets were plotted in different colors (middle panel). **b**, The number of positively and negatively associated/coupled pathways in different populations (surface fish [SF], F1_SP_, Pachón [Pa], F1_SM_, and Molino [Mo]). Surface fish enriched pathways were dominated by the positive pathways, while cave populations and F_1_ hybrids lost most couplings. **c**, The gut-enriched pathways were aligned according to the levels of Pearson’s correlation coefficients (*r*: univariate regression, far right panel). All fish individuals treated with three distinct diets per population were used for the sparse modeling regression. Circle sizes represent the -log_10_(P-values of the correlation coefficient). The z-score of each pathway’s abundance (PiCRUSt2 estimated) is shown in the middle column. Colors of the dots are as follows—orange: brine shrimp, green: spirulina, and brown: bat guano. These gut-enriched pathways were also grouped as “Pyruvate generation” and “Other microbial function”—custom-made categories based on the MetaCyc ontology (far left panel; Supplementary Table 2). “Cell surface & cell wall” and “Translation capacity” were not detected among these populations.

**Extended Data Fig. 9.**
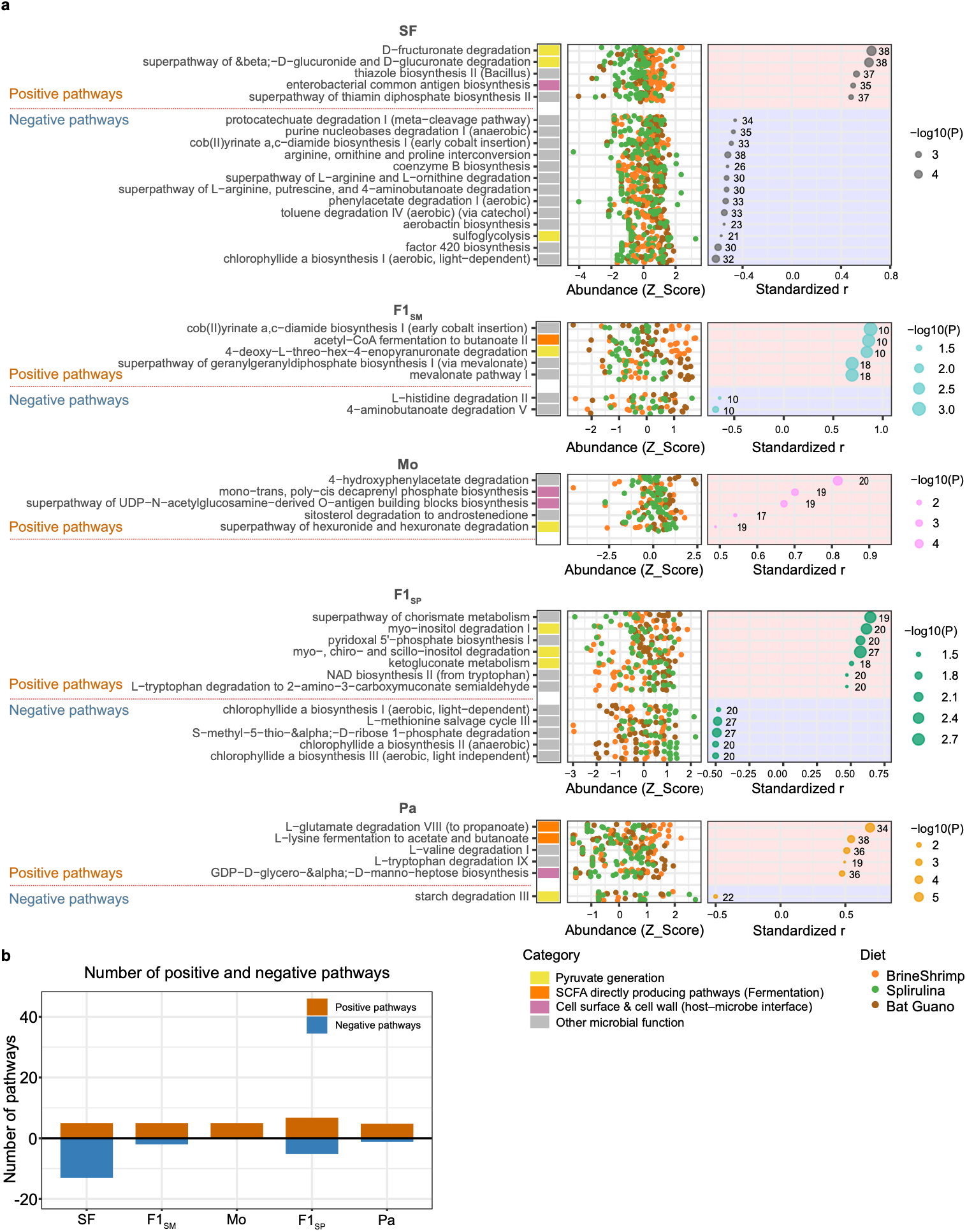
Significant associations/coupling between the midgut width and their gut-enriched pathways in the ancestor-like lineage, surface fish, Molino cavefish and F1 hybrids (F1SP and F1SM). **a,** The gut-enriched pathways were aligned according to the levels of Pearson’s correlation coefficients (*r*: univariate regression, far right panel). All fish individuals treated with three distinct diets per population were used for the sparse modeling regression. Circle sizes represent the -log_10_(P-values of the correlation coefficient). The z-score of each pathway’s abundance (PiCRUSt2 estimated) is shown in the middle column. Colors of the dots are as follows—orange: brine shrimp, green: spirulina, and brown: bat guano. These gut-enriched pathways were also grouped as “Pyruvate generation”, “SCFA direct producing pathway”, “Cell surface & cell wall” and “Other microbial function”—custom-made categories based on the MetaCyc ontology (far left panel; Supplementary Table 3). **b**, The number of positively and negatively associated/coupled pathways in different populations (surface fish [SF], F1_SP_, Pachón [Pa], F1_SM_, and Molino [Mo]). Surface fish enriched pathways were dominated by the positive pathways, while cave populations and F_1_ hybrids lost most couplings.

**Extended Data Fig. 10.**
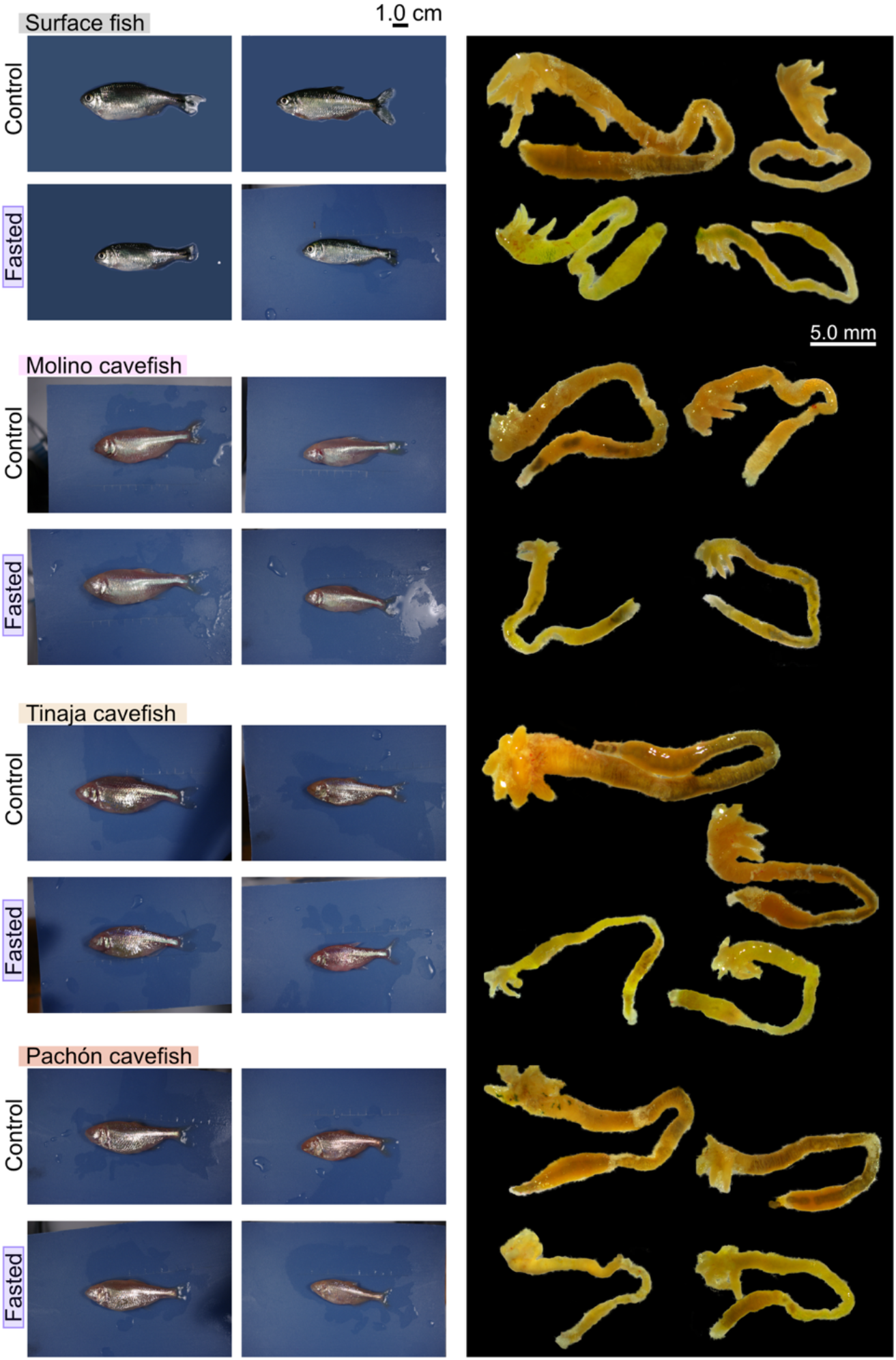
Fish body and gut images after 1 month of fasting. Two representative individuals of fish in each category (populations and fed/fasting) are shown on the left panel, and two representative gut images per category are shown in the right panels. Body shape showed subtle changes during fasting, but gut morphologies indicated that fasting reduced gut size, particularly in Tinaja and Pachón cave populations.

**Extended Data Fig. 11.**
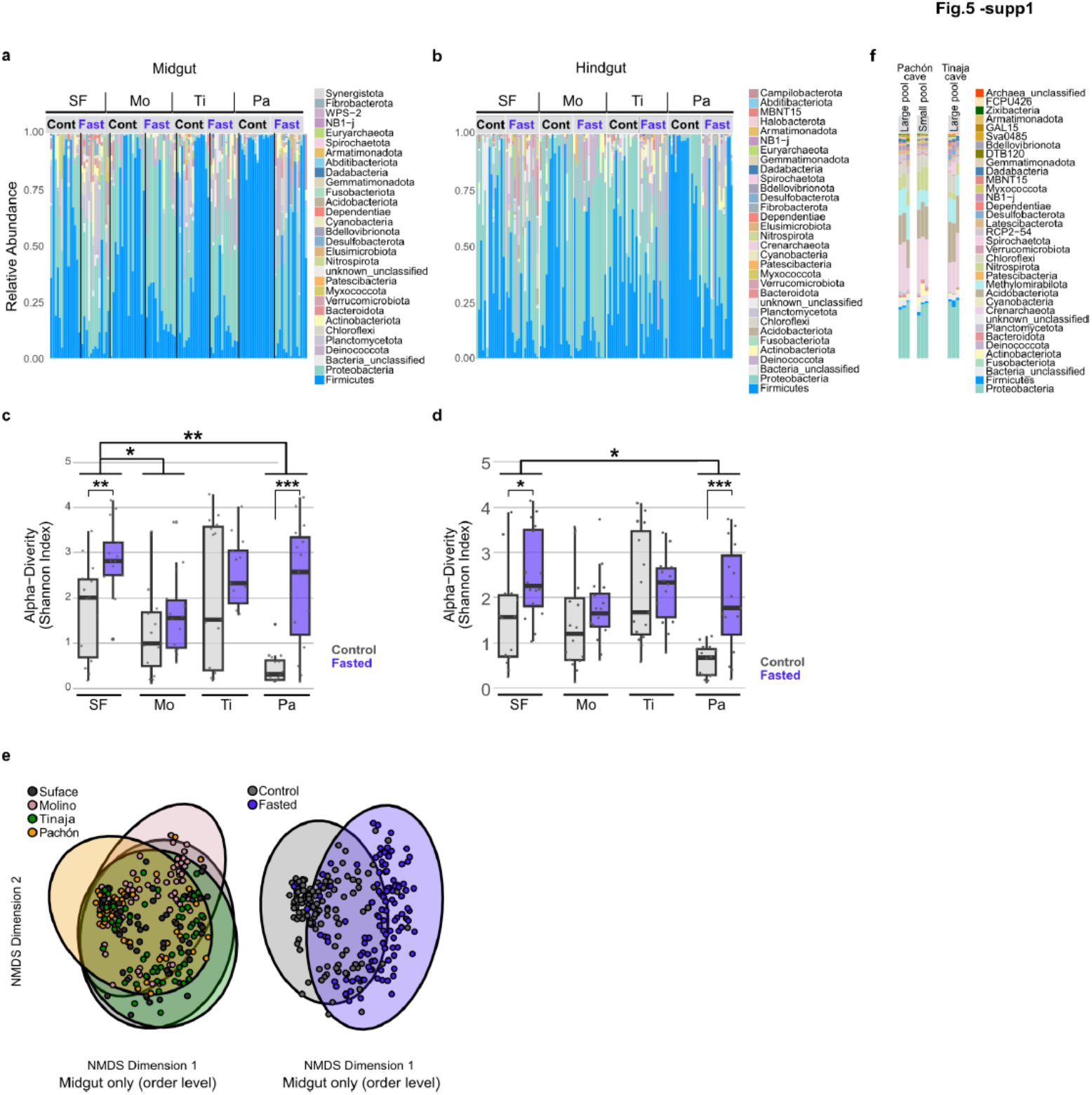
Fasting condition changed gut microbiota across the population. **a, b**, Relative abundance of gut microbiota from midgut (**a**) and hindgut (**b**) from surface fish (SF), Molino (Mo), Tinaja (Ti), and Pachón cavefish (Pa). After one month of fasting, firmicute taxa were reduced in relative abundance in all tested populations. **c, d**, Alpha diversity (Shannon-Weaver Index) of gut microbiota from midgut (**c**) and hindgut (**d**). **e**, The 2-D plot of non-metric multidimensional scaling (NMDS) colored by population (left) and experimental condition (right: the fed control and fasted). The clusters of populations and experimental conditions are largely overlapped. *: P < 0.05, **: P < 0.01, ***: P < 0.001.

**Extended Data Fig. 12.**
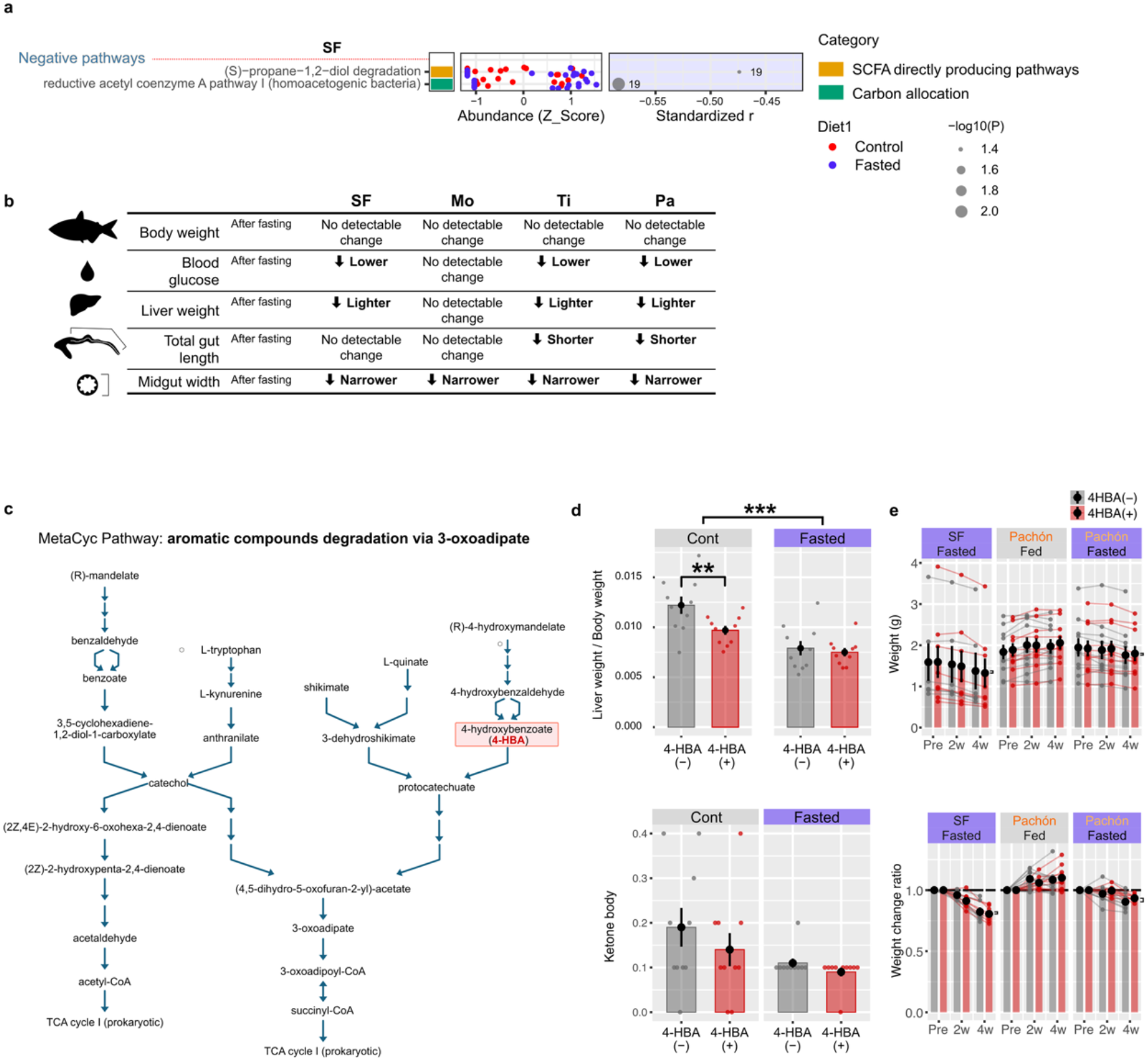
Significant associations/couplings of gut-enriched metabolic pathways with the liver weight (liver weight/body weight) in the ancestor-like surface fish. **a**, The gut-enriched pathways were aligned according to the levels of Pearson’s correlation coefficients (*r*: univariate regression, far right panel). All fish individuals treated with the fed and fasted conditions per population were used for the sparse modeling regression. Circle sizes represent the -log_10_(P-values of the correlation coefficient). The z-score of each pathway’s abundance (PiCRUSt2 estimated) is shown in the middle column. Colors of the dots are as follows—blue: fed control, red: fasted. These liver weight-coupled pathways were also grouped as “SCFA direct producing pathway” and “Carbon allocation”—custom-made categories based on the MetaCyc ontology (far left panel; Supplementary Table 4). **b,** Summary the morphological and physiological changes during fasting. **c,** Aromatic compounds degradation via 3-oxoadipate pathways showed a negative association with liver weight in Pachón cavefish, which were more abundant in the fasted condition. 4-hydroxybenzoate (4-HBA) is one of the substances in this pathway. **d,** (Top row) In the fed control condition, 4-HBA supplementation, which presumably encourages the aromatic compounds degradation pathway, reduced the relative liver weight. In contrast, 4-HBA supplementation induced no detectable effect on liver weight under fasted conditions. This fed control result supports the negative association between the aromatic compounds degradation pathway and liver weight (*c.f.,* Fig. 4a), but, contrary to our prediction, this fasting result indicates 4-HBA supplementation no further liver weight loss, and its end products (*e.g*., succinyl-CoA) might support body weight (*c.f*., **e**). (Bottom row) The serum ketone level did not show any detectable changes in either the fed and fasted condition. **e,** (Top row) Shifts of whole-body weight during 4 weeks of the fed vs. fasted and 4-HBA(-) vs. 4- HBA(+) supplementation conditions. The treatment started from 1-year-old full adults (top-left). No significant difference between 4-HBA(-) vs. 4-HBA(+) was detected. (Bottom row) Weight change rates of the liver weight starting from 3-month-old juvenile fish. 4-HBA might slightly protect against body weight loss in the fasted Pachón cavefish, but it was not detectable. Brackets indicate the differences between 4-HBA(-) vs. 4-HBA(+).

## References

1. Moran, D., Softley, R. & Warrant, E. J. The energetic cost of vision and the evolution of eyeless Mexican cavefish. Science Advances 1, e1500363 (2015).

2. Xiong, S. et al. Enhanced lipogenesis through Pparγ helps cavefish adapt to food scarcity. Current Biology 1–9 (2022) doi:10.1016/j.cub.2022.03.038.

3. Xiong, S., Krishnan, J., Peuß, R. & Rohner, N. Early adipogenesis contributes to excess fat accumulation in cave populations of Astyanax mexicanus. Developmental Biology 441, 297– 304 (2018).

4. Amézquita, E. J., Nasrin, F., Storey, K. M. & Yoshizawa, M. Genomics data analysis via spectral shape and topology. PLoS ONE 18, e0284820 (2023).

5. Bates, J. M. et al. Distinct signals from the microbiota promote different aspects of zebrafish gut differentiation. Developmental Biology 297, 374–386 (2006).

6. Subramanian, S. et al. Cultivating Healthy Growth and Nutrition through the Gut Microbiota. Cell 161, 36–48 (2015).

7. Sommer, F. & Bäckhed, F. The gut microbiota-masters of host development and physiology. Nature Reviews Microbiology 11, 227–238 (2013).

8. Abdellatif, A. M. & Sarvetnick, N. E. Current understanding of the role of gut dysbiosis in type 1 diabetes. Journal of Diabetes 11, 632–644 (2019).

9. Sharma, S. & Tripathi, P. Gut microbiome and type 2 diabetes: where we are and where to go? The Journal of Nutritional Biochemistry 63, 101–108 (2019).

10. Cryan, J. F. & Dinan, T. G. Mind-altering microorganisms: the impact of the gut microbiota on brain and behaviour. Nature reviews Neuroscience 13, 701–12 (2012).

11. Turnbaugh, P. J. & Gordon, J. I. The core gut microbiome, energy balance and obesity. The Journal of Physiology 587, 4153–4158 (2009).

12. Gevers, D. et al. The Treatment-Naive Microbiome in New-Onset Crohn’s Disease. Cell Host & Microbe 15, 382–392 (2014).

13. Morgan, X. C. et al. Dysfunction of the intestinal microbiome in inflammatory bowel disease and treatment. Genome Biol 13, R79 (2012).

14. Rawls, J. F., Mahowald, M. A., Ley, R. E. & Gordon, J. I. Reciprocal Gut Microbiota Transplants from Zebrafish and Mice to Germ-free Recipients Reveal Host Habitat Selection. Cell 127, 423–433 (2006).

15. Walter, J., Britton, R. A. & Roos, S. Host-microbial symbiosis in the vertebrate gastrointestinal tract and the Lactobacillus reuteri paradigm. Proc. Natl. Acad. Sci. U.S.A. 108, 4645–4652 (2011).

16. Dethlefsen, L., McFall-Ngai, M. & Relman, D. A. An ecological and evolutionary perspective on human–microbe mutualism and disease. Nature 449, 811–818 (2007).

17. Moran, R. L. et al. Selection-driven trait loss in independently evolved cavefish populations. Nat Commun 14, 2557 (2023).

18. Herman, A. et al. The role of gene flow in rapid and repeated evolution of cave-related traits in Mexican tetra, Astyanax mexicanus. Molecular Ecology vol. 27 4397–4416 (2018).

19. Fumey, J. et al. Evidence for late Pleistocene origin of Astyanax mexicanus cavefish. BMC Evolutionary Biology 18, 43 (2018).

20. Coghill, L. M., Darrin Hulsey, C., Chaves-Campos, J., García de Leon, F. J. & Johnson, S. G. Next generation phylogeography of cave and surface *Astyanax mexicanus*. Molecular Phylogenetics and Evolution 79, 368–374 (2014).

21. Espinasa, L. et al. Contrasting feeding habits of post-larval and adult *Astyanax* cavefish. Subterranean Biology 21, 1–17 (2017).

22. Culver, D. C. & Pipan, T. The Biology of Caves and Other Subterranean Habitats. The biology of habitats series (Oxford University Press, Oxford, 2009).

23. Fernandes, V. F. L., Macaspac, C., Lu, L. & Yoshizawa, M. Evolution of the developmental plasticity and a coupling between left mechanosensory neuromasts and an adaptive foraging behavior. Developmental Biology 441, 262–271 (2018).

24. Fernandes, V. F. L., Glaser, Y., Iwashita, M. & Yoshizawa, M. Evolution of left–right asymmetry in the sensory system and foraging behavior during adaptation to food-sparse cave environments. BMC Biol 20, 295 (2022).

25. Duboué, E. R., Keene, A. C. & Borowsky, R. L. Evolutionary convergence on sleep loss in cavefish populations. Current Biology 21, 671–676 (2011).

26. Yoshizawa, M. et al. Distinct genetic architecture underlies the emergence of sleep loss and prey-seeking behavior in the Mexican cavefish. BMC biology 13, 15 (2015).

27. Iwashita, M. & Yoshizawa, M. Social-like responses are inducible in asocial Mexican cavefish despite the exhibition of strong repetitive behavior. eLife 10, e72463 (2021).

28. Kowalko, J. E. et al. Genetic analysis of the loss of schooling behavior in cavefish reveals both sight-dependent and independent mechanisms. Curr. Biol. 23, 1874–1883 (2013).

29. Kowalko, J. E. et al. Convergence in feeding posture occurs through different genetic loci in independently evolved cave populations of Astyanax mexicanus. Proceedings of the National Academy of Sciences of the United States of America 110, 16933–16938 (2013).

30. Aspiras, A. C., Rohner, N., Martineau, B., Borowsky, R. L. & Tabin, C. J. Melanocortin 4 receptor mutations contribute to the adaptation of cavefish to nutrient-poor conditions. Proceedings of the National Academy of Sciences of the United States of America 112, 9668– 9673 (2015).

31. Olsen, L., et al. Metabolic reprogramming underlies cavefish muscular endurance despite loss of muscle mass and contractility. Proc. Natl. Acad. Sci. U.S.A. 120, e2204427120 (2023).

32. Krishnan, J. et al. Genome-wide analysis of cis-regulatory changes underlying metabolic adaptation of cavefish. Nat Genet 54, 684–693 (2022).

33. Medley, J. K. et al. The metabolome of Mexican cavefish shows a convergent signature highlighting sugar, antioxidant, and Ageing-Related metabolites. eLife 11, 1–25 (2022).

34. Riddle, M. R. et al. Insulin resistance in cavefish as an adaptation to a nutrient-limited environment. Nature 555, 647–651 (2018).

35. Ornelas-García, P., Pajares, S., Sosa-Jiménez, V. M., Rétaux, S. & Miranda-Gamboa, R. A. Microbiome differences between river-dwelling and cave-adapted populations of the fish astyanax mexicanus (De Filippi, 1853). PeerJ 2018, 1–21 (2018).

36. Riddle, M. R. et al. Host evolution shapes gut microbiome composition in *Astyanax mexicanus*. Ecology and Evolution 14, e11192 (2024).

37. Kozol, R. A. et al. Novel Husbandry Practices Result in Rapid Rates of Growth and Sexual Maturation Without Impacting Adult Behavior in the Blind Mexican Cavefish. Zebrafish 20, 86–94 (2023).

38. Hinaux, H. et al. A Developmental Staging Table for *Astyanax mexicanus* Surface Fish and Pachón Cavefish. Zebrafish 8, 155–165 (2011).

39. Protas, M. et al. Multi-trait evolution in a cave fish, *Astyanax mexicanus*. Evol. Dev. 10, 196– 209 (2008).

40. Armstrong, G. et al. Applications and Comparison of Dimensionality Reduction Methods for Microbiome Data. Front. Bioinform. 2, (2022).

41. Barreñada, L., Dhiman, P., Timmerman, D., Boulesteix, A.-L. & Van Calster, B. Understanding overfitting in random forest for probability estimation: a visualization and simulation study. Diagn Progn Res 8, (2024).

42. Han, S., Williamson, B. D. & Fong, Y. Improving random forest predictions in small datasets from two-phase sampling designs. BMC Med Inform Decis Mak 21, (2021).

43. Francisco, M. E., Carvajal, T. M. & Watanabe, K. Hybrid Machine Learning Approach to Zero-Inflated Data Improves Accuracy of Dengue Prediction. PLoS Negl Trop Dis 18, e0012599 (2024).

44. Paffenhöfer, G.-A. Caloric content of larvae of the brine shrimpArtemia salina. Helgoländer Wissenschaftliche Meeresuntersuchungen 16, 130–135 (1967).

45. Raczyk, M., Polanowska, K., Kruszewski, B., Grygier, A. & Michałowska, D. Effect of Spirulina (Arthrospira platensis) Supplementation on Physical and Chemical Properties of Semolina (Triticum durum) Based Fresh Pasta. Molecules 27, 355 (2022).

46. Mitchell, R. W., Russell, W. H. & Elliott, W. R. Mexican Eyeless Characin Fishes, Genus Astyanax: Environment, Distribution, and Evolution. Special publications the museum Texas Tech University (Texas Tech Press, Texas, 1977).

47. Buddington, R. K. & Diamond, J. M. Aristotle revisited: the function of pyloric caeca in fish. Proc. Natl. Acad. Sci. U.S.A. 83, 8012–8014 (1986).

48. Ballesteros, N. A. et al. The pyloric caeca area is a major site for IgM(+) and IgT(+) B cell recruitment in response to oral vaccination in rainbow trout. PloS one 8, e66118 (2013).

49. McFall-Ngai, M., et al. Animals in a bacterial world, a new imperative for the life sciences. Proc. Natl. Acad. Sci. U.S.A. 110, 3229–3236 (2013).

50. Ley, R. E., Lozupone, C. A., Hamady, M., Knight, R. & Gordon, J. I. Worlds within worlds: evolution of the vertebrate gut microbiota. Nat Rev Microbiol 6, 776–788 (2008).

51. Liao, T., Wei, Y., Luo, M., Zhao, G.-P. & Zhou, H. tmap: an integrative framework based on topological data analysis for population-scale microbiome stratification and association studies. Genome Biol 20, 293 (2019).

52. Chang, W. K., VanInsberghe, D. & Kelly, L. Topological analysis reveals state transitions in human gut and marine bacterial communities. npj Biofilms Microbiomes 6, 41 (2020).

53. Rakoff-Nahoum, S., Foster, K. R. & Comstock, L. E. The evolution of cooperation within the gut microbiota. Nature 533, 255–259 (2016).

54. Turnbaugh, P. J. et al. An obesity-associated gut microbiome with increased capacity for energy harvest. Nature 444, 1027–1031 (2006).

55. Ramond, P., Galand, P. E. & Logares, R. Microbial functional diversity and redundancy: moving forward. FEMS Microbiology Reviews 49, fuae031 (2025).

56. Allison, S. D. & Martiny, J. B. H. Resistance, resilience, and redundancy in microbial communities. Proc. Natl. Acad. Sci. U.S.A. 105, 11512–11519 (2008).

57. Vieira-Silva, S. et al. Species–function relationships shape ecological properties of the human gut microbiome. Nat Microbiol 1, 16088 (2016).

58. Moya, A. & Ferrer, M. Functional Redundancy-Induced Stability of Gut Microbiota Subjected to Disturbance. Trends in Microbiology 24, 402–413 (2016).

59. Douglas, G. M. et al. PICRUSt2 for prediction of metagenome functions. Nat Biotechnol 38, 685–688 (2020).

60. Caspi, R. et al. The MetaCyc database of metabolic pathways and enzymes - a 2019 update. Nucleic Acids Research 48, D445–D453 (2020).

61. Koh, A., De Vadder, F., Kovatcheva-Datchary, P. & Bäckhed, F. From Dietary Fiber to Host Physiology: Short-Chain Fatty Acids as Key Bacterial Metabolites. Cell 165, 1332–1345 (2016).

62. Huang, M. et al. Inhibition of α-amylase activity by quercetin via multi-spectroscopic and molecular docking approaches. Food Bioscience 61, 104951 (2024).

63. Vessal, M., Hemmati, M. & Vasei, M. Antidiabetic effects of quercetin in streptozocin- induced diabetic rats. Comparative Biochemistry and Physiology Part C: Toxicology & Pharmacology 135, 357–364 (2003).

64. Goswami, D., Prajapati, J., Dabhi, M., Sharkey, L. K. R. & Pidot, S. J. MurG as a potential target of quercetin in Staphylococcus aureus supported by evidence from subtractive proteomics and molecular dynamics. Sci Rep 15, 7309 (2025).

65. Riyad, P., Purohit, A., Sen, K., Panwar, A. & Ram, H. HMG – CoA reductase inhibition mediated hypocholesterolemic potential of myricetin and quercetin: *in-silico* and *in-vivo* studies. CyTA - Journal of Food 21, 115–125 (2023).

66. Wilhelm, R. C., DeRito, C. M., Shapleigh, J. P., Madsen, E. L. & Buckley, D. H. Phenolic acid-degrading *Paraburkholderia* prime decomposition in forest soil. ISME Communications 1, 4 (2021).

67. Sugumaran, M. Cuticular sclerotization in insects – A critical review. in Advances in Insect Physiology vol. 62 111–214 (Elsevier, 2022).

68. Van Beynen, P. E., Schwarcz, H. P., Ford, D. C. & Timmins, G. T. Organic substances in cave drip waters: studies from Marengo Cave, Indiana. Can. J. Earth Sci. 39, 279–284 (2002).

69. Alqudah, S. & Claesen, J. Mechanisms of gut bacterial metabolism of dietary polyphenols into bioactive compounds. Gut Microbes 16, 2426614 (2024).

70. Jeffery, W. R. Adaptive evolution of eye degeneration in the Mexican blind cavefish. J. Hered. 96, 185–196 (2005).

71. Protas, M. & Jeffery, W. R. Evolution and development in cave animals: from fish to crustaceans. Wiley Interdiscip. Rev. Dev. Biol. 1, 823–845 (2012).

72. Bairagi, A., Ghosh, K. S., Sen, S. K. & Ray, A. K. Enzyme producing bacterial flora isolated from fish digestive tracts. Aquaculture International 10, 109–121 (2002).

73. Ray, A. K., Ghosh, K. & Ringø, E. Enzyme-producing bacteria isolated from fish gut: a review. Aquaculture Nutrition 18, 465–492 (2012).

74. Ringø, E., Strøm, E. & Tabachek, J.-A. Intestinal microflora of salmonids: a review. Aquaculture Res 26, 773–789 (1995).

75. Ecology and genomics of actinobacteria and their specialised metabolism. (Universiteit Leiden, The Netherlands, 2022).

76. Van Bergeijk, D. A., Terlouw, B. R., Medema, M. H. & Van Wezel, G. P. Ecology and genomics of Actinobacteria: new concepts for natural product discovery. Nat Rev Microbiol 18, 546–558 (2020).

77. Li, Z. et al. Effects of three short-chain fatty acids on growth, intestinal microbiota composition, and ammonia tolerance of juvenile yellow catfish Pelteobagrus fulvidraco. Aquaculture Reports 36, 102066 (2024).

78. Wang, A. et al. Intestinal *Cetobacterium* and acetate modify glucose homeostasis via parasympathetic activation in zebrafish. Gut Microbes 13, 1–15 (2021).

79. Lin, C. et al. Imidazole propionate ameliorates lipid metabolism in adipocytes to attenuate high-fat diet-induced obesity via PPAR signaling pathway. Lipids Health Dis 24, 356 (2025).

80. Koh, A. et al. Microbially Produced Imidazole Propionate Impairs Insulin Signaling through mTORC1. Cell 175, 947–961.e17 (2018).

81. Chen, X., Chen, W., Tang, Y., Zhang, M. & Li, M. Effects of Dietary Sodium Butyrate on Growth Performance, Digestive Ability, Blood Biochemistry, and Ammonia Tolerance of Largemouth Bass (Micropterus salmoides). Fishes 10, 259 (2025).

82. Yan, J., et al. Gut microbiota induce IGF-1 and promote bone formation and growth. Proc. Natl. Acad. Sci. U.S.A.113, (2016).

83. Canosa, L. F. & Bertucci, J. I. Nutrient regulation of somatic growth in teleost fish. The interaction between somatic growth, feeding and metabolism. Molecular and Cellular Endocrinology 518, (2020).

84. Huppop, K. Oxygen-Consumption of Astyanax-Fasciatus (Characidae, Pisces) - a Comparison of Epigean and Hypogean Populations. Environmental Biology of Fishes 17, 299–308 (1986).

85. Bay, S. K. et al. Microbial aerotrophy enables continuous primary production in diverse cave ecosystems. Nat Commun 16, 10295 (2025).

86. Lechleitner, F. A., Dittmar, T., Baldini, J. U. L., Prufer, K. M. & Eglinton, T. I. Molecular signatures of dissolved organic matter in a tropical karst system. Organic Geochemistry 113, 141–149 (2017).

87. Klotzbücher, T., Kaiser, K., Filley, T. R. & Kalbitz, K. Processes controlling the production of aromatic water-soluble organic matter during litter decomposition. Soil Biology and Biochemistry 67, 133–139 (2013).

88. Elipot, Y., Legendre, L., Père, S., Sohm, F. & Rétaux, S. *Astyanax* transgenesis and husbandry: how cavefish enters the laboratory. Zebrafish 11, 291–9 (2014).

89. Keene, A. C., Yoshizawa, M. & McGaugh, S. E. Biology and Evolution of the Mexican Cavefish. Biology and Evolution of the Mexican Cavefish (Elsevier Inc, Amsterdam, 2016). doi:10.1016/C2014-0-01426-8.

90. Schneider, C. A., Rasband, W. S. & Eliceiri, K. W. NIH Image to ImageJ: 25 years of image analysis. Nature Methods 9, 671–675 (2012).

91. Xie, G. et al. A Metabolite Array Technology for Precision Medicine. Anal. Chem. 93, 5709– 5717 (2021).

92. Schindelin, J. et al. Fiji: an open-source platform for biological-image analysis. Nat Methods 9, 676–682 (2012).

93. Parada, A. E., Needham, D. M. & Fuhrman, J. A. Every base matters: assessing small subunit RRNA primers for marine microbiomes with mock communities, time series and global field samples. Environmental Microbiology 18, 1403–1414 (2016).

94. Callahan, B. J. et al. DADA2: High-resolution sample inference from Illumina amplicon data. Nat Methods 13, 581–583 (2016).

95. Schloss, P. D. et al. Introducing mothur: Open-Source, Platform-Independent, Community-Supported Software for Describing and Comparing Microbial Communities. Appl Environ Microbiol 75, 7537–7541 (2009).

96. Quast, C. et al. The SILVA ribosomal RNA gene database project: improved data processing and web-based tools. Nucleic Acids Research 41, D590–D596 (2012).

97. Rognes, T., Flouri, T., Nichols, B., Quince, C. & Mahé, F. VSEARCH: a versatile open source tool for metagenomics. PeerJ 4, e2584 (2016).

98. Frøslev, T. G. et al. Algorithm for post-clustering curation of DNA amplicon data yields reliable biodiversity estimates. Nat Commun 8, 1188 (2017).

99. Geniesse, C., Sporns, O., Petri, G. & Saggar, M. Generating dynamical neuroimaging spatiotemporal representations (DyNeuSR) using topological data analysis. Network Neuroscience 3, 763–778 (2019).

100. Chen, L. et al. GMPR: A robust normalization method for zero-inflated count data with application to microbiome sequencing data. PeerJ 6, e4600 (2018).

101. Walker, B. H. Biodiversity and Ecological Redundancy. Conservation Biology 6, 18–23 (1992).

102. Lawton, J. H. & Brown, V. K. Redundancy in Ecosystems. in Biodiversity and Ecosystem Function (eds Schulze, E.-D. & Mooney, H. A.) 255–270 (Springer Berlin Heidelberg, Berlin, Heidelberg, 1994). doi:10.1007/978-3-642-58001-7_12.

103. Louca, S. & Doebeli, M. Efficient comparative phylogenetics on large trees. Bioinformatics 34, 1053–1055 (2018).

104. Tian, L. et al. Deciphering functional redundancy in the human microbiome. Nat Commun 11, 6217 (2020).

105. Faul, F., Erdfelder, E., Lang, A. G. & Buchner, A. G*Power 3: A flexible statistical power analysis program for the social, behavioral, and biomedical sciences. in Behavior Research Methods vol. 39 175–191 (Springer, 2007).

106. Erdfelder, E., Faul, F., Buchner, A. & Lang, A. G. Statistical power analyses using G*Power 3.1: Tests for correlation and regression analyses. Behavior Research Methods 41, 1149–1160 (2009).

107. Holm, S. A simple sequentially rejective multiple test procedure. Scandinavian journal of statistics (1979).

108. Bates, D., Mächler, M., Bolker, B. M. & Walker, S. C. Fitting linear mixed-effects models using lme4. Journal of Statistical Software 10.18637/jss.v067.i01 (2015) doi:10.18637/jss.v067.i01.

109. Fox, J. & Weisberg, S. An R Companion to Applied Regression.

110. Kuznetsova, A., Brockhoff, P. B. & Christensen, R. H. B. lmerTest Package: Tests in Linear Mixed Effects Models. Journal of Statistical Software https://doi.org/10.18637/jss.v082.i13 (2017) doi:10.18637/jss.v082.i13.

111. Brooks, M. E., et al. glmmTMB Balances Speed and Flexibility Among Packages for Zero-inflated Generalized Linear Mixed Modeling. The R Journal 9, 378 (2017).

112. Cleveland, S. et al. The C-MĀIKI Gateway: A Modern Science Platform for Analyzing Microbiome Data. in Practice and Experience in Advanced Research Computing 1–7 (ACM, Boston MA USA, 2022). doi:10.1145/3491418.3530291.

