## Supplementary Info (Method and Discussion) for "Evolution of novel host-microbiota coupling underlies adaptation to nutrient-scarce environments"

Supplementary Information

**Supplementary Methods**

*Antibiotics treatment for* A. mexicanus *– defining the optimal concentration to attenuate gut microbiota*.

We used (**A**) 10, 20 or 30 µg/mL each of vancomycin (Vancomycin-HCL, VWR International Inc., Radnor, PA, USA), metronidazole (VWR International), cefoxitin (Cefoxitin sodium salt, VWR International), and gentamycin (Gentamycin sulfate, VWR International) for mixing them in the fish diet (see below).

These antibiotics were selected depending on the knowledge of germ-free mice generated by antibiotic treatment ^1^. In the report of Enry et al. ^1^, they used 1mg/mL of the above antibiotics in the drinking water for mice. However, larval fish can be affected and express a behavioral change in much lower concentrations (e.g., 10-30 µM of cefoxitin bath treatment ^2^. 10 µg/mL = 23.4 µM of cefoxitin). Therefore, we adjusted the antibiotics concentration as low as 10 µg/mL instead of 1 mg/mL.

We also used (**B**) commercial antibiotics, which worked to eliminate pathogens in *A. mexicanus*, BiFran+ (Hikari Sales USA, Hayward, CA, USA). BiFran+ consists of Nitrofurazone, Furazolidone and mucus alternative. We used the recommended concentration (6.5% w/v in the diet) for the oral treatment.

To administer the antibiotics, the antibiotic cocktail of 100 µg each of vancomycin, metronidazole, cefoxitin, and gentamycin (each final 10 µg/mL) or 650 mg of BiFran+ powder was mixed with 2 g of ground fish diet (Tetra Tropical Color Granules, Tetra U.S., Blacksburg, VA, USA: final 20 % w/v) and were solidified with 10 mL of 1 % agar (agar, MilliporeSigma, Burlington, MA, USA) poured into a 6-cm Petri dish (VWR international). The control diet had the same components without antibiotics.

These agar foods were cut into 5 mm^3^ cubes and fed each piece per four fish twice daily for 3 or 7 days. Fish were housed in non-sterile ziplock containers with non-sterile fresh conditioned water (pH 6.8-7.1 and conductivity ~700 µS).

To check the depletion of gut microbiota, after the 3 or 7 days of treatment of antibiotics, the experimental and control sets of surface fish were anesthetized with ice-cold water for 5 min. After the gill ventilation ceased, Pyloric caeca and midgut were dissected using sterile forceps and scissors, and transferred to a sterile 6 cm Petri dish and suspended in sterile phosphate-buffered saline (PBS; Santa Cruz Biotechnology, Inc. Dallas, Texas, USA). Then, the pyloric caeca and midgut were ground well in 500 µL of ice-cold PBS using the Bel-Art™ Micro-Tube Homogenizer: Pestles and 1.5 mL Tubes kit (Fisher Scientific, Waltham, MA, USA). The supernatant of homogenates was plated on the brain-heart infusion (BHI) plates (brain-heart infusion broth, MilliporeSigma) and was cultured under an aerobic condition at 25˚C (Extended Data Fig. 5m).

Antibiotics cocktail of vancomycin, metronidazole, cefoxitin and gentamycin were not effective on 10-30 µg/mL in the diet. It may need higher concentration although such concentration may cause behavior alteration without changing gut microbiota (c.f., cefoxitin in 96 hours post fertilization (hpf) of zebrafish ^2^). BiFran+ at the recommended concentration (6.5% w/v) for feeding was successful in depleting microorganisms at 3 days after treatment (Extended Data Fig. 5m). The number of colonies was still small at 7 days of the BiFran+ treated surface fish, although the control-diet-fed surface fish showed a similar number of colonies (Extended Data Fig. 5m).

According to these results, we used BiFran+ at 6.5 % w/v for the following feeding studies.

For the antibiotics experiment, we orally supplemented BiFuran+™ (Hikari Sales USA, Inc.), which contains Nitrofurazone and Furazolidone, and showed an effect to reduce culturable gut microorganisms, compared with other antibiotics. BiFuran+ (6.5 % w/v) was mixed with a ground fish diet (Tetra Tropical Color Granules, Tetra U.S., Blacksburg, VA, USA: final 20 % w/v) and with 1 % agar (agar, MilliporeSigma, Burlington, MA, USA) poured into a 6-cm Petri dish (VWR international, Radnor, Pennsylvania, USA). The control diet consisted of the same composition without BiFuran+. The 11-month-old age-matched surface fish and Pachón cavefish were grouped into 4 fish per still-water container (15.6 × 15.6 × 5.7 cm: Ziplock® containers, S. C. Johnson & Son, Inc.) with ~500 mL conditioned water (pH 7.0-7.2 and conductivity: 700-800 µS). The conditioned water was replaced in full twice a week. We then fed one to two 3 mm^3^ cubes of the above agar diet per four-fish container every morning (ZT 0:00–3:30) and afternoon (ZT 8:00–12:00). The fish were fed ad libitum in each feeding, and the remaining amount was removed 1 h after feeding using a pipette. We fed the antibiotics and control diets for 1 month (4 weeks) before the following assay/analyses.

*Topological Data Analysis (TDA) Mapper and Sparse Modeling for resolving hiding couplings between morphological traits and gut microorganisms*

To visualize the structure of the ASV composition data, we applied a topological data analysis (TDA) mapper^3,4^ (Extended Fig. 3a-c). The mapper partitions the high-dimensional data along a one-dimensional filter function — here the first or second axis of PCA— and clusters samples within each overlapping slice using K-means (Extended Data Fig. 3c). Each cluster becomes a vertex, drawn as a pie chart of its sample composition and scaled by the number of individuals it contains, and vertices with similar ASV composition are joined by links. These links are the informative feature of the representation: whereas ordination methods such as PCoA or NMDS compress all samples into a 2-D or 3-D low-dimensional projection and discard the variance carried by the unplotted axes, the mapper retains every sample and renders the relationships among clusters explicitly as a connectivity graph, so that neighboring and non-neighboring regions of the data can be read directly. Links encode a binary similar/non-similar relation, and link length therefore carries no metric interpretation. Applied to our data, the mapper recovered the intermingled distribution of populations and diet treatments already evident in the NMDS plots (Extended Data Fig. 2g-l vs. Extended Data Fig. 3a,b), but additionally resolved which compositional clusters are contiguous with which, revealing the connectivity structure that the ordinations leave implicit (Extended Data Fig. 3a,b).

Given the complexity of the ASV data structure, we analyzed each population and the F1s separately. Geometric mean of pairwise ratios (GMPR)-standardized ASV counts^5^ were used to construct topological graphs (Extended Data Fig. 3d) with a fish quantitative trait score, such as standard length, as the 1-D filter function (Extended Data Fig. 3c). Filtering on the trait forces individuals to cluster along it, here from −2 to +2 standard deviations of standard length (Extended Data Fig. 3d), which reveals when individuals of similar body length that carry distinct ASV compositions. Such heterogeneity obscures the ASVs coupled to the trait. We therefore extracted subsets of individuals spanning the range from smaller to larger trait values (two examples in turquoise and gray, Extended Data Fig. 3d) — a selection that requires the explicit view of data structure the topological map provides — and applied sparse modeling to each subset to identify morphology-coupled gut ASVs (see below). The same procedure was applied to PICRUSt2-estimated metabolic pathway abundances in place of ASV counts (Extended Data Fig. 3d), which we follow up in the next section.

To search gut ASVs coupled with the host trait score, we applied sparse modeling, an established regression framework for datasets in which the number of measured variables greatly exceeds the number of samples — here ~1,000 ASVs across several dozen fish. Under these conditions, conventional variable selection, such as stepwise backward regression, cannot be applied because a model with more predictors than samples has no unique solution, and no variables can be removed in the first place. Sparse modeling resolves this by penalizing the regression coefficients directly, using a balance of Lasso (L1) and Ridge (L2) regularization: the L1 term shrinks low-contribution ASVs out of the model, while the L2 term stabilizes the estimates for ASVs that co-occur and are therefore mutually correlated (Extended Data Fig. 3e). We surveyed L1 ratios $\frac{\lambda1}{\lambda1+\lambda2}$ from 0.1 (L2 is 0.9) to 1.0 (L2 is 0.0), repeating the regression ten times with random seeding at each ratio, and conservatively retained only ASVs selected in at least eight of the ten repeats to avoid overfitting (Extended Data Fig. 3e). The retained ASVs were then used to estimate standard length, and the estimates were compared with the measured values; among the L1 ratios tested, we selected as our best model the one achieving r² ≧ 0.6 with the fewest ASVs (Extended Data Fig. 3e). Selected ASVs were further validated by univariate regression, and ASVs showing high collinearity with a selected ASV were restored to the final set (Extended Data Fig. 3e), so that a taxon was not discarded merely because a co-occurring taxon carried the same signal.
Importantly, and in contrast to feature-wise univariate approaches that regress each ASV against the phenotype independently (*e.g*., MaAsLin 3^6^), the sparse modeling evaluates all candidate ASVs simultaneously within a single model. The resulting set is therefore not a collection of independently significant taxa but a combination whose abundances jointly and additively account for variation in standard length — implementing the pooled activity of multiple microbial community members^7,8^, which is expected to act on the host traits rather than any single taxon acting alone.

Supplementary Discussion

*The gut length-associated gut microorganisms and their enriched metabolic pathways diminished among cave populations.*

The total gut length (from the base of the stomach to the end of the hindgut; midgut + hindgut; Fig. 1c) was standardized with body standard length, and its associated gut pathways were investigated (Extended Data Fig. 8). Surface fish exhibited 68 gut-length coupled pathways in total with many the positive associations (*r* > 0; Extended Data Fig. 8a), which was the most among other tested groups (Extended Data Fig. 8a-c). The MetaCyc categories indicated many host-microbe interface-associated pathways (*i.e*., cell surface & cell wall) were in the positive association with the gut length in surface fish (Extended Data Fig. 8a), implying these pathways’ products—peptidoglycans, lipopolysaccharides, etc.—would stimulate gut physiology and/or immune development; therefore, they may lead to gut length growth. Such gut-length-coupled pathways were very limited in the F_1_ group and cave populations (Extended Data Fig. 8b,c), implying that genetics may be involved in this association and that retaining these couplings might be costly in the cave environment. Quercetin inhibits the blue colored pathways in Extended Data Fig. 8a, but there was no detectable effect of quercetin on gut lengths of surface fish or other groups under quercetin treatment (Fig. 2e).

Lastly, the midgut width was associated with a relatively smaller number of gut microbe-enriched metabolic pathways across all tested groups, while surface fish exhibited more coupled metabolic pathways than other groups (Extended Data Fig. 9). This was more subtle but showed a similar tendency to that we observed for gut length.

*Applying topological data analysis and sparse modeling to a challenging 16S rRNA gene-based microbiome data (Earth Microbiome Project*^9,10^*)*

The 16S ribosomal RNA gene amplicon-based approaches efficiently detected microbial profiles from small tissue samples and showed relatively high repeatability by technical improvements (*e.g*., standardized DNA extraction and PCR protocols, with improved bioinformatics methods)^11,12^. However, the typical data standardization in 16S amplicon-based studies, such as subsampling and/or converting counts to relative abundance per individual sample, poses challenges that prevent us from applying typical regression methods due to discarding data. Also, the relative abundance score, which includes the closure constraint that all components sum to 1, renders the components non-independent and imposes a negative correlation structure among them. Therefore, pointing out which microbes are associated with the host traits of interest is challenging. Recent updates of a linear regression-based approach (MaAsLin3^6^), centered-log ratio-based ALDEx2^13^ , and Machine learning-based methods^14–16^ have been significantly used for estimating the contributing microbes. However, MaAsLin3 requires a relatively large number of samples for sensitivity (but precision [~0.99] can be achieved with a relatively small number of N = 50 for MaAsLin3^6^), ALDEx2 maintains near-perfect precision but low sensitivity (≤0.27) regardless of sample size^6^, and Random Forest requires a significantly large amount of training samples^17–20^. In addition, if the training data for the random forest does not cover the given sample condition (ex., humans vs. *Astyanax*), it would result in invalid estimates^21,22^.

Our analysis method combined topological data analysis (TDA) with sparse modeling. TDA has been used in microbiome studies^3,4^, but it has not been applied to quantify associations with the host trait of interest. We proposed slicing the data space using the trait of interest (*e.g*., standard length, gut length, liver weight; called a filter function) as the filter function (Extended Data Fig. 3d), instead of the conventional approaches in TDA, which use the 1^st^ or 2^nd^ principal component/ coordinate^3,23^. Such data slices help align clusters across lower- to higher-trait scores. Using similarity links between clusters as a guide, we subset the individual fish samples along paths from lower to higher trait scores.

An advantage of TDA that uses a trait of interest as the filter function is that it can visualize paths from lower to higher scores of the trait by connecting clusters (based on the similarity of ASV profiles) – distinct cluster combinations from lower to higher trait scores represent distinct ASV-profiles (*e.g.,* two representing such cluster sets in cyan and brown shades in Extended Data Fig. 3d). Subsetting removes ASVs profiles from other subset of clusters and facilitates the detection of hidden linear relationship between the trait score and ASV counts, which would be failed to be detected by using a whole dataset. Variation in ASV profiles depends on the initial colonization of microorganisms in the gut and/or the host's immune status. Such variation may mask the genuine linear relationship between the host and gut microbiota observed in certain individuals within the same group. Sparse modeling (Lasso and Ridge regression) is used to detect linear quantitative associations between the host trait and ASV counts or pathway-enriched scores. Sparse modeling is analogous to stepwise backward regression with a distinct algorithm and builds a regression model that includes cooperative additive effects among variables. A multiple regression model seems more appropriate, given common cooperative influence on the host among microbial taxa, compared with a single regression approach (*e.g*., MaAsLin3^6^).

This pipeline consistently detected the same microorganism taxa for body standard length, gut length, and gut width under the antibiotic treatment (bold-faced taxa in Extended Data Fig. 4a-f). Also, several detected Pachón-enriched pathways (pyruvate-generating pathways—an SCFA precursor) were confirmed to promote the growth of Pachón cavefish through a pharmacological inhibition of pyruvate-generating pathways with quercetin (Fig. 2c, Extended Fig. 7a), and SCFA supplementation that promoted Pachón cavefish growth (Extended Data Fig. 7d). Fewer associated gut microbe taxa and pathways for Molino (Fig. 2b, Extended Data Fig. 6) implied that Molino cavefish did not evolve to have support from microorganisms. Indeed, many Molino cavefish deceased during feeding on the lean bat guano (Fig. 1a, b; N = 2, starting with N = 12). Overall, these results support the topological mapping + sparse modeling pipeline, which revealed hidden relationships between gut microbiota and the host.

We employed PiCRUSt2 (v2.5.2) pipeline^24^ to estimate the enrichment of metabolic pathways among microbiota. PiCRUSt2 aligns the ASV sequences to the reference tree in the Integrated Microbial Genomes (IMG) database^25^ and estimate the enriched metabolic pathways with the aid of EPA-NG, gappa, castor, and MinPath packages^26–29^. The IMG database that PiCRUSt2 v2.5 uses is slightly outdated, yet the coverage of microorganisms detected by 16S rRNA gene sequencing was fair (dropped 19 ASVs out of the 1,999 ASVs input into PiCRUSt2, which show Nearest Sequenced Taxon Index [NSTI] ≥ 2 from all archived microorganisms). All these 17 ASVs are unclassified bacteria; therefore, we couldn’t use them for pathway prediction. The most abundant taxon among the 17 ASVs was ASV_29, accounting for ~67% of the removed reads from 19 ASVs and 2.42 - 2.76% of all gut taxa in surface fish and Pachón cavefish, respectively; thus, these unclassified 19 bacteria would have minor bias in our pathway prediction if they had. However, we admit that PiCRUSt2 only provides an estimate of gut-enriched pathway profiles, and the understanding of mechanistic interactions requires further analyses, including gut microbiome transcriptome and/or metabolomics

**References**

1.	Erny, D. *et al.* Host microbiota constantly control maturation and function of microglia in the CNS. *Nat. Neurosci.* **18**, 965–977 (2015).

2.	Rihel, J. *et al.* Zebrafish behavioral profiling links drugs to biological targets and rest/wake regulation. *Science* **327**, 348–51 (2010).

3.	Liao, T., Wei, Y., Luo, M., Zhao, G.-P. & Zhou, H. tmap: an integrative framework based on topological data analysis for population-scale microbiome stratification and association studies. *Genome Biol.* **20**, 293 (2019).

4.	Chang, W. K., VanInsberghe, D. & Kelly, L. Topological analysis reveals state transitions in human gut and marine bacterial communities. *Npj Biofilms Microbiomes* **6**, 41 (2020).

5.	Chen, L. *et al.* GMPR: A robust normalization method for zero-inflated count data with application to microbiome sequencing data. *PeerJ* **6**, e4600 (2018).

6.	Nickols, W. A. *et al.* MaAsLin 3: refining and extending generalized multivariable linear models for meta-omic association discovery. *Nat. Methods* **23**, 554–564 (2026).

7.	Rakoff-Nahoum, S., Foster, K. R. & Comstock, L. E. The evolution of cooperation within the gut microbiota. *Nature* **533**, 255–259 (2016).

8.	Turnbaugh, P. J. *et al.* An obesity-associated gut microbiome with increased capacity for energy harvest. *Nature* **444**, 1027–1031 (2006).

9.	Thompson, L. R. *et al.* A communal catalogue reveals Earth’s multiscale microbial diversity. *Nature* **551**, 457–463 (2017).

10.	Gilbert, J. A., Jansson, J. K. & Knight, R. The Earth Microbiome project: Successes and aspirations. *BMC Biol.* **12**, (2014).

11.	McLaren, M. R., Willis, A. D. & Callahan, B. J. Consistent and correctable bias in metagenomic sequencing experiments. *eLife* **8**, e46923 (2019).

12.	Shaffer, J. P. *et al.* Standardized multi-omics of Earth’s microbiomes reveals microbial and metabolite diversity. *Nat. Microbiol.* **7**, 2128–2150 (2022).

13.	Fernandes, A. D. *et al.* Unifying the analysis of high-throughput sequencing datasets: characterizing RNA-seq, 16S rRNA gene sequencing and selective growth experiments by compositional data analysis. *Microbiome* **2**, 15 (2014).

14.	Pasolli, E., Truong, D. T., Malik, F., Waldron, L. & Segata, N. Machine Learning Meta-analysis of Large Metagenomic Datasets: Tools and Biological Insights. *PLOS Comput. Biol.* **12**, e1004977 (2016).

15.	Yatsunenko, T. *et al.* Human gut microbiome viewed across age and geography. *Nature* **486**, 222–227 (2012).

16.	Knights, D., Costello, E. K. & Knight, R. Supervised classification of human microbiota. *FEMS Microbiol. Rev.* **35**, 343–359 (2011).

17.	Pasolli, E., Truong, D. T., Malik, F., Waldron, L. & Segata, N. Machine Learning Meta-analysis of Large Metagenomic Datasets: Tools and Biological Insights. *PLOS Comput. Biol.* **12**, e1004977 (2016).

18.	Yatsunenko, T. *et al.* Human gut microbiome viewed across age and geography. *Nature* **486**, 222–227 (2012).

19.	Subramanian, S. *et al.* Persistent gut microbiota immaturity in malnourished Bangladeshi children. *Nature* **510**, 417–421 (2014).

20.	Knights, D., Costello, E. K. & Knight, R. Supervised classification of human microbiota. *FEMS Microbiol. Rev.* **35**, 343–359 (2011).

21.	Topçuoğlu, B. D., Lesniak, N. A., Ruffin, M. T., Wiens, J. & Schloss, P. D. A Framework for Effective Application of Machine Learning to Microbiome-Based Classification Problems. *mBio* **11**, e00434-20 (2020).

22.	Nigam, D. ForeCal: Random Forest-based Calibration for DNNs. in *Proceedings of the 8th International Conference on Data Science and Management of Data (12th ACM IKDD CODS and 30th COMAD)* 44–51 (ACM, Jodhpur India, 2024). doi:10.1145/3703323.3703330.

23.	Madukpe, V. N., Ugoala, B. C. & Zulkepli, N. F. S. A Comprehensive Review of the Mapper Algorithm, a Topological Data Analysis Technique, and Its Applications Across Various Fields (2007-2025). Preprint at https://doi.org/10.48550/ARXIV.2504.09042 (2025).

24.	Douglas, G. M. *et al.* PICRUSt2 for prediction of metagenome functions. *Nat. Biotechnol.* **38**, 685–688 (2020).

25.	Markowitz, V. M. *et al.* IMG: the integrated microbial genomes database and comparative analysis system. *Nucleic Acids Res.* **40**, D115–D122 (2012).

26.	Barbera, P. *et al.* Data from: EPA-ng: massively parallel evolutionary placement of genetic sequences. 16601287 bytes https://doi.org/10.5061/DRYAD.KB505NC (2018).

27.	Czech, L., Barbera, P. & Stamatakis, A. Genesis and Gappa: processing, analyzing and visualizing phylogenetic (placement) data. *Bioinformatics* **36**, 3263–3265 (2020).

28.	Louca, S. & Doebeli, M. Efficient comparative phylogenetics on large trees. *Bioinformatics* **34**, 1053–1055 (2018).

29.	Ye, Y. & Doak, T. G. A Parsimony Approach to Biological Pathway Reconstruction/Inference for Genomes and Metagenomes. *PLoS Comput. Biol.* **5**, e1000465 (2009).
