## Supplementary Tables (S1-S5) for "Evolution of novel host-microbiota coupling underlies adaptation to nutrient-scarce environments"

### Supplementary Table 1

| MetaCyc_pathway | category | Lv2 | Lv3 | Lv4 | Lv5 |
|---|---|---|---|---|---|
| methanogenesis from acetate | Other microbial metabolism | Degradation/Utilization/Assim | Acetate Degradation (2) | methanogenesis from acetate | |
| superpathway of polyamine biosynthesis III | Other microbial metabolism | Biosynthesis (2132) | Amide, Amidine, Amine, and Polyamine Biosynthesis (37) | superpathway of polyamine biosynthesis III | |
| creatinine degradation II | Other microbial metabolism | Degradation/Utilization/Assim | Amide, Amidine, Amine, and Polyamine Degradation (68) | Creatinine D | creatinine degradation II |
| creatinine degradation I | Other microbial metabolism | Degradation/Utilization/Assim | Amide, Amidine, Amine, and Polyamine Degradation (69) | Creatinine D | creatinine degradation I |
| glycine betaine degradation I | Other microbial metabolism | Degradation/Utilization/Assim | Amide, Amidine, Amine, and Polyamine Degradation (69) | Glycine Beta | glycine betaine degradation I |
| superpathway of L-alanine biosynthesis | Other microbial metabolism | Biosynthesis (2132) | Amino Acid Biosynthesis (124) | Proteinoger | L-alanine Biosynthesis (4) |
| superpathway of L-aspartate and L-asparagine biosynthesis | Other microbial metabolism | Biosynthesis (2132) | Amino Acid Biosynthesis (124) | Proteinoger | superpathway of L-aspartate and L-asparagine biosynthesis |
| beta-alanine biosynthesis II | Other microbial metabolism | Biosynthesis (2132) | Amino Acid Biosynthesis (124) | Other Amino | beta Alanine Biosynthesis (5) |
| L-methionine salvage cycle III | Other microbial metabolism | Biosynthesis (2132) | Amino Acid Biosynthesis (124) | Proteinoger | L-methionine Biosynthesis (13) |
| S-methyl-5-thio-alpha-D-ribose 1-phosphate degradation I | Other microbial metabolism | Biosynthesis (2132) | Amino Acid Biosynthesis (124) | Proteinoger | L-methionine Biosynthesis (13) |
| L-lysine fermentation to acetate and butanoate | Other microbial metabolism | Degradation/Utilization/Assim | Amino Acid Degradation (136) | Proteinoger | L-lysine Degradation (13) |
| L-arginine degradation II (AST pathway) | Other microbial metabolism | Degradation/Utilization/Assim | Amino Acid Degradation (136) | Proteinoger | L-arginine Degradation (16) |
| chlorosalicylate degradation | Other microbial metabolism | Degradation/Utilization/Assim | Aromatic Compound Degradation (234) | Chloroarom | chlorosalicylate degradation |
| gallate degradation I | Other microbial metabolism | Degradation/Utilization/Assim | Aromatic Compound Degradation (234) | Gallate Deg | gallate degradation I |
| 3-phenylpropanoate degradation | Other microbial metabolism | Degradation/Utilization/Assim | Aromatic Compound Degradation (234) | Phenolic Co | 3-phenylpropanoate degradation |
| coenzyme B biosynthesis | Other microbial metabolism | Biosynthesis (2132) | Biosynthesis (2132) | Cofactor, C | Carrier Biosynthesis (131) |
| 2-carboxy-1,4-naphthoquinol biosynthesis | Other microbial metabolism | Biosynthesis (2132) | Biosynthesis (2132) | Cofactor, C | Carrier Biosynthesis (131) |
| superpathway of phylloquinol biosynthesis | Other microbial metabolism | Biosynthesis (2132) | Biosynthesis (2132) | Cofactor, C | Carrier Biosynthesis (131) |
| superpathway of heme b biosynthesis from uroporphyrinogen-III | Other microbial metabolism | Biosynthesis (2132) | Biosynthesis (2132) | Cofactor, C | Enzyme Cofactor Biosynthesis (142) |
| superpathway of heme b biosynthesis from uroporphyrinogen-III | Other microbial metabolism | Biosynthesis (2132) | Biosynthesis (2132) | Tetrapyrrole | Porphyrin Compound Biosynthesis (39) |
| reductive acetyl coenzyme A pathway I (homoacetogenic bacteria) | Other microbial metabolism | Degradation/Utilization/Assim | C1 Compound Utilization and Assimilation (39) | CO2 Fixation | Autotrophic CO2 Fixation (12) |
| Calvin-Benson-Bassham cycle | Other microbial metabolism | Degradation/Utilization/Assim | C1 Compound Utilization and Assimilation (39) | CO2 Fixation | Autotrophic CO2 Fixation (12) |
| colanic acid building blocks biosynthesis | Other microbial metabolism | Biosynthesis (2132) | Carbohydrate Biosynthesis (240) | colanic acid building blocks biosynthesis | |
| glycogen biosynthesis I (from ADP-D-Glucose) | Other microbial metabolism | Biosynthesis (2132) | Carbohydrate Biosynthesis (240) | Polysaccha | Glycogen and Starch Biosynthesis (4) |
| starch biosynthesis | Other microbial metabolism | Biosynthesis (2132) | Carbohydrate Biosynthesis (240) | Polysaccha | Glycogen and Starch Biosynthesis (4) |
| UDP-N-acetyl-D-glucosamine biosynthesis I | Other microbial metabolism | Biosynthesis (2132) | Carbohydrate Biosynthesis (240) | Sugar Biosy | Sugar Nucleotide Biosynthesis (94) |
| Calvin-Benson-Bassham cycle | Other microbial metabolism | Biosynthesis (2132) | Carbohydrate Biosynthesis (240) | Sugar Biosy | Calvin-Benson-Bassham cycle |
| gluconeogenesis I | Other microbial metabolism | Biosynthesis (2132) | Carbohydrate Biosynthesis (240) | Sugar Biosy | Gluconeogenesis (3) |
| GDP-mannose biosynthesis | Other microbial metabolism | Biosynthesis (2132) | Carbohydrate Biosynthesis (240) | Sugar Biosy | Sugar Nucleotide Biosynthesis (94) |
| superpathway of GDP-mannose-derived O-antigen building blocks biosynthesis | Other microbial metabolism | Biosynthesis (2132) | Carbohydrate Biosynthesis (240) | Sugar Biosy | Sugar Nucleotide Biosynthesis (94) |
| enterobacterial common antigen biosynthesis | Other microbial metabolism | Biosynthesis (2132) | Carbohydrate Biosynthesis (240) | Polysaccha | enterobacterial common antigen biosynthesis |
| GDP-D-glycero-alpha-D-manno-heptose biosynthesis | Other microbial metabolism | Biosynthesis (2132) | Carbohydrate Biosynthesis (240) | Sugar Biosy | Sugar Nucleotide Biosynthesis (94) |
| GDP-D-glycero-alpha-D-manno-heptose biosynthesis | Other microbial metabolism | Biosynthesis (2132) | Carbohydrate Biosynthesis (240) | Sugar Biosy | Sugar Nucleotide Biosynthesis (94) |
| starch degradation V | Pyruvate generation | Degradation/Utilization/Assim | Carbohydrate Degradation (180) | Polysaccha | Starch Degradation (5) |
| glycogen degradation I | Pyruvate generation | Degradation/Utilization/Assim | Carbohydrate Degradation (180) | Polysaccha | Glycogen Degradation (3) |
| sucrose degradation IV (sucrose phosphorylase) | Pyruvate generation | Degradation/Utilization/Assim | Carbohydrate Degradation (180) | Sugar Degra | Sucrose Degradation (7) |
| glucose and glucose-1-phosphate degradation | Pyruvate generation | Degradation/Utilization/Assim | Carbohydrate Degradation (180) | Sugar Degra | glucose and glucose-1-phosphate degradation |
| sucrose degradation III (sucrose invertase) | Pyruvate generation | Degradation/Utilization/Assim | Carbohydrate Degradation (180) | Sugar Degra | Sucrose Degradation (7) |
| 1,5-anhydrofructose degradation | Pyruvate generation | Degradation/Utilization/Assim | Carbohydrate Degradation (180) | Sugar Degra | 1,5-anhydrofructose degradation |
| chitin derivatives degradation | Pyruvate generation | Degradation/Utilization/Assim | Carbohydrate Degradation (180) | Sugar Degra | Sugar Derivative Degradation (16) |
| sulfoquinovose degradation I | Pyruvate generation | Degradation/Utilization/Assim | Carbohydrate Degradation (180) | Sulfoquino | sulfoquinovose degradation I |
| mycolyl-arabinogalactan-peptidoglycan complex biosynthesis | Other microbial metabolism | Biosynthesis (2132) | Cell Structure Biosynthesis (124) | Cell Wall Bi | mycolyl-arabinogalactan-peptidoglycan complex biosynthesis |
| UDP-N-acetylmuramoyl-pentapeptide biosynthesis II (lysine-containing) | Other microbial metabolism | Biosynthesis (2132) | Cell Structure Biosynthesis (124) | Cell Wall Bi | UDP-N-Acetylmuramoyl-Pentapeptide Biosynthesis (3) |
| peptidoglycan biosynthesis I (meso-diaminopimelate containing) | Other microbial metabolism | Biosynthesis (2132) | Cell Structure Biosynthesis (124) | Cell Wall Bi | Peptidoglycan Biosynthesis (6) |
| UDP-N-acetylmuramoyl-pentapeptide biosynthesis I (meso-diaminopimelate containir | Other microbial metabolism | Biosynthesis (2132) | Cell Structure Biosynthesis (124) | Cell Wall Bi | UDP-N-Acetylmuramoyl-Pentapeptide Biosynthesis (3) |
| peptidoglycan biosynthesis III (mycobacteria) | Other microbial metabolism | Biosynthesis (2132) | Cell Structure Biosynthesis (124) | Cell Wall Bi | Peptidoglycan Biosynthesis (6) |
| peptidoglycan maturation (meso-diaminopimelate containing) | Other microbial metabolism | Biosynthesis (2132) | Cell Structure Biosynthesis (124) | Cell Wall Bi | Peptidoglycan Biosynthesis (6) |
| superpathway of (Kdo)2-lipid A biosynthesis | Other microbial metabolism | Biosynthesis (2132) | Cell Structure Biosynthesis (124) | Lipopolysac | superpathway of (Kdo)2-lipid A biosynthesis |
| chlorosalicylate degradation | Other microbial metabolism | Degradation/Utilization/Assim | Chlorinated Compound Degradation (45) | Chloroarom | chlorosalicylate degradation |
| biotin biosynthesis II | Other microbial metabolism | Biosynthesis (2132) | Cofactor, Carrier, and Vitamin Biosynthesis (346) | Enzyme Co | Biotin Biosynthesis (5) |
| NAD de novo biosynthesis I | Other microbial metabolism | Biosynthesis (2132) | Cofactor, Carrier, and Vitamin Biosynthesis (346) | Carrier Biosy | Electron Carrier Biosynthesis (92) |
| heme b biosynthesis II (oxygen-independent) | Other microbial metabolism | Biosynthesis (2132) | Cofactor, Carrier, and Vitamin Biosynthesis (346) | Enzyme Co | Heme Biosynthesis (13) |
| superpathway of heme b biosynthesis from glutamate | Other microbial metabolism | Biosynthesis (2132) | Cofactor, Carrier, and Vitamin Biosynthesis (346) | Enzyme Co | Heme Biosynthesis (13) |
| heme b biosynthesis I (aerobic) | Other microbial metabolism | Biosynthesis (2132) | Cofactor, Carrier, and Vitamin Biosynthesis (346) | Enzyme Co | Heme Biosynthesis (13) |
| NAD de novo biosynthesis I | Other microbial metabolism | Biosynthesis (2132) | Cofactor, Carrier, and Vitamin Biosynthesis (346) | Enzyme Co | NAD Biosynthesis (15) |
| NAD de novo biosynthesis I | Other microbial metabolism | Biosynthesis (2132) | Cofactor, Carrier, and Vitamin Biosynthesis (346) | Enzyme Co | NAD Metabolism (22) |
| 5,8-dihydroxy-2-naphthoate biosynthesis I | Other microbial metabolism | Biosynthesis (2132) | Cofactor, Carrier, and Vitamin Biosynthesis (346) | Carrier Biosy | Electron Carrier Biosynthesis (92) |
| vitamin E biosynthesis (tocopherols) | Other microbial metabolism | Biosynthesis (2132) | Cofactor, Carrier, and Vitamin Biosynthesis (346) | Carrier Biosy | Electron Carrier Biosynthesis (92) |
| vitamin E biosynthesis (tocopherols) | Other microbial metabolism | Biosynthesis (2132) | Cofactor, Carrier, and Vitamin Biosynthesis (346) | Vitamin Bios | vitamin E biosynthesis (tocopherols) |
| nylon-6 oligomer degradation | Other microbial metabolism | Degradation/Utilization/Assim | Degradation/Utilization/Assimilation - Other (45) | nylon-6 oligomer degradation | |
| (S)-propane-1,2-diol degradation | Other microbial metabolism | Degradation/Utilization/Assim | Degradation/Utilization/Assimilation (1140) | Alcohol Deg | (S)-propane-1,2-diol degradation |
| superpathway of methylglyoxal degradation | Pyruvate generation | Degradation/Utilization/Assim | Degradation/Utilization/Assimilation (1140) | Aldehyde D | superpathway of methylglyoxal degradation |
| catechol degradation I (meta-cleavage pathway) | Other microbial metabolism | Degradation/Utilization/Assim | Degradation/Utilization/Assimilation (1140) | Aromatic Co | Catechol Degradation (7) |
| 3-phenylpropanoate degradation | Other microbial metabolism | Degradation/Utilization/Assim | Degradation/Utilization/Assimilation (1140) | Aromatic Co | Phenolic Compound Degradation (27) |
| beta-D-glucuronide and D-glucuronate degradation | Pyruvate generation | Degradation/Utilization/Assim | Degradation/Utilization/Assimilation (1140) | Carbohydra | Sugar Degradation (95) |
| beta-D-glucuronide and D-glucuronate degradation | Pyruvate generation | Degradation/Utilization/Assim | Degradation/Utilization/Assimilation (1140) | Carboxylic A | Uronic Acid Degradation (11) |
| myo-, chiro- and scyllo-inositol degradation | Pyruvate generation | Degradation/Utilization/Assim | Degradation/Utilization/Assimilation (1140) | Cyclitol Deg | myo-, chiro- and scyllo-inositol degradation |
| myo-inositol degradation I | Pyruvate generation | Degradation/Utilization/Assim | Degradation/Utilization/Assimilation (1140) | Cyclitol Deg | myo-inositol degradation I |
| superpathway of pyrimidine deoxyribonucleosides degradation | Other microbial metabolism | Degradation/Utilization/Assim | Degradation/Utilization/Assimilation (1140) | Nucleoside | Pyrimidine Nucleotide Degradation (13) |
| aerobic respiration I (cytochrome c) | Other microbial metabolism | Generation of Precursor Metal | Electron Transfer Chains (48) | aerobic respiration I (cytochrome c) | |
| superpathway of (Kdo)2-lipid A biosynthesis | Other microbial metabolism | Biosynthesis (2132) | Fatty Acid and Lipid Biosynthesis (262) | superpathway of (Kdo)2-lipid A biosynthesis | |
| L-lysine fermentation to acetate and butanoate | SCFA directly producing pathways | Generation of Precursor Metal | Fermentation (128) | Fermentatio | Fermentation to Acetate (23) |
| L-lysine fermentation to acetate and butanoate | SCFA directly producing pathways | Generation of Precursor Metal | Fermentation (128) | Fermentatio | Fermentation to Butanoate (8) |
| mixed acid fermentation | SCFA directly producing pathways | Generation of Precursor Metal | Fermentation (128) | Fermentatio | Pyruvate Fermentation to Ethanol (7) |
| acetylene degradation (anaerobic) | SCFA directly producing pathways | Generation of Precursor Metal | Fermentation (128) | Fermentatio | Pyruvate Fermentation to Ethanol (7) |
| mixed acid fermentation | SCFA directly producing pathways | Generation of Precursor Metal | Fermentation (128) | Fermentatio | Pyruvate Fermentation to Ethanol (7) |
| acetylene degradation (anaerobic) | SCFA directly producing pathways | Generation of Precursor Metal | Fermentation (128) | Fermentatio | Pyruvate Fermentation to Ethanol (7) |
| mixed acid fermentation | SCFA directly producing pathways | Generation of Precursor Metal | Fermentation (128) | Fermentatio | mixed acid fermentation |
| acetylene degradation (anaerobic) | SCFA directly producing pathways | Generation of Precursor Metal | Fermentation (128) | Fermentatio | Fermentation to Acetate (23) |
| (S)-propane-1,2-diol degradation | Other microbial metabolism | Generation of Precursor Metal | Generation of Precursor Metabolites and Energy (256) | Fermentatio | Fermentation to Short-Chain Fatty Acids (42) |
| glycogen degradation I | Pyruvate generation | Glycan Pathways (175) | glycogen degradation I | | |
| glycolysis I (from glucose 6-phosphate) | Pyruvate generation | Generation of Precursor Metal | Glycolysis (6) | glycolysis I (from glucose 6-phosphate) | |
| glycolysis II (from fructose 6-phosphate) | Pyruvate generation | Generation of Precursor Metal | Glycolysis (6) | glycolysis II (from fructose 6-phosphate) | |
| nitrate reduction VI (assimilatory) | Other microbial metabolism | Degradation/Utilization/Assim | Inorganic Nutrient Metabolism (159) | Nitrogen Co | Nitrate Reduction (11) |
| superpathway of (Kdo)2-lipid A biosynthesis | Other microbial metabolism | Glycan Pathways (175) | Lipopolysaccharide Biosynthesis (86) | superpathway of (Kdo)2-lipid A biosynthesis | |
| mycolyl-arabinogalactan-peptidoglycan complex biosynthesis | Other microbial metabolism | Glycan Pathways (175) | mycolyl-arabinogalactan-peptidoglycan complex biosynthesis | | |
| 5-aminoimidazole ribonucleotide biosynthesis II | Other microbial metabolism | Biosynthesis (2132) | Nucleoside and Nucleotide Biosynthesis (68) | Purine Nucl | 5-Aminoimidazole Ribonucleotide Biosynthesis (3) |
| superpathway of 5-aminoimidazole ribonucleotide biosynthesis | Other microbial metabolism | Biosynthesis (2132) | Nucleoside and Nucleotide Biosynthesis (68) | Purine Nucl | 5-Aminoimidazole Ribonucleotide Biosynthesis (3) |
| superpathway of pyrimidine ribonucleotides de novo biosynthesis | Other microbial metabolism | Biosynthesis (2132) | Nucleoside and Nucleotide Biosynthesis (68) | Pyrimidine I | Pyrimidine Nucleotide De Novo Biosynthesis (11) |
| pyrimidine deoxyribonucleotide phosphorylation | Other microbial metabolism | Biosynthesis (2132) | Nucleoside and Nucleotide Biosynthesis (68) | Pyrimidine I | Pyrimidine Nucleotide Salvage (12) |
| superpathway of pyrimidine nucleobases salvage | Other microbial metabolism | Biosynthesis (2132) | Nucleoside and Nucleotide Biosynthesis (68) | Pyrimidine I | Pyrimidine Nucleotide Salvage (12) |
| S-methyl-5-thio-alpha-D-ribose 1-phosphate degradation I | Other microbial metabolism | Degradation/Utilization/Assim | Nucleoside and Nucleotide Degradation (42) | S-methyl-5- | S-methyl-5-thio-alpha-D-ribose 1-phosphate degradation I |
| 6-hydroxymethyl-dihydropterin diphosphate biosynthesis I | Other microbial metabolism | Biosynthesis (2132) | Other Biosynthesis (65) | 6-Hydroxym | 6-hydroxymethyl-dihydropterin diphosphate biosynthesis I |
| pentose phosphate pathway | Other microbial metabolism | Generation of Precursor Metal | Pentose Phosphate Pathways (6) | pentose phosphate pathway | |
| starch degradation V | Pyruvate generation | Degradation/Utilization/Assim | Polymeric Compound Degradation (50) | Polysaccha | Starch Degradation (5) |
| glycogen degradation I | Pyruvate generation | Degradation/Utilization/Assim | Polymeric Compound Degradation (50) | Polysaccha | Glycogen Degradation (3) |
| superpathway of geranylgeranyl diphosphate biosynthesis II (via MEP) | Other microbial metabolism | Biosynthesis (2132) | Polyprenyl Biosynthesis (18) | All-trans Po | Geranylgeranyl Diphosphate Biosynthesis (3) |
| glycogen biosynthesis I (from ADP-D-Glucose) | Other microbial metabolism | Glycan Pathways (175) | Polysaccharide Biosynthesis (78) | Glycogen ar | glycogen biosynthesis I (from ADP-D-Glucose) |
| starch biosynthesis | Other microbial metabolism | Glycan Pathways (175) | Polysaccharide Biosynthesis (78) | Glycogen ar | starch biosynthesis |
| enterobacterial common antigen biosynthesis | Other microbial metabolism | Glycan Pathways (175) | Polysaccharide Biosynthesis (78) | enterobacterial common antigen biosynthesis | |
| pyrimidine deoxyribonucleotide phosphorylation | Other microbial metabolism | Metabolic Clusters (70) | pyrimidine deoxyribonucleotide phosphorylation | | |
| methanogenesis from acetate | Other microbial metabolism | Generation of Precursor Metal | Respiration (70) | Anaerobic F | Methanogenesis (16) |
| aerobic respiration I (cytochrome c) | Other microbial metabolism | Generation of Precursor Metal | Respiration (70) | Aerobic Res | aerobic respiration I (cytochrome c) |
| preQ0 biosynthesis | Other microbial metabolism | Biosynthesis (2132) | Secondary Metabolite Biosynthesis (765) | preQ0 biosynthesis | |
| methylerythritol phosphate pathway I | Other microbial metabolism | Biosynthesis (2132) | Secondary Metabolite Biosynthesis (765) | Terpenoid E | Hemiterpene Biosynthesis (10) |
| methylerythritol phosphate pathway II | Other microbial metabolism | Biosynthesis (2132) | Secondary Metabolite Biosynthesis (765) | Terpenoid E | Hemiterpene Biosynthesis (10) |
| superpathway of geranylgeranyl diphosphate biosynthesis II (via MEP) | Other microbial metabolism | Biosynthesis (2132) | Secondary Metabolite Biosynthesis (765) | Terpenoid E | Diterpenoid Biosynthesis (44) |
| mevalonate pathway II (haloarchaea) | Other microbial metabolism | Biosynthesis (2132) | Secondary Metabolite Biosynthesis (765) | Terpenoid E | Hemiterpene Biosynthesis (10) |
| ergothioneine biosynthesis I (bacteria) | Other microbial metabolism | Biosynthesis (2132) | Secondary Metabolite Biosynthesis (908) | Ergothionei | ergothioneine biosynthesis I (bacteria) |
| heme b biosynthesis II (oxygen-independent) | Other microbial metabolism | Biosynthesis (2132) | Tetrapyrrole Biosynthesis (39) | Porphyrin C | Heme Biosynthesis (13) |
| superpathway of heme b biosynthesis from glutamate | Other microbial metabolism | Biosynthesis (2132) | Tetrapyrrole Biosynthesis (39) | Porphyrin C | Heme Biosynthesis (13) |
| uroporphyrinogen-III II (from glycine) | Other microbial metabolism | Biosynthesis (2132) | Tetrapyrrole Biosynthesis (39) | Porphyrin C | Uroporphyrinogen-III Biosynthesis (2) |
| heme b biosynthesis I (aerobic) | Other microbial metabolism | Biosynthesis (2132) | Tetrapyrrole Biosynthesis (39) | Porphyrin C | Heme Biosynthesis (13) |
| tetrapyrrole biosynthesis I (from glutamate) | Other microbial metabolism | Biosynthesis (2132) | Tetrapyrrole Biosynthesis (39) | Porphyrin C | Uroporphyrinogen-III Biosynthesis (2) |

### Supplementary Table 2

| MetaCyc_pathway | category | Lv2 | Lv3 | Lv4 | Lv5 |
|---|---|---|---|---|---|
| superpathway of L-threonine biosynthesis | Other microbial metabolism | Biosynthesis (2132) | Amino Acid Biosynthesis (124) | Proteinogenic Amino Acid Biosynthesis (95) | L-threonine Biosynthesis (2) |
| superpathway of L-isoleucine biosynthesis I | Other microbial metabolism | Biosynthesis (2132) | Amino Acid Biosynthesis (124) | Proteinogenic Amino Acid Biosynthesis (95) | L-isoleucine Biosynthesis (6) |
| L-lysine biosynthesis VI | Other microbial metabolism | Biosynthesis (2132) | Amino Acid Biosynthesis (124) | Proteinogenic Amino Acid Biosynthesis (95) | L-lysine Biosynthesis (6) |
| L-lysine biosynthesis III | Other microbial metabolism | Biosynthesis (2132) | Amino Acid Biosynthesis (124) | Proteinogenic Amino Acid Biosynthesis (95) | L-lysine Biosynthesis (6) |
| L-isoleucine biosynthesis III | Other microbial metabolism | Biosynthesis (2132) | Amino Acid Biosynthesis (124) | Proteinogenic Amino Acid Biosynthesis (95) | L-isoleucine Biosynthesis (6) |
| tRNA charging | Translation capacity | Biosynthesis (2132) | Aminoacyl-tRNA Charging (4) | tRNA charging | |
| starch biosynthesis | Other microbial metabolism | Biosynthesis (2132) | Carbohydrate Biosynthesis (240) | Polysaccharide Biosynthesis (78) | Glycogen and Starch Biosynthesis (4) |
| superpathway of UDP-N-acetylglucosamine-derived O-antigen building blocks biosynthesis | Cell surface & cell wall (host–microbe interface) | Biosynthesis (2132) | Carbohydrate Biosynthesis (240) | Sugar Biosynthesis (110) | Sugar Nucleotide Biosynthesis (94) |
| UDP-N-acetyl-D-glucosamine biosynthesis I | Cell surface & cell wall (host–microbe interface) | Biosynthesis (2132) | Carbohydrate Biosynthesis (240) | Sugar Biosynthesis (110) | Sugar Nucleotide Biosynthesis (94) |
| Calvin-Benson-Bassham cycle | Other microbial metabolism | Biosynthesis (2132) | Carbohydrate Biosynthesis (240) | Sugar Biosynthesis (110) | Calvin-Benson-Bassham cycle |
| CMP-3-deoxy-D-manno-octulosonate biosynthesis | Cell surface & cell wall (host–microbe interface) | Biosynthesis (2132) | Carbohydrate Biosynthesis (240) | Sugar Biosynthesis (110) | Sugar Nucleotide Biosynthesis (94) |
| O-antigen building blocks biosynthesis (E. coli) | Cell surface & cell wall (host–microbe interface) | Biosynthesis (2132) | Carbohydrate Biosynthesis (240) | Sugar Biosynthesis (110) | Sugar Nucleotide Biosynthesis (94) |
| dTDP-L-rhamnose biosynthesis | Cell surface & cell wall (host–microbe interface) | Biosynthesis (2132) | Carbohydrate Biosynthesis (240) | Sugar Biosynthesis (110) | Sugar Nucleotide Biosynthesis (94) |
| UDP-2,3-diacetamido-2,3-dideoxy-alpha-D-mannuronate biosynthesis | Cell surface & cell wall (host–microbe interface) | Biosynthesis (2132) | Carbohydrate Biosynthesis (240) | Sugar Biosynthesis (110) | Sugar Nucleotide Biosynthesis (94) |
| peptidoglycan biosynthesis I (meso-diaminopimelate containing) | Cell surface & cell wall (host–microbe interface) | Biosynthesis (2132) | Cell Structure Biosynthesis (124) | Cell Wall Biosynthesis (27) | Peptidoglycan Biosynthesis (6) |
| UDP-N-acetylmuramoyl-pentapeptide biosynthesis I (meso-diaminopimelate containing) | Cell surface & cell wall (host–microbe interface) | Biosynthesis (2132) | Cell Structure Biosynthesis (124) | Cell Wall Biosynthesis (27) | UDP-N-Acetylmuramoyl-Pentapeptide Biosynthesis (3) |
| peptidoglycan biosynthesis III (mycobacteria) | Cell surface & cell wall (host–microbe interface) | Biosynthesis (2132) | Cell Structure Biosynthesis (124) | Cell Wall Biosynthesis (27) | Peptidoglycan Biosynthesis (6) |
| UDP-N-acetylmuramoyl-pentapeptide biosynthesis II (lysine-containing) | Cell surface & cell wall (host–microbe interface) | Biosynthesis (2132) | Cell Structure Biosynthesis (124) | Cell Wall Biosynthesis (27) | UDP-N-Acetylmuramoyl-Pentapeptide Biosynthesis (3) |
| peptidoglycan biosynthesis II (staphylococci) | Cell surface & cell wall (host–microbe interface) | Biosynthesis (2132) | Cell Structure Biosynthesis (124) | Cell Wall Biosynthesis (27) | Peptidoglycan Biosynthesis (6) |
| lipid IVA biosynthesis (E. coli) | Cell surface & cell wall (host–microbe interface) | Biosynthesis (2132) | Cell Structure Biosynthesis (124) | Lipopolysaccharide Biosynthesis (86) | Lipid IVA Biosynthesis (7) |
| coenzyme B biosynthesis | Other microbial metabolism | Biosynthesis (2132) | Cofactor, Carrier, and Vitamin Biosynthesis (346) | Carrier Biosynthesis (131) | Electron Carrier Biosynthesis (92) |
| flavin biosynthesis I (bacteria and plants) | Other microbial metabolism | Biosynthesis (2132) | Cofactor, Carrier, and Vitamin Biosynthesis (346) | Carrier Biosynthesis (131) | Electron Carrier Biosynthesis (92) |
| coenzyme A biosynthesis I (bacteria) | Other microbial metabolism | Biosynthesis (2132) | Cofactor, Carrier, and Vitamin Biosynthesis (346) | Carrier Biosynthesis (131) | Coenzyme A Biosynthesis (10) |
| folate transformations III (E. coli) | Other microbial metabolism | Biosynthesis (2132) | Cofactor, Carrier, and Vitamin Biosynthesis (346) | Carrier Biosynthesis (131) | Single Carbon Carrier Biosynthesis (23) |
| superpathway of tetrahydrofolate biosynthesis and salvage | Other microbial metabolism | Biosynthesis (2132) | Cofactor, Carrier, and Vitamin Biosynthesis (346) | Carrier Biosynthesis (131) | Single Carbon Carrier Biosynthesis (23) |
| superpathway of tetrahydrofolate biosynthesis | Other microbial metabolism | Biosynthesis (2132) | Cofactor, Carrier, and Vitamin Biosynthesis (346) | Carrier Biosynthesis (131) | Single Carbon Carrier Biosynthesis (23) |
| NAD de novo biosynthesis I | Other microbial metabolism | Biosynthesis (2132) | Cofactor, Carrier, and Vitamin Biosynthesis (346) | Carrier Biosynthesis (131) | Electron Carrier Biosynthesis (92) |
| flavin biosynthesis I (bacteria and plants) | Other microbial metabolism | Biosynthesis (2132) | Cofactor, Carrier, and Vitamin Biosynthesis (346) | Enzyme Cofactor Biosynthesis (157) | Flavin Biosynthesis (5) |
| NAD de novo biosynthesis I | Other microbial metabolism | Biosynthesis (2132) | Cofactor, Carrier, and Vitamin Biosynthesis (346) | Enzyme Cofactor Biosynthesis (157) | NAD Biosynthesis (15) |
| NAD de novo biosynthesis I | Other microbial metabolism | Biosynthesis (2132) | Cofactor, Carrier, and Vitamin Biosynthesis (346) | Enzyme Cofactor Biosynthesis (157) | NAD Metabolism (22) |
| folate transformations III (E. coli) | Other microbial metabolism | Biosynthesis (2132) | Cofactor, Carrier, and Vitamin Biosynthesis (346) | Vitamin Biosynthesis (58) | Folate Biosynthesis (10) |
| superpathway of tetrahydrofolate biosynthesis and salvage | Other microbial metabolism | Biosynthesis (2132) | Cofactor, Carrier, and Vitamin Biosynthesis (346) | Vitamin Biosynthesis (58) | Folate Biosynthesis (10) |
| superpathway of tetrahydrofolate biosynthesis | Other microbial metabolism | Biosynthesis (2132) | Cofactor, Carrier, and Vitamin Biosynthesis (346) | Vitamin Biosynthesis (58) | Folate Biosynthesis (10) |
| palmitate biosynthesis II (type II fatty acid synthase) | Other microbial metabolism | Biosynthesis (2132) | Fatty Acid and Lipid Biosynthesis (262) | Fatty Acid Biosynthesis (99) | Palmitate Biosynthesis (3) |
| lipid IVA biosynthesis (E. coli) | Cell surface & cell wall (host–microbe interface) | Biosynthesis (2132) | Fatty Acid and Lipid Biosynthesis (262) | Lipid IVA Biosynthesis (7) | lipid IVA biosynthesis (E. coli) |
| adenosine deoxyribonucleotides de novo biosynthesis II | Other microbial metabolism | Biosynthesis (2132) | Nucleoside and Nucleotide Biosynthesis (68) | 2'-Deoxyribonucleotide Biosynthesis (11) | Purine Deoxyribonucleotide De Novo Biosynthesis (5) |
| guanosine deoxyribonucleotides de novo biosynthesis II | Other microbial metabolism | Biosynthesis (2132) | Nucleoside and Nucleotide Biosynthesis (68) | 2'-Deoxyribonucleotide Biosynthesis (11) | Purine Deoxyribonucleotide De Novo Biosynthesis (5) |
| pyrimidine deoxyribonucleotides de novo biosynthesis I | Other microbial metabolism | Biosynthesis (2132) | Nucleoside and Nucleotide Biosynthesis (68) | 2'-Deoxyribonucleotide Biosynthesis (11) | Pyrimidine Deoxyribonucleotide De Novo Biosynthesis (6) |
| superpathway of pyrimidine deoxyribonucleotides de novo biosynthesis (E. coli) | Other microbial metabolism | Biosynthesis (2132) | Nucleoside and Nucleotide Biosynthesis (68) | 2'-Deoxyribonucleotide Biosynthesis (11) | Pyrimidine Deoxyribonucleotide De Novo Biosynthesis (6) |
| 5-aminoimidazole ribonucleotide biosynthesis II | Other microbial metabolism | Biosynthesis (2132) | Nucleoside and Nucleotide Biosynthesis (68) | Purine Nucleotide Biosynthesis (33) | 5-Aminoimidazole Ribonucleotide Biosynthesis (3) |
| superpathway of 5-aminoimidazole ribonucleotide biosynthesis | Other microbial metabolism | Biosynthesis (2132) | Nucleoside and Nucleotide Biosynthesis (68) | Purine Nucleotide Biosynthesis (33) | 5-Aminoimidazole Ribonucleotide Biosynthesis (3) |
| inosine-5'-phosphate biosynthesis I | Other microbial metabolism | Biosynthesis (2132) | Nucleoside and Nucleotide Biosynthesis (68) | Purine Nucleotide Biosynthesis (33) | Purine Nucleotide De Novo Biosynthesis (16) |
| superpathway of adenosine nucleotides de novo biosynthesis I | Other microbial metabolism | Biosynthesis (2132) | Nucleoside and Nucleotide Biosynthesis (68) | Purine Nucleotide Biosynthesis (33) | Purine Nucleotide De Novo Biosynthesis (16) |
| 5-aminoimidazole ribonucleotide biosynthesis I | Other microbial metabolism | Biosynthesis (2132) | Nucleoside and Nucleotide Biosynthesis (68) | Purine Nucleotide Biosynthesis (33) | 5-Aminoimidazole Ribonucleotide Biosynthesis (3) |
| adenosine ribonucleotides de novo biosynthesis | Other microbial metabolism | Biosynthesis (2132) | Nucleoside and Nucleotide Biosynthesis (68) | Purine Nucleotide Biosynthesis (33) | Purine Nucleotide De Novo Biosynthesis (16) |
| guanosine ribonucleotides de novo biosynthesis | Other microbial metabolism | Biosynthesis (2132) | Nucleoside and Nucleotide Biosynthesis (68) | Purine Nucleotide Biosynthesis (33) | Purine Nucleotide De Novo Biosynthesis (16) |
| superpathway of purine nucleotides de novo biosynthesis I | Other microbial metabolism | Biosynthesis (2132) | Nucleoside and Nucleotide Biosynthesis (68) | Purine Nucleotide Biosynthesis (33) | Purine Nucleotide De Novo Biosynthesis (16) |
| superpathway of guanosine nucleotides de novo biosynthesis I | Other microbial metabolism | Biosynthesis (2132) | Nucleoside and Nucleotide Biosynthesis (68) | Purine Nucleotide Biosynthesis (33) | Purine Nucleotide De Novo Biosynthesis (16) |
| superpathway of adenosine nucleotides de novo biosynthesis II | Other microbial metabolism | Biosynthesis (2132) | Nucleoside and Nucleotide Biosynthesis (68) | Purine Nucleotide Biosynthesis (33) | Purine Nucleotide De Novo Biosynthesis (16) |
| superpathway of guanosine nucleotides de novo biosynthesis II | Other microbial metabolism | Biosynthesis (2132) | Nucleoside and Nucleotide Biosynthesis (68) | Purine Nucleotide Biosynthesis (33) | Purine Nucleotide De Novo Biosynthesis (16) |
| adenosine deoxyribonucleotides de novo biosynthesis II | Other microbial metabolism | Biosynthesis (2132) | Nucleoside and Nucleotide Biosynthesis (68) | Purine Nucleotide Biosynthesis (33) | Purine Nucleotide De Novo Biosynthesis (16) |
| guanosine deoxyribonucleotides de novo biosynthesis II | Other microbial metabolism | Biosynthesis (2132) | Nucleoside and Nucleotide Biosynthesis (68) | Purine Nucleotide Biosynthesis (33) | Purine Nucleotide De Novo Biosynthesis (16) |
| superpathway of pyrimidine nucleobases salvage | Other microbial metabolism | Biosynthesis (2132) | Nucleoside and Nucleotide Biosynthesis (68) | Pyrimidine Nucleotide Biosynthesis (23) | Pyrimidine Nucleotide Salvage (12) |
| UMP biosynthesis I | Other microbial metabolism | Biosynthesis (2132) | Nucleoside and Nucleotide Biosynthesis (68) | Pyrimidine Nucleotide Biosynthesis (23) | Pyrimidine Nucleotide De Novo Biosynthesis (11) |
| superpathway of pyrimidine ribonucleotides de novo biosynthesis | Other microbial metabolism | Biosynthesis (2132) | Nucleoside and Nucleotide Biosynthesis (68) | Pyrimidine Nucleotide Biosynthesis (23) | Pyrimidine Nucleotide De Novo Biosynthesis (11) |
| pyrimidine deoxyribonucleotide phosphorylation | Other microbial metabolism | Biosynthesis (2132) | Nucleoside and Nucleotide Biosynthesis (68) | Pyrimidine Nucleotide Biosynthesis (23) | Pyrimidine Nucleotide Salvage (12) |
| pyrimidine deoxyribonucleotides de novo biosynthesis I | Other microbial metabolism | Biosynthesis (2132) | Nucleoside and Nucleotide Biosynthesis (68) | Pyrimidine Nucleotide Biosynthesis (23) | Pyrimidine Nucleotide De Novo Biosynthesis (11) |
| superpathway of pyrimidine deoxyribonucleotides de novo biosynthesis (E. coli) | Other microbial metabolism | Biosynthesis (2132) | Nucleoside and Nucleotide Biosynthesis (68) | Pyrimidine Nucleotide Biosynthesis (23) | Pyrimidine Nucleotide De Novo Biosynthesis (11) |
| 6-hydroxymethyl-dihydropterin diphosphate biosynthesis III (Chlamydia) | Other microbial metabolism | Biosynthesis (2132) | Other Biosynthesis (65) | 6-Hydroxymethyl-Dihydropterin Diphosphate Biosynt | 6-hydroxymethyl-dihydropterin diphosphate biosynthesis III (Chlamydia) |
| 6-hydroxymethyl-dihydropterin diphosphate biosynthesis I | Other microbial metabolism | Biosynthesis (2132) | Other Biosynthesis (65) | 6-Hydroxymethyl-Dihydropterin Diphosphate Biosynt | 6-hydroxymethyl-dihydropterin diphosphate biosynthesis I |
| 3,8-divinyl-chlorophyllide a biosynthesis I (aerobic, light-dependent) | Other microbial metabolism | Biosynthesis (2132) | Secondary Metabolite Biosynthesis (908) | Pigment Biosynthesis (89) | Chlorophyll Biosynthesis (15) |
| preQ0 biosynthesis | Other microbial metabolism | Biosynthesis (2132) | Secondary Metabolite Biosynthesis (908) | preQ0 biosynthesis | |
| methylerythritol phosphate pathway I | Other microbial metabolism | Biosynthesis (2132) | Secondary Metabolite Biosynthesis (908) | Terpenoid Biosynthesis (252) | Hemiterpene Biosynthesis (10) |
| methylerythritol phosphate pathway II | Other microbial metabolism | Biosynthesis (2132) | Secondary Metabolite Biosynthesis (908) | Terpenoid Biosynthesis (252) | Hemiterpene Biosynthesis (10) |
| uroporphyrinogen-III II (from glycine) | Other microbial metabolism | Biosynthesis (2132) | Tetrapyrrole Biosynthesis (39) | Porphyrin Compound Biosynthesis (39) | Uroporphyrinogen-III Biosynthesis (2) |
| 3,8-divinyl-chlorophyllide a biosynthesis I (aerobic, light-dependent) | Other microbial metabolism | Biosynthesis (2132) | Tetrapyrrole Biosynthesis (39) | Porphyrin Compound Biosynthesis (39) | Chlorophyll Biosynthesis (15) |
| tetrapyrrole biosynthesis I (from glutamate) | Other microbial metabolism | Biosynthesis (2132) | Tetrapyrrole Biosynthesis (39) | Porphyrin Compound Biosynthesis (39) | Uroporphyrinogen-III Biosynthesis (2) |
| allantoin degradation IV (anaerobic) | Other microbial metabolism | Degradation/Utilization/Assimilati | Amide, Amidine, Amine, and Polyamine Degradatior | Allantoin Degradation (8) | allantoin degradation IV (anaerobic) |
| L-histidine degradation I | Other microbial metabolism | Degradation/Utilization/Assimilati | Amino Acid Degradation (136) | Proteinogenic Amino Acid Degradation (115) | L-histidine Degradation (6) |
| 2-aminophenol degradation | Other microbial metabolism | Degradation/Utilization/Assimilati | Aromatic Compound Degradation (234) | 2-aminophenol degradation | |
| benzoyl-CoA degradation I (aerobic) | Other microbial metabolism | Degradation/Utilization/Assimilati | Aromatic Compound Degradation (234) | Benzoyl-CoA Degradation (3) | benzoyl-CoA degradation I (aerobic) |
| gallate degradation II | Other microbial metabolism | Degradation/Utilization/Assimilati | Aromatic Compound Degradation (234) | Gallate Degradation (3) | gallate degradation II |
| methylgallate degradation | Other microbial metabolism | Degradation/Utilization/Assimilati | Aromatic Compound Degradation (234) | methylgallate degradation | |
| Calvin-Benson-Bassham cycle | Other microbial metabolism | Degradation/Utilization/Assimilati | C1 Compound Utilization and Assimilation (39) | CO2 Fixation (15) | Autotrophic CO2 Fixation (12) |
| sucrose degradation III (sucrose invertase) | Pyruvate generation | Degradation/Utilization/Assimilati | Carbohydrate Degradation (180) | Sugar Degradation (96) | Sucrose Degradation (7) |
| lactose degradation I | Pyruvate generation | Degradation/Utilization/Assimilati | Carbohydrate Degradation (180) | Sugar Degradation (96) | Lactose Degradation (3) |
| glucose degradation (oxidative) | Pyruvate generation | Degradation/Utilization/Assimilati | Carbohydrate Degradation (180) | Sugar Degradation (96) | glucose degradation (oxidative) |
| phospholipases | Other microbial metabolism | Degradation/Utilization/Assimilati | Fatty Acid and Lipid Degradation (55) | phospholipases | |
| inosine 5'-phosphate degradation | Other microbial metabolism | Degradation/Utilization/Assimilati | Nucleoside and Nucleotide Degradation (42) | Purine Nucleotide Degradation (16) | inosine 5'-phosphate degradation |
| glycolysis III (from glucose) | Pyruvate generation | Generation of Precursor Metaboli | Glycolysis (6) | glycolysis III (from glucose) | |
| pentose phosphate pathway (non-oxidative branch) I | Other microbial metabolism | Generation of Precursor Metaboli | Pentose Phosphate Pathways (6) | pentose phosphate pathway (non-oxidative branch) I | |
| lipid IVA biosynthesis (E. coli) | Cell surface & cell wall (host–microbe interface) | Glycan Pathways (175) | Lipopolysaccharide Biosynthesis (86) | Lipid IVA Biosynthesis (7) | lipid IVA biosynthesis (E. coli) |
| starch biosynthesis | Other microbial metabolism | Glycan Pathways (175) | Polysaccharide Biosynthesis (78) | Glycogen and Starch Biosynthesis (4) | starch biosynthesis |
| O-antigen building blocks biosynthesis (E. coli) | Cell surface & cell wall (host–microbe interface) | Metabolic Clusters (70) | O-antigen building blocks biosynthesis (E. coli) | | |
| phospholipases | Other microbial metabolism | Metabolic Clusters (70) | phospholipases | | |
| pyrimidine deoxyribonucleotide phosphorylation | Other microbial metabolism | Metabolic Clusters (70) | pyrimidine deoxyribonucleotide phosphorylation | | |
| pyrimidine deoxyribonucleotides de novo biosynthesis I | Other microbial metabolism | Metabolic Clusters (70) | pyrimidine deoxyribonucleotides de novo biosynthesis I | | |
| tRNA charging | Translation capacity | Metabolic Clusters (70) | tRNA charging | | |
| beta-alanine biosynthesis II | Other microbial metabolism | Biosynthesis (2132) | Amino Acid Biosynthesis (124) | Other Amino Acid Biosynthesis (26) | beta Alanine Biosynthesis (5) |
| syringate degradation | Other microbial metabolism | Degradation/Utilization/Assimilati | Aromatic Compound Degradation (234) | syringate degradation | |
| nitrate reduction VI (assimilatory) | Other microbial metabolism | Degradation/Utilization/Assimilati | Inorganic Nutrient Metabolism (159) | Nitrogen Compound Metabolism (36) | Nitrate Reduction (11) |
| syringate degradation | Other microbial metabolism | Superpathways (384) | syringate degradation | | |
| sucrose biosynthesis III | Other microbial metabolism | Biosynthesis (2132) | Carbohydrate Biosynthesis (240) | Sugar Biosynthesis (110) | Disaccharide Biosynthesis (11) |
| sucrose biosynthesis I (from photosynthesis) | Other microbial metabolism | Biosynthesis (2132) | Carbohydrate Biosynthesis (240) | Sugar Biosynthesis (110) | Disaccharide Biosynthesis (11) |
| L-arabinose degradation IV | Pyruvate generation | Degradation/Utilization/Assimilati | Carbohydrate Degradation (180) | Sugar Degradation (95) | L-arabinose Degradation (5) |
| L-fucose degradation I | Pyruvate generation | Degradation/Utilization/Assimilati | Carbohydrate Degradation (180) | Sugar Degradation (95) | L-Fucose Degradation (3) |
| sucrose biosynthesis I (from photosynthesis) | Other microbial metabolism | Superpathways (384) | sucrose biosynthesis I (from photosynthesis) | | |
| allantoin degradation to glyoxylate III | Other microbial metabolism | Superpathways (384) | allantoin degradation to glyoxylate III | | |
| ectoine biosynthesis | Other microbial metabolism | Biosynthesis (2132) | Amide, Amidine, Amine, and Polyamine Biosynthesi | ectoine biosynthesis | |
| norspermidine biosynthesis | Other microbial metabolism | Biosynthesis (2132) | Amide, Amidine, Amine, and Polyamine Biosynthesi | norspermidine biosynthesis | |
| allantoin degradation to glyoxylate III | Other microbial metabolism | Degradation/Utilization/Assimilati | Amide, Amidine, Amine, and Polyamine Degradatior | Allantoin Degradation (8) | allantoin degradation to glyoxylate III |
| creatinine degradation I | Other microbial metabolism | Degradation/Utilization/Assimilati | Amide, Amidine, Amine, and Polyamine Degradatior | Creatinine Degradation (3) | creatinine degradation I |
| superpathway of menaquinol-8 biosynthesis II | Other microbial metabolism | Biosynthesis (2132) | Cofactor, Carrier, and Vitamin Biosynthesis (252) | Carrier Biosynthesis (131) | Electron Carrier Biosynthesis (92) |
| nylon-6 oligomer degradation | Other microbial metabolism | Degradation/Utilization/Assimilati | Degradation/Utilization/Assimilation - Other (45) | nylon-6 oligomer degradation | |
| enterobactin biosynthesis | Other microbial metabolism | Superpathways (384) | enterobactin biosynthesis | | |
| ethylmalonyl-CoA pathway | Other microbial metabolism | Generation of Precursor Metaboli | ethylmalonyl-CoA pathway | | |
| enterobactin biosynthesis | Other microbial metabolism | Biosynthesis (2132) | Secondary Metabolite Biosynthesis (765) | Siderophore and Metallophore Biosynthesis (33) | enterobactin biosynthesis |
| superpathway of menaquinol-8 biosynthesis II | Other microbial metabolism | Superpathways (384) | superpathway of menaquinol-8 biosynthesis II | | |

### Supplementary Table 3

| MetaCyc_pathway | category | Lv2 | Lv3 | Lv4 | Lv5 |
|---|---|---|---|---|---|
| L-arginine degradation XIII (reductive Stickland reaction) | Other microbial metabolism | Generation of Precursor Metabolites and Energy (256) | Fermentation (84) | Amino Acid Fermentation (Stickland reactions) (16) | Stickland Fermentation (Reductive Branch) (9) |
| L-arginine degradation XIII (reductive Stickland reaction) | Other microbial metabolism | Degradation/Utilization/Assimilation (1140) | Amino Acid Degradation (135) | Proteinogenic Amino Acid Degradation (115) | L-arginine Degradation (16) |
| sulfoquinovose degradation I | Pyruvate generation | Degradation/Utilization/Assimilation (1140) | Carbohydrate Degradation (178) | Sulfoquinovose Degradation (6) | sulfoquinovose degradation I |
| phenylacetate degradation I (aerobic) | Other microbial metabolism | Degradation/Utilization/Assimilation (1140) | Aromatic Compound Degradation (230) | Phenolic Compound Degradation (27) | Phenylacetate Degradation (2) |
| coenzyme B biosynthesis | Other microbial metabolism | Biosynthesis (2132) | Cofactor, Carrier, and Vitamin Biosynthesis (252) | Carrier Biosynthesis (131) | Electron Carrier Biosynthesis (92) |
| D-fructuronate degradation | Pyruvate generation | Degradation/Utilization/Assimilation (1140) | Carboxylic Acid Degradation (85) | Uronic Acid Degradation (11) | D-fructuronate degradation |
| D-fructuronate degradation | Pyruvate generation | Degradation/Utilization/Assimilation (1140) | Carbohydrate Degradation (178) | Sugar Degradation (95) | Uronic Acid Degradation (11) |
| protocatechuate degradation I (meta-cleavage pathway) | Other microbial metabolism | Degradation/Utilization/Assimilation (1140) | Aromatic Compound Degradation (230) | Protocatechuate Degradation (4) | protocatechuate degradation I (meta-cleavage pathway) |
| aerobactin biosynthesis | Other microbial metabolism | Biosynthesis (2132) | Secondary Metabolite Biosynthesis (765) | Siderophore and Metallophore Biosynthesis (33) | aerobactin biosynthesis |
| toluene degradation IV (aerobic) (via catechol) | Other microbial metabolism | Superpathways (384) | toluene degradation IV (aerobic) (via catechol) | | |
| toluene degradation IV (aerobic) (via catechol) | Other microbial metabolism | Degradation/Utilization/Assimilation (1140) | Aromatic Compound Degradation (230) | Toluene Degradation (14) | toluene degradation IV (aerobic) (via catechol) |
| 3PG-factor 420 biosynthesis | Other microbial metabolism | Biosynthesis (2132) | Cofactor, Carrier, and Vitamin Biosynthesis (252) | Carrier Biosynthesis (131) | Electron Carrier Biosynthesis (92) |
| cob(II)yrinate a,c-diamide biosynthesis I (early cobalt insertion) | Other microbial metabolism | Biosynthesis (2132) | Cofactor, Carrier, and Vitamin Biosynthesis (252) | Enzyme Cofactor Biosynthesis (142) | cob(II)yrinate a,c-diamide biosynthesis (2) |
| superpathway of L-arginine and L-ornithine degradation | Other microbial metabolism | Superpathways (384) | superpathway of L-arginine and L-ornithine degradation | | |
| superpathway of L-arginine and L-ornithine degradation | Other microbial metabolism | Degradation/Utilization/Assimilation (1140) | Amino Acid Degradation (135) | Proteinogenic Amino Acid Degradation (115) | L-arginine Degradation (16) |
| superpathway of L-arginine, putrescine, and 4-aminobutanoate degradation | Other microbial metabolism | Superpathways (384) | superpathway of L-arginine, putrescine, and 4-aminobutanoate degradation | | |
| superpathway of L-arginine, putrescine, and 4-aminobutanoate degradation | Other microbial metabolism | Degradation/Utilization/Assimilation (1140) | Amino Acid Degradation (135) | Proteinogenic Amino Acid Degradation (115) | L-arginine Degradation (16) |
| superpathway of thiamine diphosphate biosynthesis II | Other microbial metabolism | Superpathways (384) | superpathway of thiamine diphosphate biosynthesis II | | |
| superpathway of thiamine diphosphate biosynthesis II | Other microbial metabolism | Biosynthesis (2132) | Cofactor, Carrier, and Vitamin Biosynthesis (252) | Vitamin Biosynthesis (58) | Thiamine Biosynthesis (21) |
| superpathway of thiamine diphosphate biosynthesis II | Other microbial metabolism | Biosynthesis (2132) | Cofactor, Carrier, and Vitamin Biosynthesis (252) | Enzyme Cofactor Biosynthesis (142) | Thiamine Biosynthesis (21) |
| thiazole component of thiamine diphosphate biosynthesis II | Other microbial metabolism | Biosynthesis (2132) | Cofactor, Carrier, and Vitamin Biosynthesis (252) | Vitamin Biosynthesis (58) | Thiamine Biosynthesis (21) |
| thiazole component of thiamine diphosphate biosynthesis II | Other microbial metabolism | Biosynthesis (2132) | Cofactor, Carrier, and Vitamin Biosynthesis (252) | Enzyme Cofactor Biosynthesis (142) | Thiamine Biosynthesis (21) |
| beta-D-glucuronide and D-glucuronate degradation | Pyruvate generation | Degradation/Utilization/Assimilation (1140) | Carboxylic Acid Degradation (85) | Uronic Acid Degradation (11) | D-Glucuronate Degradation (5) |
| beta-D-glucuronide and D-glucuronate degradation | Pyruvate generation | Degradation/Utilization/Assimilation (1140) | Carbohydrate Degradation (178) | Sugar Degradation (95) | Uronic Acid Degradation (11) |
| enterobacterial common antigen biosynthesis | Cell surface & cell wall (host–microbe int | Glycan Pathways (175) | Polysaccharide Biosynthesis (78) | enterobacterial common antigen biosynthesis | |
| enterobacterial common antigen biosynthesis | Cell surface & cell wall (host–microbe int | Biosynthesis (2132) | Carbohydrate Biosynthesis (236) | Polysaccharide Biosynthesis (78) | enterobacterial common antigen biosynthesis |
| purine nucleobases degradation I (anaerobic) | Other microbial metabolism | Generation of Precursor Metabolites and Energy (256) | Fermentation (84) | purine nucleobases degradation I (anaerobic) | |
| purine nucleobases degradation I (anaerobic) | Other microbial metabolism | Degradation/Utilization/Assimilation (1140) | Nucleoside and Nucleotide Degradation (41) | Purine Nucleotide Degradation (16) | purine nucleobases degradation I (anaerobic) |
| 3,8-divinyl-chlorophyllide a biosynthesis I (aerobic, light-dependent) | Other microbial metabolism | Biosynthesis (2132) | Tetrapyrrole Biosynthesis (39) | Porphyrin Compound Biosynthesis (39) | Chlorophyll Biosynthesis (15) |
| 3,8-divinyl-chlorophyllide a biosynthesis I (aerobic, light-dependent) | Other microbial metabolism | Biosynthesis (2132) | Secondary Metabolite Biosynthesis (765) | Pigment Biosynthesis (89) | Chlorophyll Biosynthesis (15) |
| L-glutamate degradation VIII (to propanoate) | SCFA directly producing pathways | Superpathways (384) | L-glutamate degradation VIII (to propanoate) | | |
| L-glutamate degradation VIII (to propanoate) | SCFA directly producing pathways | Generation of Precursor Metabolites and Energy (256) | Fermentation (84) | Fermentation to Short-Chain Fatty Acids (42) | Fermentation to Propanoate (7) |
| L-glutamate degradation VIII (to propanoate) | SCFA directly producing pathways | Degradation/Utilization/Assimilation (1140) | Amino Acid Degradation (135) | Proteinogenic Amino Acid Degradation (115) | L-glutamate Degradation (11) |
| L-valine degradation I | Other microbial metabolism | Degradation/Utilization/Assimilation (1140) | Amino Acid Degradation (135) | Proteinogenic Amino Acid Degradation (115) | L-valine Degradation (3) |
| L-tryptophan degradation IX | Other microbial metabolism | Superpathways (384) | L-tryptophan degradation IX | | |
| L-tryptophan degradation IX | Other microbial metabolism | Degradation/Utilization/Assimilation (1140) | Amino Acid Degradation (135) | Proteinogenic Amino Acid Degradation (115) | L-tryptophan Degradation (13) |
| starch degradation III | Pyruvate generation | Degradation/Utilization/Assimilation (1140) | Polymeric Compound Degradation (50) | Polysaccharide Degradation (45) | Starch Degradation (5) |
| starch degradation III | Pyruvate generation | Degradation/Utilization/Assimilation (1140) | Carbohydrate Degradation (178) | Polysaccharide Degradation (45) | Starch Degradation (5) |
| L-lysine fermentation to acetate and butanoate | SCFA directly producing pathways | Generation of Precursor Metabolites and Energy (256) | Fermentation (84) | Fermentation to Short-Chain Fatty Acids (42) | Fermentation to Butanoate (8) |
| L-lysine fermentation to acetate and butanoate | SCFA directly producing pathways | Generation of Precursor Metabolites and Energy (256) | Fermentation (84) | Fermentation to Short-Chain Fatty Acids (42) | Fermentation to Acetate (23) |
| L-lysine fermentation to acetate and butanoate | SCFA directly producing pathways | Degradation/Utilization/Assimilation (1140) | Amino Acid Degradation (135) | Proteinogenic Amino Acid Degradation (115) | L-lysine Degradation (13) |
| GDP-D-glycero-alpha-D-manno-heptose biosynthesis | Cell surface & cell wall (host–microbe int | Biosynthesis (2132) | Carbohydrate Biosynthesis (236) | Sugar Biosynthesis (110) | Sugar Nucleotide Biosynthesis (94) |
| 4-hydroxyphenylacetate degradation | Other microbial metabolism | Degradation/Utilization/Assimilation (1140) | Aromatic Compound Degradation (230) | 4-hydroxyphenylacetate degradation | |
| trans,octa-cis decaprenyl phosphate biosynthesis | Cell surface & cell wall (host–microbe int | Biosynthesis (2132) | Polyprenyl Biosynthesis (18) | trans,octa-cis decaprenyl phosphate biosynthesis | |
| trans,octa-cis decaprenyl phosphate biosynthesis | Cell surface & cell wall (host–microbe int | Biosynthesis (2132) | Cofactor, Carrier, and Vitamin Biosynthesis (252) | Carrier Biosynthesis (131) | Lipid Anchor Biosynthesis (3) |
| superpathway of hexuronide and hexuronate degradation | Pyruvate generation | Superpathways (384) | superpathway of hexuronide and hexuronate degradation | | |
| superpathway of UDP-N-acetylglucosamine-derived O-antigen building blocks biosynthesis | Cell surface & cell wall (host–microbe int | Superpathways (384) | superpathway of UDP-N-acetylglucosamine-derived O-antigen building blocks biosynthesis | | |
| superpathway of UDP-N-acetylglucosamine-derived O-antigen building blocks biosynthesis | Cell surface & cell wall (host–microbe int | Biosynthesis (2132) | Carbohydrate Biosynthesis (236) | Sugar Biosynthesis (110) | Sugar Nucleotide Biosynthesis (94) |
| sitosterol degradation to androstenedione | Other microbial metabolism | Degradation/Utilization/Assimilation (1140) | Fatty Acid and Lipid Degradation (54) | Steroid Degradation (19) | sitosterol degradation to androstenedione |
| myo-, chiro- and scyllo-inositol degradation | Pyruvate generation | Superpathways (384) | myo-, chiro- and scyllo-inositol degradation | | |
| myo-, chiro- and scyllo-inositol degradation | Pyruvate generation | Degradation/Utilization/Assimilation (1140) | Cyclitol Degradation (13) | myo-, chiro- and scyllo-inositol degradation | |
| L-methionine salvage cycle III | Other microbial metabolism | Superpathways (384) | L-methionine salvage cycle III | | |
| L-methionine salvage cycle III | Other microbial metabolism | Biosynthesis (2132) | Amino Acid Biosynthesis (124) | Proteinogenic Amino Acid Biosynthesis (95) | L-methionine Biosynthesis (13) |
| S-methyl-5-thio-alpha-D-ribose 1-phosphate degradation I | Other microbial metabolism | Degradation/Utilization/Assimilation (1140) | Nucleoside and Nucleotide Degradation (41) | S-methyl-5-thio-alpha-D-ribose 1-phosphate Degradation (3) | S-methyl-5-thio-alpha-D-ribose 1-phosphate degradation I |
| S-methyl-5-thio-alpha-D-ribose 1-phosphate degradation I | Other microbial metabolism | Biosynthesis (2132) | Amino Acid Biosynthesis (124) | Proteinogenic Amino Acid Biosynthesis (95) | L-methionine Biosynthesis (13) |
| superpathway of chorismate metabolism | Other microbial metabolism | Superpathways (384) | superpathway of chorismate metabolism | | |
| pyridoxal 5'-phosphate biosynthesis I | Other microbial metabolism | Biosynthesis (2132) | Cofactor, Carrier, and Vitamin Biosynthesis (252) | Vitamin Biosynthesis (58) | Vitamin B6 Biosynthesis (5) |
| pyridoxal 5'-phosphate biosynthesis I | Other microbial metabolism | Biosynthesis (2132) | Cofactor, Carrier, and Vitamin Biosynthesis (252) | Enzyme Cofactor Biosynthesis (142) | Vitamin B6 Biosynthesis (5) |
| myo-inositol degradation I | Pyruvate generation | Degradation/Utilization/Assimilation (1140) | Cyclitol Degradation (13) | myo-inositol degradation I | |
| ketogluconate metabolism | Pyruvate generation | Superpathways (384) | ketogluconate metabolism | | |
| ketogluconate metabolism | Pyruvate generation | Degradation/Utilization/Assimilation (1140) | Carboxylic Acid Degradation (85) | Aldonic Acid Degradation (11) | ketogluconate metabolism |
| 3,8-divinyl-chlorophyllide a biosynthesis III (aerobic, light independent) | Other microbial metabolism | Biosynthesis (2132) | Tetrapyrrole Biosynthesis (39) | Porphyrin Compound Biosynthesis (39) | Chlorophyll Biosynthesis (15) |
| 3,8-divinyl-chlorophyllide a biosynthesis III (aerobic, light independent) | Other microbial metabolism | Biosynthesis (2132) | Secondary Metabolite Biosynthesis (765) | Pigment Biosynthesis (89) | Chlorophyll Biosynthesis (15) |
| 3,8-divinyl-chlorophyllide a biosynthesis II (anaerobic) | Other microbial metabolism | Biosynthesis (2132) | Tetrapyrrole Biosynthesis (39) | Porphyrin Compound Biosynthesis (39) | Chlorophyll Biosynthesis (15) |
| 3,8-divinyl-chlorophyllide a biosynthesis II (anaerobic) | Other microbial metabolism | Biosynthesis (2132) | Secondary Metabolite Biosynthesis (765) | Pigment Biosynthesis (89) | Chlorophyll Biosynthesis (15) |
| NAD de novo biosynthesis II (from tryptophan) | Other microbial metabolism | Superpathways (384) | NAD de novo biosynthesis II (from tryptophan) | | |
| NAD de novo biosynthesis II (from tryptophan) | Other microbial metabolism | Biosynthesis (2132) | Cofactor, Carrier, and Vitamin Biosynthesis (252) | Enzyme Cofactor Biosynthesis (142) | NAD Metabolism (22) |
| NAD de novo biosynthesis II (from tryptophan) | Other microbial metabolism | Biosynthesis (2132) | Cofactor, Carrier, and Vitamin Biosynthesis (252) | Enzyme Cofactor Biosynthesis (142) | NAD Biosynthesis (15) |
| NAD de novo biosynthesis II (from tryptophan) | Other microbial metabolism | Biosynthesis (2132) | Cofactor, Carrier, and Vitamin Biosynthesis (252) | Carrier Biosynthesis (131) | Electron Carrier Biosynthesis (92) |
| L-tryptophan degradation to 2-amino-3-carboxymuconate semialdehyde | Other microbial metabolism | Degradation/Utilization/Assimilation (1140) | Amino Acid Degradation (135) | Proteinogenic Amino Acid Degradation (115) | L-tryptophan Degradation (13) |
| acetyl-CoA fermentation to butanoate | SCFA directly producing pathways | Superpathways (384) | acetyl-CoA fermentation to butanoate | | |
| acetyl-CoA fermentation to butanoate | SCFA directly producing pathways | Generation of Precursor Metabolites and Energy (256) | Fermentation (84) | Fermentation to Short-Chain Fatty Acids (42) | Fermentation to Butanoate (8) |
| L-histidine degradation II | Other microbial metabolism | Degradation/Utilization/Assimilation (1140) | Amino Acid Degradation (135) | Proteinogenic Amino Acid Degradation (115) | L-histidine Degradation (6) |
| 4-deoxy-L-threo-hex-4-enopyranuronate degradation | Pyruvate generation | Degradation/Utilization/Assimilation (1140) | Carbohydrate Degradation (178) | Sugar Degradation (95) | Sugar Derivative Degradation (15) |
| 4-aminobutanoate degradation V | Other microbial metabolism | Generation of Precursor Metabolites and Energy (256) | Fermentation (84) | Fermentation to Short-Chain Fatty Acids (42) | Fermentation to Butanoate (8) |
| 4-aminobutanoate degradation V | Other microbial metabolism | Degradation/Utilization/Assimilation (1140) | Amide, Amidine, Amine, and Polyamine Degradation (68) | 4-Aminobutanoate Degradation (8) | 4-aminobutanoate degradation V |
| superpathway of geranylgeranyldiphosphate biosynthesis I (via mevalonate) | Other microbial metabolism | Superpathways (384) | superpathway of geranylgeranyldiphosphate biosynthesis I (via mevalonate) | | |
| superpathway of geranylgeranyldiphosphate biosynthesis I (via mevalonate) | Other microbial metabolism | Biosynthesis (2132) | Secondary Metabolite Biosynthesis (765) | Terpenoid Biosynthesis (208) | Diterpenoid Biosynthesis (44) |
| superpathway of geranylgeranyldiphosphate biosynthesis I (via mevalonate) | Other microbial metabolism | Biosynthesis (2132) | Polyprenyl Biosynthesis (18) | All-trans Polyprenyl Diphosphate Biosynthesis (15) | Geranylgeranyl Diphosphate Biosynthesis (3) |
| mevalonate pathway I (eukaryotes and bacteria) | Other microbial metabolism | Biosynthesis (2132) | Secondary Metabolite Biosynthesis (765) | Terpenoid Biosynthesis (208) | Hemiterpene Biosynthesis (10) |

### Supplementary Table 4

| MetaCyc_pathway | category | Lv2 | Lv3 | Lv4 | Lv5 |
|---|---|---|---|---|---|
| (S)-propane-1,2-diol degradation | SCFA directly producing pathways | Degradation/Utilization/Assimilation (1140) | Alcohol Degradation (41) | (S)-propane-1,2-diol degradation | |
| (S)-propane-1,2-diol degradation | SCFA directly producing pathways | Generation of Precursor Metabolites and Energy (256) | Fermentation (84) | Fermentation to Short-Chain Fatty Acids (42) | Fermentation to Propanoate (7) |
| reductive acetyl coenzyme A pathway I (homoacetogenic bact | Carbon allocation | Degradation/Utilization/Assimilation (1140) | C1 Compound Utilization and Assimilation (39) | CO2 Fixation (15) | Autotrophic CO2 Fixation (12) |
| sucrose biosynthesis I (from photosynthesis) | Carbon allocation | Biosynthesis (2132) | Carbohydrate Biosynthesis (236) | Sugar Biosynthesis (110) | Disaccharide Biosynthesis (11) |
| sucrose biosynthesis I (from photosynthesis) | Carbon allocation | Superpathways (384) | sucrose biosynthesis I (from photosynthesis) | | |
| peptidoglycan biosynthesis II (staphylococci) | Cell wall | Biosynthesis (2132) | Cell Structure Biosynthesis (123) | Cell Wall Biosynthesis (27) | Peptidoglycan Biosynthesis (6) |
| peptidoglycan biosynthesis II (staphylococci) | Cell wall | Superpathways (384) | peptidoglycan biosynthesis II (staphylococci) | | |
| adenine and adenosine salvage III | Nucleotide turnover | Biosynthesis (2132) | Nucleoside and Nucleotide Biosynthesis (57) | Purine Nucleotide Biosynthesis (33) | Purine Nucleotide Salvage (13) |
| sucrose biosynthesis III | Carbon allocation | Biosynthesis (2132) | Carbohydrate Biosynthesis (236) | Sugar Biosynthesis (110) | Disaccharide Biosynthesis (11) |
| superpathway of glyoxylate bypass and TCA | TCA & oxidative central metabolism | Superpathways (384) | superpathway of glyoxylate bypass and TCA | | |
| pyrimidine deoxyribonucleosides salvage | Nucleotide turnover | Biosynthesis (2132) | Nucleoside and Nucleotide Biosynthesis (57) | Pyrimidine Nucleotide Biosynthesis (23) | Pyrimidine Nucleotide Salvage (12) |
| TCA cycle VI (Helicobacter) | TCA & oxidative central metabolism | Generation of Precursor Metabolites and Energy (256) | TCA cycle (9) | TCA cycle VI (Helicobacter) | |
| purine ribonucleosides degradation | Nucleotide turnover | Degradation/Utilization/Assimilation (1140) | Nucleoside and Nucleotide Degradation (41) | Purine Nucleotide Degradation (16) | purine ribonucleosides degradation |
| superpathway of purine deoxyribonucleosides degradation | Nucleotide turnover | Degradation/Utilization/Assimilation (1140) | Nucleoside and Nucleotide Degradation (41) | superpathway of purine deoxyribonucleosides degradation | |
| superpathway of purine deoxyribonucleosides degradation | Nucleotide turnover | Superpathways (384) | superpathway of purine deoxyribonucleosides degradation | | |
| peptidoglycan maturation (meso-diaminopimelate containin | Cell wall | Biosynthesis (2132) | Cell Structure Biosynthesis (123) | Cell Wall Biosynthesis (27) | Peptidoglycan Biosynthesis (6) |
| pyruvate fermentation to acetate and lactate II | SCFA directly producing pathways | Generation of Precursor Metabolites and Energy (256) | Fermentation (84) | Fermentation of Pyruvate (36) | Pyruvate Fermentation to Acetate (11) |
| pyruvate fermentation to acetate and lactate II | SCFA directly producing pathways | Generation of Precursor Metabolites and Energy (256) | Fermentation (84) | Fermentation of Pyruvate (36) | Pyruvate Fermentation to Lactate (4) |
| pyruvate fermentation to acetate and lactate II | SCFA directly producing pathways | Generation of Precursor Metabolites and Energy (256) | Fermentation (84) | Fermentation to Short-Chain Fatty Acids (42) | Fermentation to Acetate (23) |
| pyruvate fermentation to acetate and lactate II | SCFA directly producing pathways | Generation of Precursor Metabolites and Energy (256) | Fermentation (84) | Fermentation to Short-Chain Fatty Acids (42) | Fermentation to Lactate (8) |
| pyruvate fermentation to acetate and lactate II | SCFA directly producing pathways | Superpathways (384) | pyruvate fermentation to acetate and lactate II | | |
| myo-, chiro- and scyllo-inositol degradation | Pyruvate generation | Degradation/Utilization/Assimilation (1140) | Cyclitol Degradation (13) | myo-, chiro- and scyllo-inositol degradation | |
| myo-, chiro- and scyllo-inositol degradation | Pyruvate generation | Superpathways (384) | myo-, chiro- and scyllo-inositol degradation | | |
| TCA cycle VII (acetate-producers) | SCFA directly producing pathways | Generation of Precursor Metabolites and Energy (256) | TCA cycle (9) | TCA cycle VII (acetate-producers) | |
| TCA cycle VII (acetate-producers) | TCA & oxidative central metabolism | Generation of Precursor Metabolites and Energy (256) | TCA cycle (9) | TCA cycle VII (acetate-producers) | |
| catechol degradation III (ortho-cleavage pathway) | Aromatic compound degradation | Degradation/Utilization/Assimilation (1140) | Aromatic Compound Degradation (230) | Catechol Degradation (7) | catechol degradation III (ortho-cleavage pathway) |
| catechol degradation III (ortho-cleavage pathway) | Aromatic compound degradation | Superpathways (384) | catechol degradation III (ortho-cleavage pathway) | | |
| aromatic compounds degradation via 3-oxoadipate | Aromatic compound degradation | Degradation/Utilization/Assimilation (1140) | Aromatic Compound Degradation (230) | Catechol Degradation (7) | aromatic compounds degradation via 3-oxoadipate |
| aromatic compounds degradation via 3-oxoadipate | Aromatic compound degradation | Superpathways (384) | aromatic compounds degradation via 3-oxoadipate | | |
| superpathway of pyrimidine deoxyribonucleosides degradati | Nucleotide turnover | Degradation/Utilization/Assimilation (1140) | Nucleoside and Nucleotide Degradation (41) | Pyrimidine Nucleotide Degradation (13) | superpathway of pyrimidine deoxyribonucleosides degradation |
| superpathway of pyrimidine deoxyribonucleosides degradati | Nucleotide turnover | Superpathways (384) | superpathway of pyrimidine deoxyribonucleosides degradation | | |
| acetylene degradation (anaerobic) | SCFA directly producing pathways | Generation of Precursor Metabolites and Energy (256) | Fermentation (84) | Fermentation of Pyruvate (36) | Pyruvate Fermentation to Ethanol (7) |
| acetylene degradation (anaerobic) | SCFA directly producing pathways | Generation of Precursor Metabolites and Energy (256) | Fermentation (84) | Fermentation to Alcohols (17) | Pyruvate Fermentation to Ethanol (7) |
| acetylene degradation (anaerobic) | SCFA directly producing pathways | Generation of Precursor Metabolites and Energy (256) | Fermentation (84) | Fermentation to Short-Chain Fatty Acids (42) | Fermentation to Acetate (23) |
| catechol degradation to beta-ketoadipate | Aromatic compound degradation | Degradation/Utilization/Assimilation (1140) | Aromatic Compound Degradation (230) | Catechol Degradation (7) | catechol degradation to beta-ketoadipate |
| peptidoglycan biosynthesis IV (Enterococcus faecium) | Cell wall | Biosynthesis (2132) | Cell Structure Biosynthesis (123) | Cell Wall Biosynthesis (27) | Peptidoglycan Biosynthesis (6) |
| peptidoglycan biosynthesis IV (Enterococcus faecium) | Cell wall | Detoxification (69) | Antibiotic Resistance (8) | peptidoglycan biosynthesis IV (Enterococcus faecium) | |
| peptidoglycan biosynthesis IV (Enterococcus faecium) | Cell wall | Superpathways (384) | peptidoglycan biosynthesis IV (Enterococcus faecium) | | |
| superpathway of N-acetylglucosamine, N-acetylmannosamine | Pyruvate generation | Degradation/Utilization/Assimilation (1140) | Amide, Amidine, Amine, and Polyamine Degradation (68) | superpathway of N-acetylglucosamine, N-acetylmannosamine and N-acetylneuraminate degradation | |
| superpathway of N-acetylglucosamine, N-acetylmannosamine | Pyruvate generation | Superpathways (384) | superpathway of N-acetylglucosamine, N-acetylmannosamine and N-acetylneuraminate degradation | | |
| glycolysis III (from glucose) | Pyruvate generation | Generation of Precursor Metabolites and Energy (256) | Glycolysis (6) | glycolysis III (from glucose) | |
| tRNA charging | Translation capacity | Biosynthesis (2132) | Aminoacyl-tRNA Charging (4) | tRNA charging | |
| tRNA charging | Translation capacity | Metabolic Clusters (70) | tRNA charging | | |
| UDP-N-acetylmuramoyl-pentapeptide biosynthesis I (meso-di | Cell wall | Biosynthesis (2132) | Cell Structure Biosynthesis (123) | Cell Wall Biosynthesis (27) | UDP-N-Acetylmuramoyl-Pentapeptide Biosynthesis (3) |
| peptidoglycan biosynthesis I (meso-diaminopimelate contain | Cell wall | Biosynthesis (2132) | Cell Structure Biosynthesis (123) | Cell Wall Biosynthesis (27) | Peptidoglycan Biosynthesis (6) |
| peptidoglycan biosynthesis I (meso-diaminopimelate contain | Cell wall | Superpathways (384) | peptidoglycan biosynthesis I (meso-diaminopimelate containing) | | |
| peptidoglycan biosynthesis III (mycobacteria) | Cell wall | Biosynthesis (2132) | Cell Structure Biosynthesis (123) | Cell Wall Biosynthesis (27) | Peptidoglycan Biosynthesis (6) |
| peptidoglycan biosynthesis III (mycobacteria) | Cell wall | Superpathways (384) | peptidoglycan biosynthesis III (mycobacteria) | | |
| UDP-N-acetylmuramoyl-pentapeptide biosynthesis II (lysine-c | Cell wall | Biosynthesis (2132) | Cell Structure Biosynthesis (123) | Cell Wall Biosynthesis (27) | UDP-N-Acetylmuramoyl-Pentapeptide Biosynthesis (3) |

#### Abundance of aromatic-degradation-associated genera (ASVs) in cave pool water

Units: abundance per million (sum of ASV counts / sample total x $10^6$)

n_ASV = number of ASVs collapsed into that row.

| Phylum | Family | Genus | n_ASV | Pachon cave Large_Pool _Site_1 | Large_Pool _Site_2 | Large_Pool _Site_3 | Small_Pool _Site_1 | Small_Pool _Site_2 | Small_Pool _Site_3 | Tinaja Large_Pool _Site_1 | Large_Pool _Site_2 | Large_Pool _Site_3 |
|---|---|---|---|---|---|---|---|---|---|---|---|---|
| Proteobacteria | Hyphomicrobiaceae | Pedomicrobium | 26 | 2890.6 | 2893.2 | 3294.8 | 2407.2 | 3621.0 | 3740.4 | 7886.4 | 5478.9 | 4426.3 |
| Proteobacteria | Pseudomonadaceae | Pseudomonas | 68 | 1473.5 | 3441.6 | 4402.8 | 5572.0 | 6225.3 | 8007.8 | 99.3 | 71.2 | 2536.8 |
| Proteobacteria | Chromobacteriaceae | Vogesella | 4 | 300.3 | 293.1 | 7209.2 | 1405.6 | 1980.0 | 1394.1 | 19.9 | 128.1 | 124.2 |
| Proteobacteria | Rhizobiales_Incertae_Sedis | Nordella | 9 | 610.0 | 737.5 | 772.7 | 1254.1 | 1783.7 | 1462.1 | 516.5 | 597.7 | 738.8 |
| Actinobacteriota | Pseudonocardiaceae | Pseudonocardia | 6 | 2421.4 | 567.3 | 590.4 | 673.3 | 1034.6 | 646.1 | 973.4 | 583.5 | 379.2 |
| Proteobacteria | Xanthobacteraceae | Pseudorhodoplanes | 4 | 225.2 | 245.8 | 379.0 | 227.3 | 909.7 | 748.1 | 774.7 | 740.0 | 601.5 |
| Proteobacteria | Xanthobacteraceae | Bradyrhizobium | 2 | 544.3 | 208.0 | 269.7 | 370.3 | 642.1 | 493.0 | 238.4 | 156.5 | 71.9 |
| Planctomycetota | Rubinisphaeraceae | sh_pl14 | 10 | 197.1 | 122.9 | 160.4 | 193.6 | 624.3 | 374.0 | 298.0 | 270.4 | 353.1 |
| Planctomycetota | Gemmataceae | Gemmata | 18 | 197.1 | 132.4 | 269.7 | 286.2 | 321.1 | 204.0 | 278.1 | 56.9 | 313.8 |
| Proteobacteria | Comamonadaceae | Rubrivivax | 1 | 150.2 | 122.9 | 539.4 | 218.8 | 338.9 | 663.1 | 59.6 | 14.2 | 39.2 |
| Proteobacteria | Rhodobacteraceae | Pseudorhodobacte | 7 | 0.0 | 37.8 | 189.5 | 606.0 | 410.3 | 102.0 | 0.0 | 42.7 | 13.1 |
| Proteobacteria | Oxalobacteraceae | Pseudoduganella | 1 | 46.9 | 132.4 | 357.2 | 50.5 | 89.2 | 306.0 | 0.0 | 14.2 | 98.1 |
| Proteobacteria | Oxalobacteraceae | Undibacterium | 5 | 9.4 | 18.9 | 466.5 | 75.8 | 285.4 | 306.0 | 0.0 | 0.0 | 0.0 |
| Firmicutes | Bacillaceae | Anoxybacillus | 4 | 159.5 | 85.1 | 65.6 | 193.6 | 17.8 | 68.0 | 238.4 | 213.5 | 91.5 |
| | | **Total read count** | | 106,552 | 105,764 | 137,186 | 118,809 | 56,062 | 58,818 | 50,340 | 70,269 | 152,949 |

#### Abundance of the metabolic pathways positively associated with Pachón body standard length in cave pool

Units: abundance per million (pathway abundance / sample total x $10^6$)

| pathway | description | | Pachon cave Large_Pool _Site_1 | Large_Pool _Site_2 | Large_Pool _Site_3 | Small_Pool _Site_1 | Small_Pool _Site_2 | Small_Pool _Site_3 | Tinaja Large_Pool _Site_1 | Large_Pool _Site_2 | Large_Pool _Site_3 |
|---|---|---|---|---|---|---|---|---|---|---|---|
| **PWY-5431** | aromatic compounds degrad | | 144.5 | 153.0 | 199.4 | 214.6 | 228.8 | 241.6 | 104.5 | 101.4 | 128.9 |
| **CATECHOL-ORTHO-CLEAVA** | catechol degradation to β-ke | | 175.1 | 175.5 | 204.2 | 237.1 | 250.5 | 244.0 | 110.7 | 101.2 | 139.2 |
| **PWY-7237** | myo-, chiro- and scillo-inositol | | 195.2 | 205.4 | 227.8 | 273.9 | 313.7 | 267.8 | 208.7 | 172.4 | 180.1 |

Note: Cells are shaded with Red color scale
